# Mapping CAR T-cell design space using agent-based models

**DOI:** 10.1101/2022.04.07.487561

**Authors:** Alexis N. Prybutok, Jessica S. Yu, Joshua N. Leonard, Neda Bagheri

## Abstract

Chimeric antigen receptor (CAR) T-cell therapy shows promise for treating liquid cancers and increasingly for solid tumors as well. While potential design strategies exist to address translational challenges, including the lack of unique tumor antigens and the presence of an immunosuppressive tumor microenvironment, testing all possible design choices *in vitro* and *in vivo* is prohibitively expensive, time consuming, and laborious. To address this gap, we extended the modeling framework ARCADE (Agent-based Representation of Cells And Dynamic Environments) to include CAR T-cell agents (CAR T-cell ARCADE, or CARCADE). We conducted *in silico* experiments to investigate how clinically relevant design choices and inherent tumor features—CAR T-cell dose, CD4^+^:CD8^+^ CAR T-cell ratio, CAR-antigen affinity, cancer and healthy cell antigen expression—individually and collectively impact treatment outcomes. Our analysis revealed that tuning CAR affinity modulates IL-2 production by balancing CAR T-cell proliferation and effector function. It also identified a novel multi-feature tuned treatment strategy for balancing selectivity and efficacy and provided insights into how spatial effects can impact relative treatment performance in different contexts. CARCADE facilitates deeper biological understanding of treatment design and could ultimately enable identification of promising treatment strategies to accelerate solid tumor CAR T-cell design-build-test cycles.

## 1 Introduction

Chimeric antigen receptor (CAR) T-cell therapy combines advances in cellular engineering and personalized medicine for patient-specific, targeted cancer treatment (Barrett et al., 2014; Jackson et al., 2016). This therapy involves collecting, purifying, and genetically modifying a patient’s own T-cells to express a CAR that specifically targets the patient’s tumor(s) (Barrett et al., 2014; Jackson et al., 2016). These engineered cells are expanded *ex vivo* and then re-infused into the patient where the CAR T-cells target and kill antigen-expressing tumor cells. The two FDA-approved CAR T-cell therapies and many studies expanding CAR designs exclusively target “liquid” cancers derived from CD19^+^ B-cells (Jackson et al., 2016; Castellarin et al., 2018; Yanez-Munoz and Grupp, 2018). CD19 CAR T-cell therapies have shown great success in the clinic with response rates between 70-90% reported (Lim and June, 2017). In contrast, response rates for solid cancers are significantly lower at 4-16% (Hou et al., 2019).

CAR T-cells are currently less effective for treating solid tumors due unique complexities of both the tumor microenvironment (TME) and tumors themselves. First, TME barriers prevent CAR T-cell infiltration (Castellarin et al., 2018). These barriers include the intricate influence of both tumor-suppressing and tumor-promoting cells on the TME (Whiteside, 2008; Galluzzi et al., 2018), immune-evading cell markers promoting tumor escape (Maus and June, 2016; Galluzzi et al., 2018), and physical and chemical barriers that impact spatial dynamics and nutrient availability (Whiteside, 2008; Castellarin et al., 2018). Thus, developing CAR T-cells that remodel the immunosuppressive TME has been an active area of research (Cherkassky et al., 2016; Liu et al., 2016; Lim and June, 2017; Huang et al., 2018; Rafiq et al., 2018). Second, solid tumors often lack unique tumor antigens for selective targeting (Kakarla and Gottschalk, 2014). Cross-reactivity with healthy tissues present harmful or fatal off-tumor effects (Bonifant et al., 2016; Lim and June, 2017). Cellular engineering efforts have focused on increasing CAR specificity by tuning the affinity of receptor-antigen interactions to avoid healthy cells (Caruso et al., 2015; Johnson et al., 2015; Liu et al., 2015; Castellarin et al., 2018). Similarly, creating CAR T-cells that perform Boolean logic can enhance tumor recognition specificity (Wilkie et al., 2012; Lanitis et al., 2013; Wu et al., 2015; Castellarin et al., 2018; Cho et al., 2018). Designing CAR T-cells that target multiple antigens simultaneously can also prevent formation of antigen escape variant tumors (Hegde et al., 2013; Hegde et al., 2016; Cho et al., 2018). Finally, additional factors that have not proven problematic for “liquid” cancers, such as the need for site-specific trafficking of CAR T-cells to solid tumors and tumor antigen heterogeneity, further complicate solid-tumor CAR T-cell therapy design (Lim and June, 2017).

In combination with the array of engineering design choices presented by addressing the constraints above, additional design choices impact CAR T-cell effector functions and long-term persistence regardless of tumor type. These features include CD4^+^:CD8^+^ CAR T-cell ratios (Zhao et al., 2015; Sommermeyer et al., 2016; Turtle et al., 2016), choice of intracellular co-stimulatory domain (ICD) in the CAR (Kawalekar et al., 2016; Guedan et al., 2018), and the stage of T-cell differentiation (Sommermeyer et al., 2016). Collectively, the vast number of design choices complicates interpreting and comparing studies of and iteratively tuning CAR T-cell therapies.

Simultaneously tuning multiple features of a CAR T-cell therapy and forecasting their impact on emergent population dynamics remains a grand challenge. Exploring the multidimensional design space becomes prohibitively expensive and laborious *in vitro* and *in vivo*, particularly when considering the time and resources required for mouse experiments. Additionally, some design aspects and emergent properties are difficult to interrogate experimentally, such as cell-level behavioral states that impact treatment efficacy. Employing *in silico* experiments has proven to be a resource-saving and valuable way to understand how underlying biological processes impact CAR treatment outcome and hypothesizing new design features to improve efficacy. Recent CAR T-cell modeling efforts have used ordinary differential equation (ODE) models to understand factors influencing CAR T-cell receptor signaling and downstream activation (Rohrs et al., 2018; Cess and Finley, 2020a; Rohrs et al., 2020). Other CAR T-cell ODE modeling efforts aim to optimize patient pre-conditioning with chemotherapy (Owens and Bozic, 2021). However, these models lack spatial resolution, test a limited set of features, and do not assess emergent cell population dynamics; these important contributions do not yet enable predictions of the sort needed to guide the design of CAR T-cell therapies.

Agent-based models (ABMs) provide ideal *in silico* testbeds for interrogating emergent population dynamics. ABMs are bottom-up computational frameworks that describe the behavior of autonomous agents through defined rules that guide agent actions and interactions within their local environment. The ABM framework provides single-cell spatial and temporal resolution, incorporates quantitative and qualitative experimental observations, and enables tuning and measuring properties of interest through *in silico* experiments (Chavali et al., 2008; Narang et al., 2012; Yu and Bagheri, 2016; Vodovotz et al., 2017). Past ABMs have explored how cell properties influence tumor growth (Zhang et al., 2009; Waclaw et al., 2015; Norton et al., 2017; Yu and Bagheri, 2020), vasculature and microenvironment dynamics (Anderson et al., 2006; Yu and Bagheri, 2021), immune response to infection and tumors (Folcik et al., 2007; Cess and Finley, 2020b), and tumor response to checkpoint inhibitor therapy (Gong et al., 2017). However, to our knowledge, no ABM reported to date has characterized CAR T-cell dynamics in solid tumors, or explored how CAR T-cell and tumor features impact outcomes.

In this study, we systematically explore CAR T-cell therapy designs in solid tumor contexts by adding CAR T-cell agents to an established ABM (Agent-based Representation of Cells And Dynamic Environments, or ARCADE) comprising tissue cell agents (Yu and Bagheri, 2020) and dynamic vasculature (Yu and Bagheri, 2021). We use this model—CAR T-cell ARCADE (CARCADE)—to simulate CAR T-cell interactions with tissue cells and analyze a multidimensional design space. We demonstrate that CARCADE recapitulates known observations and predicts responses to new designs for solid tumor CAR T-cell therapies.

## 2 Results

### 2.1 CARCADE characterizes CAR T-cell behavior, metabolism, and effector function

CARCADE provides a flexible framework for characterizing and exploring hypothesized dynamics of population-level tumor responses to CAR T-cell treatment by defining individual CAR T-cell, cancer, and healthy cell features and rules.

#### 2.1.1 CAR T-cell agents recapitulate CAR T-cell behavior

ARCADE comprises tissue cell agents with individual subcellular metabolism and signaling modules that influence the cell-level decision making rules and drive emergent population-and environment-level dynamics (**Figure 1A**). Tissue cell agent rules and parameters can be tuned to represent either cancer or healthy cells. We introduce a new cell agent representing CAR T-cells into this framework (**Figure 1A**). All cell agents are simulated in a microenvironment that comprises either constant nutrient sources (representing a dish context) or vasculature (representing a vascularized tissue context). To distinguish between simulation and experiment, we denote simulated dish and tissue contexts as `dish` and `tissue`, respectively.

**FIGURE 1.**
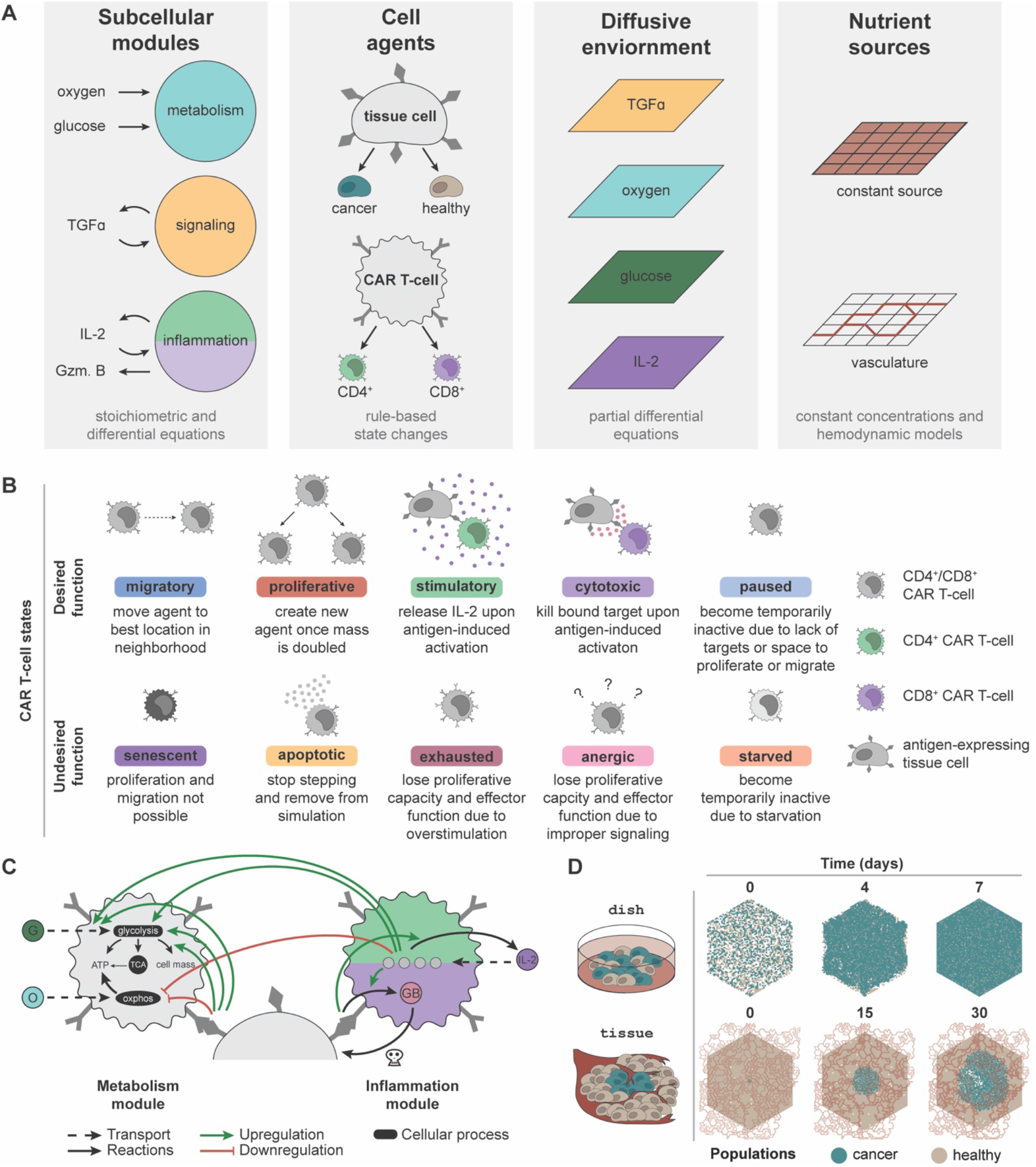
CARCADE structure and CAR T-cell agent design. **(A) Depiction of CARCADE components.** Subcellular modules guide underlying cellular function to influence behavior (Gzm. B: granzyme B). Agents include tissue cell and CAR T-cell agents, each of which has separate rule sets and is depicted with surface ligands and CARs (dark gray). Tissue cells include both healthy cells and cancer cells. Agents exist in an environment where diffusion is controlled by partial differential equations and constant sources or vasculature provide nutrients. (B) Descriptions of each CAR T-cell agent state, separated by whether the state is desired or undesired for efficacious treatment. (C) Diagram of CAR T-cell metabolism and inflammation module interactions with small molecules, proteins, and regulatory edges. The inflammation module diagram is broken into two parts, showing differences between CD4^+^ CAR T-cells (light green, top) and CD8^+^ CAR T-cells (purple, bottom). All CAR T-cells use identical metabolism modules. Regulatory edges (upregulation: green arrow, downregulation: red flathead arrow) result from IL-2 binding and antigen-induced activation. G: glucose, O: oxygen, GB: granzyme B., OXPHOS: oxidative phosphorylation. Legend for cell color is consistent with panel B. (D) An example of a single `dish` and `tissue` simulation of untreated cancer cells shown at select time points. For `tissue`, the dynamic vasculature architecture is overlaid.

Agents navigate through a set of defined, cell-type specific states and rules derived from experimentally observed states and transitions. Each tissue cell can be in one of six states—migratory, proliferative, quiescent, senescent, necrotic, and apoptotic—at each time step. CAR T-cell agents follow a unique rule set with additional states designed to capture T-cell behaviors (**Figure 1B**). There are two subtypes of CAR T-cell agents: CD8^+^ T-cells that primarily provide cytotoxic functions and CD4^+^ T-cells that primarily provide stimulatory functions (Liadi et al., 2015; Golubovskaya and Wu, 2016; Sommermeyer et al., 2016). Although both T-cell subtypes can provide cytotoxic and stimulatory functions, for simplicity, we specified that each of these T-cell subtypes would perform only their primary function. CAR T-cell agents can enter ten different subtype-dependent states, broadly categorized as desirable and undesirable during treatment. Desired states include migratory, proliferative, stimulatory (CD4^+^ only), cytotoxic (CD8^+^ only), and paused. Undesired states include apoptotic, senescent, exhausted, anergic, and starved. Cells change state according to the rule set and to their current state (**Supplementary Figures 1** and **2, Supplementary Methods Details**). All new model parameters are listed in **Supplementary Table 1** (Kuse et al., 1985; Lauffenburger and Linderman, 1993; Robertson et al., 1996; Frauwirth et al., 2002; De Boer et al., 2003; Deenick et al., 2003; Iwashima, 2003; Jacobs et al., 2008; Busse et al., 2010; Yoon et al., 2010; Wang et al., 2011; Altman and Dang, 2012; Robertson-Tessi et al., 2012; Stone et al., 2012; Cheng et al., 2013; Hegde et al., 2013; Heskamp et al., 2015; Kinjyo et al., 2015; Liu et al., 2015; Obst, 2015; Harris and Kranz, 2016; Hegde et al., 2016; Arcangeli et al., 2017; Borghans and Ribeiro, 2017; Gong et al., 2017; Gherbi et al., 2018; Guedan et al., 2018; Salter et al., 2018; Yu and Bagheri, 2020; 2021).

Each agent utilizes subcellular modules to capture underlying metabolic and signaling states. ARCADE tissue agents use two subcellular modules that control metabolism and signaling. The metabolism module uses stoichiometric equations to determine cellular uptake of glucose and oxygen, which is then converted to energy and cell mass. The signaling module uses an ODE model with regulatory nodes to determine the influence of tumor growth factor alpha (TGFα) on a tissue cell’s decision to proliferate or migrate. CAR T-cell agents use the tissue cell metabolism module with modifications to capture the influence of IL-2 signaling and antigen-induced activation on T-cell metabolism: (i) increased metabolic preference for glycolysis; (ii) increased glucose uptake rate; And (iii) increased fraction of glucose used to produce cell mass (**Figure 1C, Supplementary Methods Details**) (Frauwirth et al., 2002; Jones and Thompson, 2007; Pearce, 2010; Altman and Dang, 2012; Gerriets and Rathmell, 2012; MacIver et al., 2013; Chang and Pearce, 2016; Mehta et al., 2017). CAR T-cell agents also contain an inflammation module to capture the impact of IL-2 binding and antigen-induced activation on IL-2 production in CD4^+^ CAR T-cells (Malek and Castro, 2010; Liao et al., 2013; Rosenberg, 2014) and on granzyme production in CD8^+^ CAR T-cells (Liadi et al., 2015) (**Figure 1C, Supplementary Methods Details**). For both CAR T-cell subtypes, the inflammation module uses an ODE model to determine the amount of IL-2 bound to various IL-2 receptor species (Malek and Castro, 2010; Liao et al., 2013; Ross and Cantrell, 2018).

#### 2.1.2 *In silico* experiments mimic *in vitro* and *in vivo* contexts

To provide an *in silico* testbed that can be related to physical experiments, simulations were designed to represent two experimental contexts: `dish` and `tissue` (**Figure 1D**). Each configuration utilizes an environment in which four nutrient and signaling molecules—oxygen, glucose, TGFα, and IL— diffuse. Additionally, the environment contains distinct sources from which oxygen and glucose are produced. `Dish` uses a constant nutrient source environment to represent the well-mixed cell media of an *in vitro* experiment. These simulations are initialized with a defined number of tissue cells placed randomly in the environment. CAR T-cells are introduced after 10 min and simulated for 7 d of treatment. `Tissue` uses vasculature to represent realistic hemodynamics of nutrients diffusing through the environment to represent an *in vivo* solid tumor experiment. Vasculature can be degraded and collapse due to cancer cell crowding and movement. These simulations are initialized with a confluent bed of healthy cells and a small colony of cancer cells added to the center of the simulation environment. The cancer cell colony grows for 21 d to form a tumor before CAR T-cells are added and simulated for 9 d of treatment. Untreated `dish` and `tissue` simulations highlight how *in silico* experimental design leads to diverse outcomes (**Figure 1D**).

### 2.2 Monoculture and co-culture simulations are consistent with *in vitro* observations

CAR T-cell agents were developed *de novo* based on established cell-level observations; resulting emergent dynamics of the simulation were used for model validation. The comparison between *in silico* and *in vitro/in vivo* experiments is a critical and common method for validating ABMs. To confirm that emergent dynamics follow experimental observations, we tested how outcomes vary as a function of four CAR and tumor features—CAR T-cell dose (Sampson et al., 2014), CD4^+^:CD8^+^ CAR T-cell ratio (Sommermeyer et al., 2016; Turtle et al., 2016), CAR-antigen affinity (Chmielewski et al., 2004; Hudecek et al., 2013; Caruso et al., 2015; Johnson et al., 2015; Liu et al., 2015; Ghorashian et al., 2019), and antigen density on cancer cells (Stone et al., 2012; Liu et al., 2015; Watanabe et al., 2015; Majzner et al., 2020).

In a clinical setting, CAR T-cells necessarily interact with both healthy and cancer cells, and healthy cell antigen expression can impact off-target effects (Harris and Kranz, 2016). It is critical to consider how these CAR and tumor features impact both cancer and healthy cell populations. We simulated CAR T-cell treatment in three different contexts—(i) monoculture with only cancer cells, (ii) ideal co-culture with cancer cells and antigen-negative healthy cells, and (iii) realistic co-culture with cancer cells and low-level antigen expressing healthy cells—modulating CAR T cells and tumor features in each context to assess how *in silico* dynamics compare to observations *in vitro*. Using `dish` removes confounding effects of nutrient constraints and TME factors. We simulated 10 replicates of each combination of features (**Supplementary Table 2** for monoculture, and **Supplementary Table 3** for co-culture). In monoculture, `dish` was randomly plated at t = 0 s with 2 × 10^3^ antigen-expressing cancer cells. At t = 10 min, treatment begins by adding a dose of CAR T-cells, each expressing 5 × 10^4^ CARs with a defined CAR affinity and CD4^+^:CD8^+^ ratio. We simulated 7 d of treatment. Co-culture is identical except initial plating uses 1 × 10^3^ cancer cells and 1 × 10^3^ healthy cells. Simulation trajectories—including each cell’s location, state, volume, and average cell cycle length—were collected every half day. The input files used to generate `dish` simulations are described in the **Supplementary Material** (**Supplementary Data 1** and **Supplementary Table 4** for monoculture and **Supplementary Data 2** and **Supplementary Table 5** for co-culture).

#### 2.2.1 Cancer cell and CAR T-cell dynamics are independent of context

We first consider the impact of individual features on cell counts and behavior in `dish` (holding other features constant at intermediate values). In all simulations, cancer cell and CAR T-cell counts follow experimentally observed trends, including conditions with effector-to-target (E:T) ratios less than one where cancer cell killing occurs over several days (**Figure 2A** for monoculture, **Supplementary Figure 3A** for ideal co-culture, **Supplementary Figure 3B** for realistic co-culture) (Chmielewski et al., 2004; Arcangeli et al., 2017). Increasing CAR T-cell dose increases T-cell counts and accelerates cancer cell killing (Sampson et al., 2014; Hamieh et al., 2019). Our simulations mirror this trend; when E:T ratios are increased beyond the initial range explored (i.e., to explore ratios greater than one), substantial cancer cell killing occurred in monoculture in half the time (all other features are held at intermediate values) (**Supplementary Data 3** and **Supplementary Figure 4A**). Intermediate CD4^+^:CD8^+^ ratios maximize cancer killing and increase CAR T-cell proliferation (Sommermeyer et al., 2016; Turtle et al., 2016). Higher fractions of CD8^+^ CAR T-cell treatments prove less effective because cytotoxic CD8^+^ cells need the support of the cytokines primarily produced by CD4^+^ cells (Liadi et al., 2015; Golubovskaya and Wu, 2016). We tested an expanded range of CD4^+^:CD8^+^ ratios to include 90:10 and 10:90 in monoculture and co-culture; these extensions further validated observed trends and provided no additional treatment benefit, and thus we do not carry these conditions forward in subsequent analyses (see **Supplementary Note 1, Supplementary Data 4**, and **Supplementary Figure 5**). Increasing CAR affinity increases the chances of CAR T-cell antigen binding and subsequent activation, resulting in increased cancer cell killing (Chmielewski et al., 2004; Liu et al., 2015; Hernandez-Lopez et al., 2021). This increased activation also leads to increased proliferation and thus increased T-cell count (Caruso et al., 2015). Increased antigen expression on cancer cells increases cancer cell killing (Chmielewski et al., 2004; Liu et al., 2015; Watanabe et al., 2015; Arcangeli et al., 2017). Similarly, because CAR T-cells are more likely to be activated by high antigen density cancer cells, CAR T-cell proliferation, and thus counts, increase with increasing antigen count (Hernandez-Lopez et al., 2021). CD8^+^ T-cells counts exceed CD4^+^ T-cell counts even when cells are delivered at a 50:50 ratio, especially in conditions where cells are more likely to be activated (Sommermeyer et al., 2016; Turtle et al., 2016). The lowest CAR T-cell counts occur when we treat with only one subset of CAR T-cells. Cancer cells cannot be killed off without CD8^+^ cells. CD8^+^ cells have limited killing and proliferative capacity without cytokines produced by CD4^+^ cells, and lack of cancer cell killing presents spatial limitations on CAR T-cell proliferation. Overall, all `dish` simulations, regardless of healthy cell context, support experimental observations of cancer and CAR T-cell dynamics, suggesting that healthy cell presence and antigen expression do not strongly influence cancer and CAR T-cell dynamics or individual feature trends *in vitro*.

**FIGURE 2.**
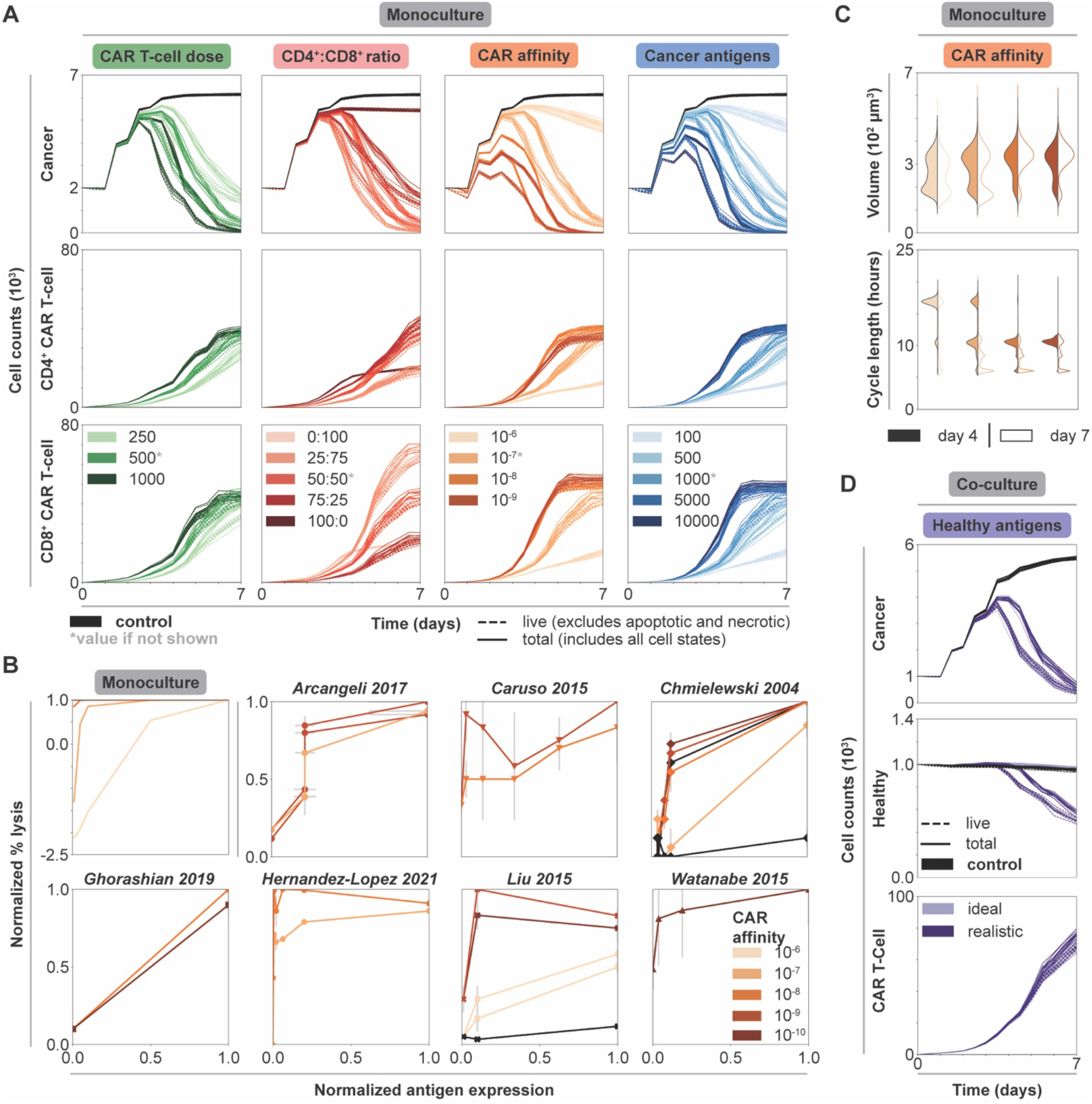
Impact of individual CAR T-cell and tumor features on cytotoxicity and CAR T-cell growth in dish. **(A)** Cell counts over time of untreated (black) and treated conditions (graded hues) holding all but one feature constant. Each column shows the axis being changed, where all other features are held constant at indicated intermediate values (indicated by asterisk, CAR T-cell dose–500 CAR T-cells, CD4^+^:CD8^+^ ratio–50:50, CAR affinity–10^−7^ M, cancer antigens–1000 antigens/cell), while rows show the cell type being plotted. **(B)** Normalized percent lysis curves for *in silico* and published experimental *in vitro* data. Plot for simulated data shows percent lysis for each set of CAR affinity values across normalized cancer antigen values. All other axes were held constant, and the data were averaged across replicates. Simulations with negative percent lysis indicate cancer cell growth. Experimental data—representing an array of CAR types, effector to target (E:T) ratios, ICDs, and cancer cell lines (**Supplementary Table 6** and **Supplementary Data 5**)—were normalized to maximum percent lysis and antigen levels with estimated error bars. The plots show percent lysis for each set of CARs tested per paper, each with unique CAR affinity and tested across a range of antigen target values. **(C)** Volume and cell cycle distributions for CAR T-cell populations at t = 4 d (filled) and t = 7 d (outline) holding all but CAR affinity constant at an intermediate value in monoculture. Legend is consistent with panel B. The data for cancer cell populations and for all other features can be found in the in supplement. **(D)** Cell counts over time of untreated (black) and treated conditions (graded hues) holding all features constant at an intermediate value. Legend is consistent with panel B for both ideal and realistic co-culture. Solid lines represent total cell counts, dashed lines represent live cell counts.

#### 2.2.2 Monoculture data qualitatively recapitulate a range of *in vitro* CAR T-cell studies

Quantifying percent lysis as a function of cancer antigen density is a common experimental analysis. In monoculture, percent lysis increases as a function of both antigen count and CAR affinity. This qualitative trend and the general shape of the data agrees with prior *in vitro* observations (**Figure 2B**) (Chmielewski et al., 2004; Caruso et al., 2015; Liu et al., 2015; Watanabe et al., 2015; Arcangeli et al., 2017; Ghorashian et al., 2019; Hernandez-Lopez et al., 2021). Additionally, for monoculture and most *in vitro* data, higher CAR affinities promote higher percent lysis across all antigen expression values. Our simulations reproduce general trends observed across diverse *in vitro* studies varying in CAR, intracellular co-stimulatory domain, effector to target ratio, and cell lines (**Supplementary Table 6** and **Supplementary Data 5**). Notably, CARCADE captures known experimental trends relevant to many different experimental CAR T-cell scenarios without being trained to any specific CAR T-cell experiment. Consistency in these emergent dynamics provides baseline validation that supports our use of the model to interrogate CAR T-cell design.

#### 2.2.3 Trends in cell-level features support population-level observations and model validation

Treatment efficacy can be evaluated by volume (Jacobs et al., 2008) and cell cycle length (Yoon et al., 2010) distributions, which serve as proxies for CAR T-cell growth and proliferation resulting from antigen-induced activation and IL-2 binding. As an increasing number of CAR T-cells undergo antigen-induced activation, CAR T-cell volumes increase and cycle lengths decrease both over time and with increasing CAR affinity (**Figure 2C**). In T-cells, antigen-induced activation and IL-2 binding influence metabolism to help T-cells rapidly proliferate by increasing nutrient uptake, metabolic preference for glucose, and flux of nutrients towards producing cell mass (Frauwirth et al., 2002; Jacobs et al., 2008; Pearce, 2010; Altman and Dang, 2012; Gerriets and Rathmell, 2012; Buck et al., 2015; Chang and Pearce, 2016; Mehta et al., 2017). These internal cellular changes increase cell growth rates, increase volumes, and decrease cell cycle lengths (van Stipdonk et al., 2003; Jacobs et al., 2008; Yoon et al., 2010; Altman and Dang, 2012; Kinjyo et al., 2015). The cell cycle length observed *in silico*—an emergent property of the simulations—ranged from around 6-24 h and falls within the range of 2-24 h found *in vitro, in vivo*, and for other *in silico* models (De Boer et al., 2003; van Stipdonk et al., 2003; Yoon et al., 2010; Altman and Dang, 2012; Kinjyo et al., 2015; Gong et al., 2017). Cancer cell volumes increase slightly and cycle lengths decrease slightly with increasing CAR affinity and over time, as cancer cells proliferate to compensate for cell death (**Supplementary Figure 6A**). Similar trends in volume and cell length distributions are observed across all other modulated features, where conditions with more activated CAR T-cells result in increased CAR T-cell volume and decreased cell cycle lengths (**Supplementary Figure 6B-D**). Altogether, the model recapitulates known *in vitro* observations, and furthermore, it enables us to observe single cell-level properties that are non-trivial to measure experimentally.

### 2.3 Varying individual features highlights tradeoffs within co-culture

Due to the lack of unique tumor antigens, CAR T-cell designs must rely on target antigens that are more highly expressed on cancer cells than healthy cells (Harris and Kranz, 2016). Investigating the difference in treatment outcomes—cancer cell killing, healthy cell sparing, and CAR T-cell growth— between the ideal co-culture (containing antigen-negative healthy cells) and realistic co-culture (containing antigen-expressing healthy cells) is critical for understanding successful CAR T-cell design (Caruso et al., 2015).

#### 2.3.1 Healthy cell antigen expression and tumor/CAR T-cell features impact healthy cell killing

Healthy cell antigen density does not affect cancer cell killing, CAR T-cell proliferation, or previously noted trends across individual features for these populations (**Supplementary Figure 3**). However, healthy cell antigen density dramatically impacts healthy cell killing (**Figure 2D**) (Arcangeli et al., 2017). The seeming lack of influence that minimal healthy antigen expression has on CAR T-cell proliferation is demonstrated by a lack of clear difference in CAR T-cell volume and cell cycle length distributions (**Supplementary Figure 7**) or fraction of cells in the proliferative state (**Supplementary Figure 8**) between the ideal and realistic co-culture. In general, we hypothesize that the low healthy cell antigen level is too weak to impact these other factors but enables the CAR T-cells to target healthy cells. Thus, healthy cell antigen expression only needs to be considered in avoiding healthy cell death and not in tuning CAR T-cell behavior or cancer cell killing.

To further investigate the impact of healthy cell antigen expression on feature trends, we directly compare cell counts between the ideal and realistic co-culture along the CAR affinity feature axis (**Figure 3A**). Cancer cell killing dynamics are nearly identical in both contexts, increasing with increased CAR affinity. In contrast, healthy cell dynamics differ dramatically between contexts. When healthy cells do not express antigen, increasing CAR affinity leads to increased healthy cell count as healthy cells grow to fill the space left behind by targeted cancer cells. However, when healthy cells do express antigen, healthy cell killing increases with increasing CAR affinity. Additionally, healthy cell counts begin to decrease at increasingly earlier time points with increasing CAR affinity. Comparing cell counts along other features exhibits similar trends: presence of healthy cell antigen generally only impacts healthy cell dynamics, resulting in varying degrees of healthy cell killing (**Supplementary Figure 3**). These data are consistent with experimental studies demonstrating a detrimental effect of high CAR affinity designs on healthy cells (Caruso et al., 2015; Harris and Kranz, 2016). Low affinity CARs successfully target tumors that overexpress the desired antigen and produce minimal off-tumor effects when healthy cells express low antigen levels (Caruso et al., 2015; Johnson et al., 2015; Liu et al., 2015). When healthy cells express antigen, it is not always desirable to have the strongest affinity CAR T-cells.

**FIGURE 3.**
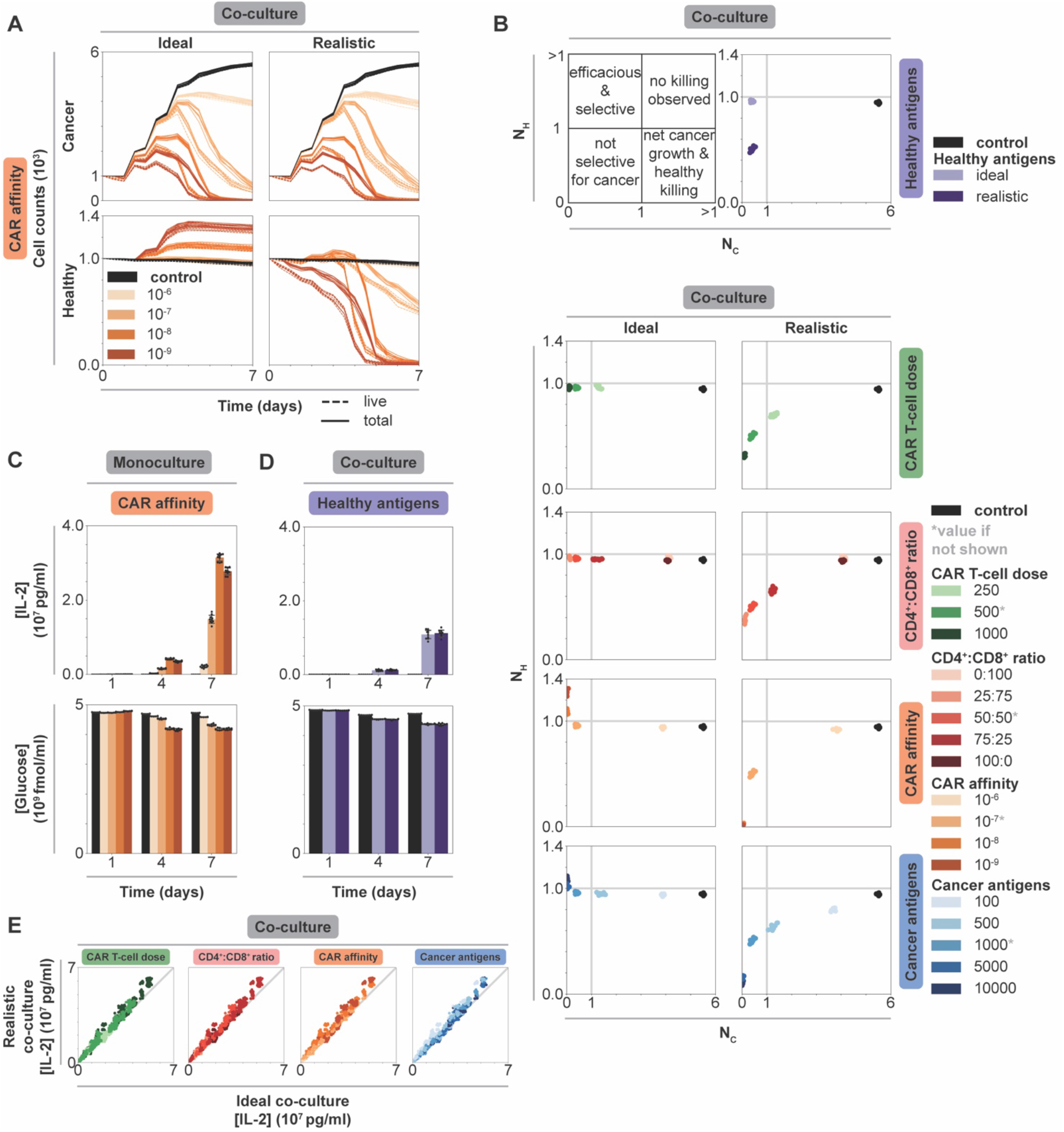
Impact of individual CAR T-cell and tumor features on efficacy, selectivity, and cytokine production in monoculture vs co-culture. **(A)** Cancer and healthy cell counts over time of untreated (black) and treated (graded hues) conditions holding all but CAR affinity, which is reported in units of M, constant at an intermediate value and separating data by co-culture context. Column shows co-culture type, row shows cell type. Solid lines represent total cell counts, dashed lines represent live cell counts (live, excludes necrotic and apoptotic states as in **Figure 2**). Intermediate values of other features indicated by asterisk in panel B: CAR T-cell dose–500 CAR T-cells, CD4^+^:CD8^+^ ratio–50:50, cancer antigens–1000 antigens/cell. **(B)** Scatter plots of normalized live healthy cell count (*N*_*H*_) vs normalized live cancer cell count (*N*_*C*_) for untreated (black) and treated conditions (graded hues) holding all but one axis constant at an intermediate value. Upper left plot shows quadrant meanings. Upper right plot shows scatter plot for different co-culture contexts. Columns show co-culture type, and each row indicates which feature is being plotted. **(C)** IL-2 and glucose concentrations over time holding all but CAR affinity constant at an intermediate value in monoculture. Legend is consistent with panel B. **(D)** IL-2 and glucose concentrations over time varying CAR affinity while holding all features constant at an intermediate value in ideal and realistic co-culture. Legend is consistent with panel B. **(E)** Parity plot of IL-2 concentration at final time point (t = 7 d) for all conditions in realistic (y-axis) vs ideal (x-axis) co-culture colored by each feature (column). Legend is consistent with panel B.

#### 2.3.2 Cell dynamics reveal potential new treatment strategy that spares healthy cells

Comparing trends in cell dynamics between ideal and realistic co-culture provides insight as to why each feature differentially impacts healthy cell killing. In ideal co-culture, increasing CAR affinity and cancer antigen expression level leads to healthy cell growth beyond their original numbers. Increasing CAR T-cell dose and CD4^+^:CD8^+^ ratio leads to healthy cell counts similar to those in the untreated control (**Supplementary Figure 3**). Interestingly, increasing CAR affinity results in more healthy cell growth compared to the case in which cancer cell antigen expression is increased. We hypothesize that this difference occurs because cancer cell killing is more strongly impacted by CAR affinity than cancer antigen density, providing healthy cells more opportunity to grow as more cancer cells die. However, in realistic co-culture, increasing cancer antigen level results in more healthy cell growth before being killed off compared to the scenario in which CAR affinity is increased. Cancer antigen expression primarily impacts cancer cell killing, which gives healthy cells the ability to grow before being targeted after cancer cell populations decline. Meanwhile, CAR affinity impacts both cancer and healthy cell killing, so healthy cells are killed at the same time as cancer cells. These data highlight how each feature differentially impacts the dynamics of this system. A large difference in cancer and healthy cell antigen levels can create a time delay between when cancer killing completes and when healthy cell killing starts, whereas tuning CAR affinity cannot create such a window. This time delay is an emergent phenomenon that occurs in some scenarios—it is not a trained, optimized, or hard-wired parameter in the model. One can design a strategy to take advantage of this time delay in scenarios when it occurs, for example, by deactivating CAR T-cells with an antibody or small-molecule induced off-switch that shuts down effector function after cancer cells are killed but before lower antigen expressing healthy cells are targeted.

#### 2.3.3 Individual feature analysis highlights tradeoffs in a Pareto curve

To quantify cancer and healthy cell killing, we use two metrics: normalized live healthy and cancer cell counts. The normalized count for each population (*N*_*P*_) is calculated as follows:

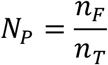

where *n*_*F*_ and *n*_*T*_ are the total number of live cancer or healthy cells at the final (t = 7 d) and treatment start (t = 0 d) timepoints, respectively. Values below one indicate net killing, and values above one indicate net growth. Together, these metrics place treatment outcomes within quadrants that can be used as guidelines for classifying efficacy (**Figure 3B**). Ideally, treatment conditions would appear in the upper left quadrant with maximal healthy cell sparing and maximal cancer cell killing. In both contexts, the trends match those of experimental observations—more aggressive treatments with more overall killing result from increasing CAR T-cell dose, intermediate CD4^+^:CD8^+^ ratio, increasing CAR-antigen affinity, and increasing cancer antigen density. These conditions allow for healthy cell maintenance or growth in ideal co-culture, nearing or entering the efficacious and selective treatment quadrant. However, in realistic co-culture, there exists a dramatic tradeoff between cancer cell killing and healthy cell killing, presenting a Pareto curve across each feature. Aggressive treatments exist toward the lower left quadrant (not selective for cancer cells). This observation suggests that it is not possible to optimize both efficacy and safety when healthy cells express antigen, and the most useful strategies—typically less aggressive treatments—balance these objectives (Caruso et al., 2015; Johnson et al., 2015; Liu et al., 2015).

### 2.4 IL-2 production is more strongly impacted by tuned features than context

IL-2 production is a standard *in vitro* measurements to quantify T-cell activation (Liu et al., 2015; Sommermeyer et al., 2016; Arcangeli et al., 2017). Similarly, glucose consumption can quantify T-cell activation through nutrient usage and competition (Frauwirth et al., 2002). We compare nutrient consumption and cytokine production across features and contexts to identify strategies for understanding, and potentially controlling, IL-2 production.

#### 2.4.1 Tuning CAR affinity modulates IL-2 production by balancing CAR T-cell proliferation and effector function

In `dish` (**Figure 3C** and **Supplementary Figure 9A** for monoculture; **Supplementary Figure 10A** for co-culture), IL-2 increases over time and with increasing values of CAR T-cell dose, CD4^+^:CD8^+^ ratio, CAR affinity, and cancer antigen expression level due to increased numbers of activated CD4^+^ CAR T-cells. Across all contexts and features, glucose decreases as IL-2 increases, indicating that glucose consumption follows CAR T-cell activation and proliferation (**Figure 3D, Supplementary Figure 9B, Supplementary Figure 10B**).

Unintuitively, IL-2 concentration is not maximized at the highest CAR-antigen affinity in monoculture where CAR T-cell activation is maximized. At the highest CAR affinity, more CAR T-cells spend time in effector, non-proliferative states (**Supplementary Figure 11**), resulting in fewer total CD4^+^ T-cells producing IL-2 (**Figure 2A**). This decrease is not observed in co-culture where cancer cell numbers are lower, reducing the likelihood that CAR T-cells will be activated. Decreased activation in co-culture produces lower IL-2 concentrations compared to monoculture. Thus, CAR T-cells in co-culture remain outside of the regime at which this tradeoff between activated and proliferating T-cells is observed. We hypothesize that maximum IL-2 production occurs at intermediate CAR affinity where these exists a balance between proliferation and frequent antigen binding. Excessively high CAR affinity leads to frequent target antigen binding, causing CAR T-cells to spend more time in effector rather than proliferating states, leading to fewer total CAR T-cells that can later produce cytokines. On the other hand, very weak affinity CARs drive cells primarily into states other than proliferative and effector states. Maximizing CAR-antigen affinity can therefore prove counterproductive for achieving CAR T-cell proliferation, survival, and cytokine production at the tumor site; moderate CAR-antigen affinities may be more effective.

#### 2.4.2 IL-2 production is independent of healthy cell antigen expression

In co-culture, healthy antigen expression minimally impacts IL-2 production and glucose consumption over time (**Figure 3D**). We speculate that healthy antigen expression is too low to strongly impact CAR T-cell proliferation and thus IL-2 production. Comparing final IL-2 concentration in all ideal versus realistic co-culture conditions reveal that IL-2 levels are independent of context for a given condition, further supporting this hypothesis (**Figure 3E**). CAR T-cell IL-2 production and overall glucose consumption are more strongly impacted by the higher level of antigen expression on the cancer cells than by the low antigen expression on healthy cells. When considering desired IL-2 levels produced by CAR T-cells in patient treatment, IL-2 production can be mostly attributed to and designed around cancer cells in isolation as healthy cell antigen expression does have a significant impact.

### 2.5 Multidimensional data analysis reveals context-specific treatment strategies

Since tuning individual features has different impacts on treatment efficacy based on the type of `dish`, we rank-ordered treatment outcomes across all individual simulated conditions, tuning all features simultaneously, within each context. Comparing the strongest treatments between monoculture and co-culture will enable us to determine how optimal treatments vary between contexts.

#### 2.5.1 Aggressive feature choices additively benefit treatment in monoculture

For monoculture, outcome is sorted by normalized live cancer cell count (**Figure 4A**). The best outcomes typically occur at the highest CAR T-cell doses, at a 25:75 CD4^+^:CD8^+^ ratio, at moderate to strong CAR affinity, and with high cancer cell antigen density. These trends are consistent with individual feature analyses in monoculture, and the same trends are observed in scenarios in which we considered expanded CAR T-cell doses (**Supplementary Figure 4A** and **4B**) and CD4^+^:CD8^+^ ratios (**Supplementary Figure 5F**). Choosing aggressive values for all features and using large E:T ratios yield cancer cell killing rates that are comparable with those observed in most experimental studies that use E:T ratios greater than one (i.e., killing most cancer cells occurs within hours), (**Supplementary Figure 4A**). Worse outcomes, in which cancer cells grow beyond their initial plated count, occur at low CAR T-cell doses, at 100:0 and 0:100 CD4^+^:CD8^+^ ratios, with the weakest CAR affinity, or with lower cancer cell antigen expression. Overall, combining aggressive choices for individual features additively benefits treatment outcome in monoculture. Effective CAR T-cell designs in the absence of healthy cells combine design choices from individually optimized features.

**FIGURE 4.**
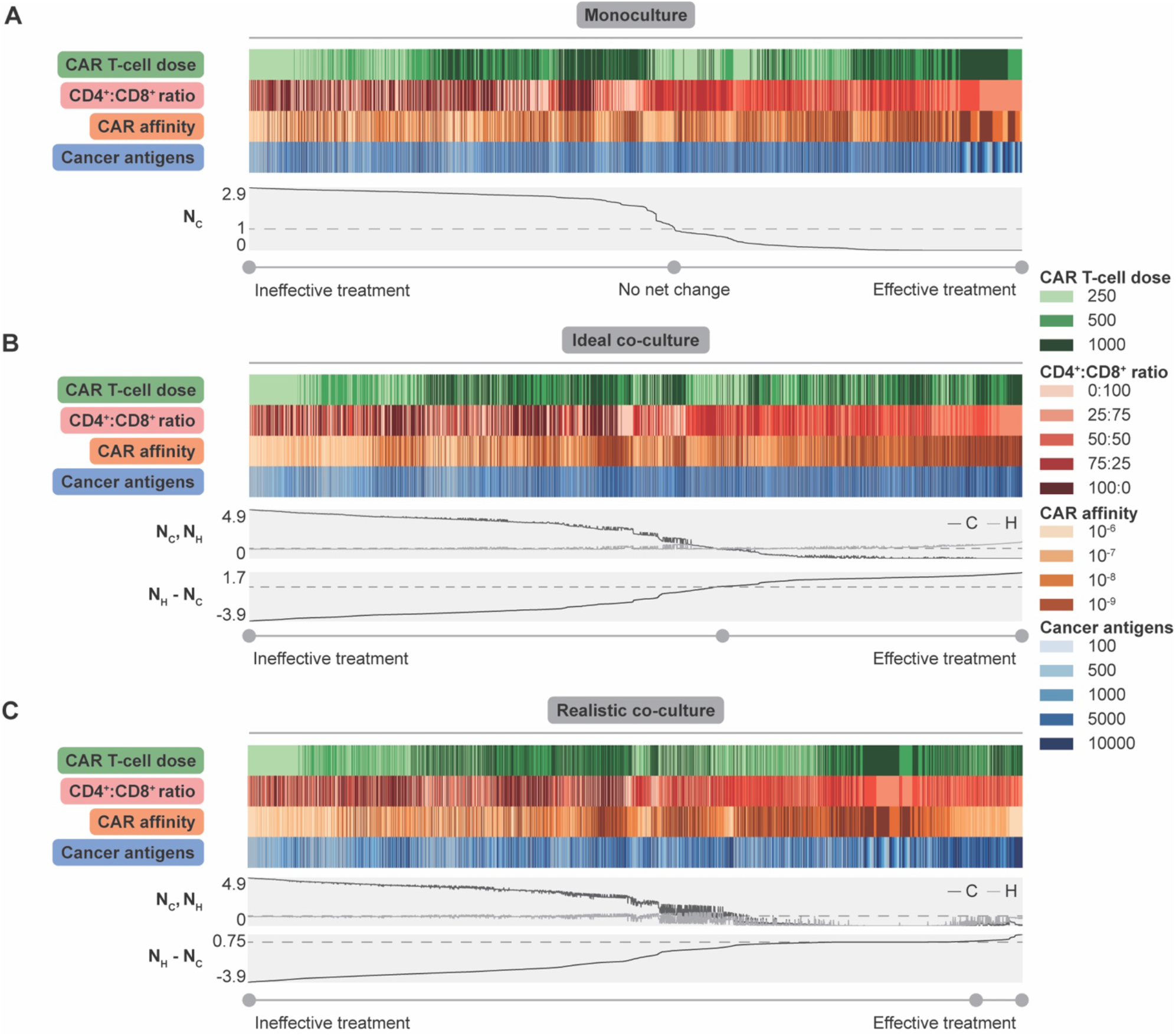
Collective impact of CAR T-cell and tumor features on dish outcomes. **(A)** Heatmap showing values for each feature with line plots showing normalized live cancer cell count (*N*_*C*_) sorted from highest (left) to lowest (right). The dashed line indicates value of *N*_*C*_ = 1, meaning no net change due to treatment. Values of *N*_*C*_ > 1 indicate net growth and values of *N*_*C*_ < 1 indicate net killing. **(B)** Heatmap showing values for each feature with line plots showing normalized live cancer cell count (*N*_*C*_) and normalized live healthy cell count (*N*_*H*_) (dashed line indicates normalized live cell count of 1) and the difference in normalized live healthy and cancer cell counts (*N*_*H*_ − *N*_*C*_) for each ideal co-culture simulation individually (dashed line indicates *N*_*H*_ − *N*_*C*_ = 0). The heatmap has been sorted from lowest (left) to highest (right) difference. All metrics were calculated at the final time point (t = 7 d). **(C)** Heatmap and normalized cell counts for realistic co-culture. Labels are consistent with panel B. Each feature is reported in the following units; CAR T-cell dose–500 CAR T-cells, CD4^+^:CD8^+^ ratio–50:50, CAR affinity—M, cancer antigens–1000 antigens/cell.

#### 2.5.2 Addressing off-target effects requires tuning multiple parameters

To identify general conclusions across diverse co-culture conditions, we considered treatment outcomes across all individual simulated conditions, sorted by the difference in the normalized live healthy and cancer cell count at the endpoint (**Figure 4B** for ideal co-culture, **Figure 4C** for realistic co-culture). This difference is maximized when healthy cells are spared and cancer cells are killed. We expect aggressive treatments to be most effective in the ideal cases, as healthy cells that do not express antigen cannot be killed. Trends in ideal co-culture match those in monoculture, supporting the idea that “invisible” healthy cells do not change observed trends.

However, the realistic co-culture where healthy cells express antigen, and can therefore be targeted by CAR T-cells, is more clinically relevant. In this context, there is a distinct tradeoff between cancer cell killing and healthy cell sparing. Conditions with the lowest normalized live cancer cell counts also show the lowest normalized live healthy cell counts (**Figure 4C**). Treatments with a positive difference all have some amount of healthy cell killing, but this killing is minimal compared to other conditions. Effective treatments have the highest doses of CAR T-cells, weaker CARs, CD4^+^:CD8^+^ ratios of 25:75 or 50:50, and higher cancer cell antigen count (**Figure 4C**). These observations agree with experimental findings that optimization of CAR T-cell therapy design yields different conclusions when balancing cancer cell killing and healthy cell sparing, versus focusing on the former objective alone (Caruso et al., 2015; Johnson et al., 2015; Liu et al., 2015). Though choosing high doses of weak CAR T-cells might seem unintuitive, using weak CARs minimizes the probability of targeting healthy cells while the high dose maximizes the probability that these weaker CARs successfully interact with high antigen density cancer cells. These results suggest that delivering higher doses of weaker CAR T-cells with CD4^+^:CD8^+^ ratios of 25:75 or 50:50 kill more cancer cells and spare more healthy cells for tumors where on-target off-tumor killing is undesired or detrimental.

### 2.6 Spatial dynamics drive vascularized tissue treatment efficacy

CAR T-cell therapy has great potential for use in solid tumor contexts, which include a complex tumor microenvironment, vasculature, spatial dynamics, and potentially antigen-expressing healthy cells. Predicting how the *in vitro* behavior conferred by various CAR T-cell designs corresponds to *in vivo* performance is not straightforward. We investigate the translation and efficacy of select treatment strategies in vascularized `tissue` where a solid tumor exists in a bed of antigen-expressing healthy cells within a dynamic microenvironment. We chose a subset of simulations—the realistic co-culture conditions deemed effective after averaging across replicates (**Supplementary Table 7**)—to analyze in `tissue`. Effective treatments were those that met the following two conditions: (1) cancer cells did not grow beyond the initial number, and (2) no more than 50% of the initial healthy cells were killed off.

A `tissue` is initialized with a bed of healthy cells in vascularized tissue that was inoculated with cancer cells and grown for 30 d. At t = 21 d, treatment began by adding a specified total dose of CAR T-cells, each expressing 5 × 10^4^ CARs with the given CAR affinity, and CD4^+^:CD8^+^ ratio. CAR T-cells were spawned at locations adjacent to vasculature to mimic intravenous trafficking to the tumor; they were not spawned adjacent to vessels that are too small in diameter for CAR T-cells to pass through. Files used to generate `tissue` simulations are described in **Supplementary Data 6** and **Supplementary Table 8**.

#### 2.6.1 Tested treatments are effective in tissue but differ in healthy cell killing

All treated tumors resulted in far fewer cancer cells and somewhat fewer healthy cells compared to untreated conditions, indicating that all strategies identified as effective in realistic co-culture proved effective in `tissue` (**Figure 5A**). In `dish`, healthy cell killing occurred primarily after most cancer cells were removed. This again motivates treatment strategies in which CAR T-cells include an inducible off-switch that shuts down effector function after cancer cells are killed but before lower antigen expressing healthy cells are targeted.

**FIGURE 5.**
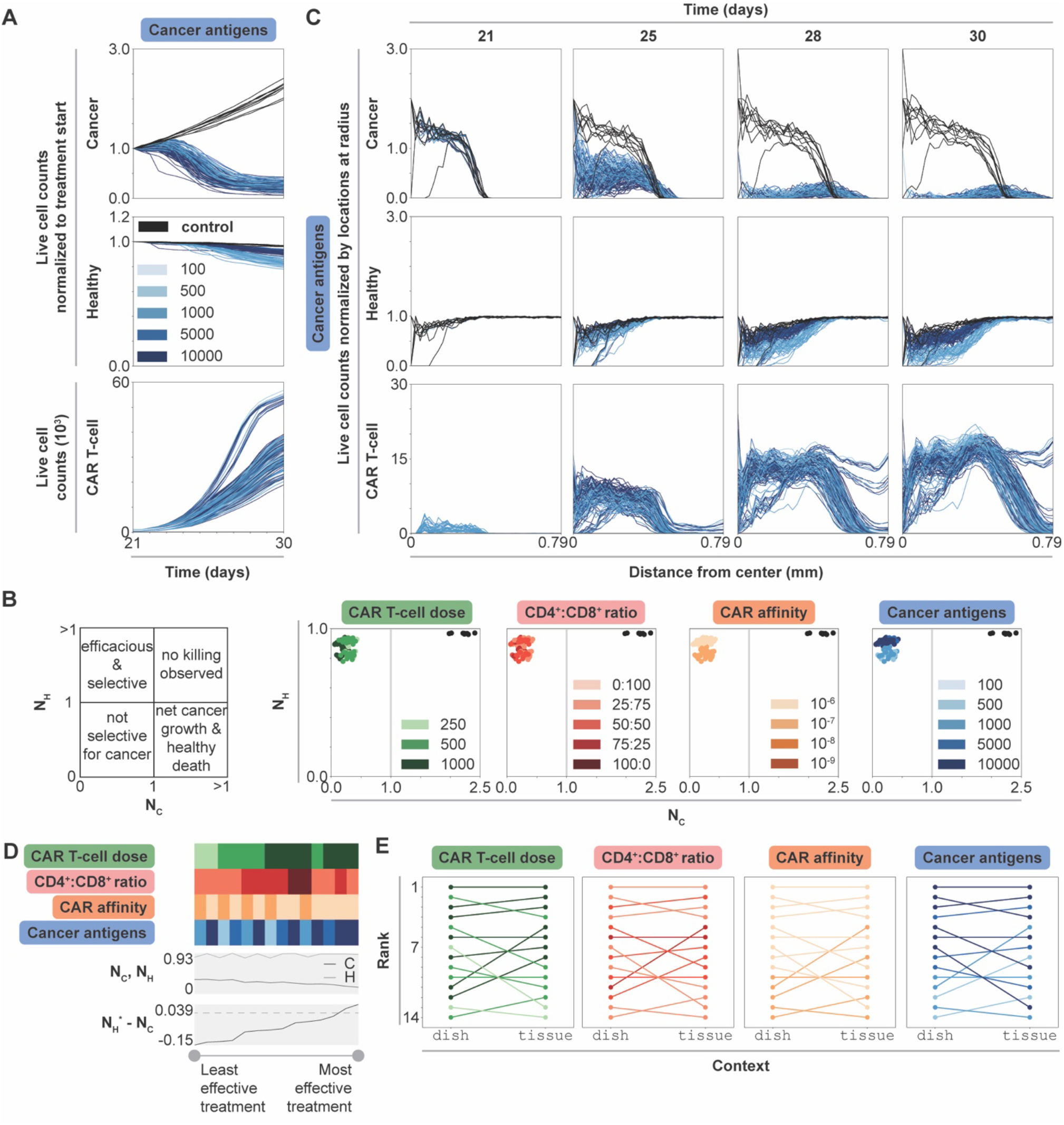
Dynamic, spatial, and ranked outcomes for selected promising treatment combinations in tissue. **(A)** Live cell counts over time of untreated (black) and treated conditions (graded hues) normalized to cell count at start of treatment (t = 21 d), for all simulations, colored by cancer antigens (other features may be changing as well). Cancer antigens reported in antigens/cell. The same data colored by other features are shown in **Supplementary Figure 12. (B)** Scatter plots of normalized live cancer cell count (*N*_*C*_, x-axis) vs. normalized live healthy cell count (*N*_*H*_, y-axis), each normalized to initial value at start of treatment (t = 21 d), for untreated (black) and treated conditions (graded hues) for all simulations, colored by one feature at a time. Each feature is reported in the following units; CAR T-cell dose–500 CAR T-cells, CD4^+^:CD8^+^ ratio–50:50, CAR affinity—M, cancer antigens–1000 antigens/cell. **(C)** Normalized live cell counts over time (t = 21 d, 25 d, 28 d, and 30 d shown) for untreated (black) and treated conditions (graded hues), normalized to locations per radius, for all simulations, colored by cancer antigens. The columns indicate the timepoint in the simulation (day), while the rows indicate cell type plotted, and the x-axis for each plot shows the distance from the center. Legend is consistent with panel B. **(D)** Heatmap showing values for each feature with line plots showing normalized live cancer and healthy cell counts and difference in normalized live healthy and cancer cell counts (*N*^*H*^ − *N*_*C*_, where healthy cell value is multiplied by the ratio of cancer to healthy cells at the start of treatment to ensure equal weighting since initial cell population sizes are not equal; dashed line indicates value of 0) at final time point averaged across replicates. The heatmap is sorted from lowest (left) to highest (right) difference. Feature legends are consistent with panels A and C. **(F)** Ladder plots of condition rankings in both `dish` and `tissue`, where condition outcome (averaged across replicates) is colored by each corresponding feature.

Comparing normalized live cancer and healthy cell counts at treatment endpoint enables direct comparison of treatment efficacy (**Figure 5B**). Notably, the primarily difference between treatment strategies is in degree of healthy cell killing. CAR affinity and cancer antigen expression, but not CAR T-cell dose or CD4^+^:CD8^+^ ratio, dictate this difference. In general, increasing CAR T-cell dose and using higher CD4^+^:CD8^+^ ratio treatments results in increased CAR T-cell counts, but it has little effect on cancer and healthy cell killing. Interestingly, most of the simulations that show the highest CAR T-cell production use the highest CAR T-cell doses and the weakest CAR affinity (**Supplementary Figure 12, Figure 5A**), indicating that designs with high doses of weak CAR T-cells result in the highest CAR T-cell growth rate *in vivo*. Overall, these observations reinforce the previously identified treatment strategy: use weaker CARs and select antigens with the highest differential between cancer and healthy cell expression. With this strategy, even though the CAR is weaker, the cancer antigen density is high enough to result in effective, selective treatment.

#### 2.6.2 Cancer cells with higher antigen density keep CAR T-cells near the tumor core

Though changing multiple features simultaneously complicates analysis, we noted an interesting pattern in which increasing cancer cell antigen density spares more healthy cells in tissue, representing a stark contrast to our `dish` findings. We thus investigated the spatial dynamics of each cell type to probe whether the mechanism by which CAR T-cells navigate within the solid tumor gives rise to this observation. At t = 21 d, cancer cells exist primarily in the center of the simulation, between the center and a radius of about 0.39 mm, while healthy cells are evenly spread across the simulation. (**Figure 5C**). In untreated conditions, cancer cells grow to cover a radius of 0.58 mm by t = 30 d and healthy cell count remains unchanged over time. In treated conditions, cancer and healthy cell counts decrease over time, primarily starting from the center where most CAR T-cells are initially spawned and moving outward. Cancer cell counts decrease with increasing cancer antigen density. CAR T-cell counts increase as a function of time and cancer cell count, but not as a function of cancer antigen density. Meanwhile, higher cancer antigen levels result in decreased healthy cell killing. We hypothesize that this phenomenon occurs when high antigen density cancer cells produce more CAR T-cells in effector states, leaving fewer CAR T-cells to migrate away from the center and toward the healthy cells. Thus, CAR T-cells that successfully traffic to a tumor core are more likely to remain localized and selectively target cancer cells when cancer cell antigen density is higher.

#### 2.6.3 Spatial differences between dish and tissue explain treatment performance

Comparing simulation rankings between `dish` and `tissue` reveals how context impacts treatment efficacy. To rank treatment strategies in `tissue`, we consider treatment outcomes across simulations (averaged across replicates) sorted from best to worst outcome in terms of difference in healthy and cancer cell counts normalized to start of treatment (**Figure 5D**). Nearly all highest ranked simulations use the highest CAR T-cell dose, a CD4^+^:CD8^+^ ratio of 25:75, the lowest CAR affinity, and the highest cancer antigen level. The four highest ranked treatment conditions in `dish` remain the four highest ranked treatment conditions in `tissue` (**Figure 5E, Supplementary Table 9**). The rankings for the mid and lower tier ranked simulations (5^th^-14^th^ in `tissue`) are shuffled from their original rankings in `dish`. One of the worst ranked treatments in `dish` (11^th^) jumped to 5^th^ in `tissue`, while a middle-ranked simulation (7^th^) fell to 13^th^ in `tissue`. These data predict that the most effective treatment conditions in `dish` will perform similarly in `tissue` assuming perfect CAR T-cell trafficking. Even with perfect trafficking, performance in `dish` does not exactly correlate with performance in `tissue`. Certain conditions may outperform in *in vivo* conditions compared to their performance *in vitro*.

Trends in how each feature impacts relative rank reveal which features most strongly dictate performance in `tissue` (**Figure 5E**). There are no distinct trends as a function of CD4^+^:CD8^+^ ratio. Most conditions that improve in rank use the relatively higher (though still objectively moderate) CAR affinity and higher CAR T-cell dose, and nearly all conditions that decrease in rank (from `dish` to `tissue`) have higher cancer antigen expression level. This finding is surprising given earlier observations that lowest CAR affinity with highest cancer antigen expression level combinations were most effective.

We hypothesize the differences in `dish` and `tissue` trends/rank result from differences in spatial dynamics. In `dish`, both healthy and cancer cells are well-mixed across the simulation, even after treatment, which results in an even spatial distribution of CAR T-cells (**Supplementary Figure 13**). In `tissue`, cancer cells sit in the center of the simulation surrounded by a large bed of healthy cells, with few healthy cells in the tumor core. When CAR T-cells are spawned with bias towards locations with more cancer cells to mimic perfect trafficking, the probability that spawn locations are adjacent to that of a healthy cell is higher in `dish` compared to `tissue`. Analyzing CAR T-cell state dynamics in both realistic co-culture `dish` and `tissue` for the selected promising treatment strategies further informs this spatial analysis. When we examine the distribution of CAR T-cell states only considering T-cells that are adjacent to a cancer cell (i.e., somewhat controlling for the local environment that a T-cell experiences), we find similar distributions of cells in effector states across `dish` and `tissue` simulations (**Supplementary Figure 14**). Thus, CAR T-cells in proximity to cancer cells exhibit similar behavior independent of experimental setup, and differences in overall trends/rank between contexts result from differences in collective cancer and healthy cell spatial distributions. Overall, this spatial difference in cancer and healthy cell distribution parallels comparisons between physical *in vitro* and *in vivo* experiments, even if CAR T-cell trafficking deviates from the perfect mechanism employed in our simulations, reinforcing the key role that spatial dynamics play in treatment outcome.

## 3 Discussion

We developed CARCADE as an open-access *in silico* testbed that enables systematic interrogation of the multidimensional design landscape of cellular engineering strategies, therapeutic optimization, and hypothesis generation. After verifying that the developed model recapitulates known trends *in vitro*, we explored design strategies in both `dish` and `tissue` contexts to gain insight into CAR T-cell design.

Tuning individual features in `dish` revealed key insights as to how these features impact CAR T-cell design. For example, we determined that healthy cell antigen expression results in healthy cell killing but has no impact on CAR T-cell or cancer cell dynamics. Modulating individual features recapitulated known tradeoffs between cancer cell killing and healthy cell sparing in realistic co-cultures. A new observation uniquely enabled by our model’s high resolution is that maximizing CAR affinity not only increases healthy cell killing but can also be counterproductive to CAR T-cell proliferation and cytokine production. In a related finding, we observed that IL-2 production is influenced more by tunable CAR T-cell design features than by healthy cell-related context.

Multidimensional analysis revealed that the relative performance of various treatment strategies is context dependent. Aggressive treatments are more effective in monoculture and ideal co-culture experiments, but effective treatment in realistic co-culture requires balancing all tuned features. We identified a particularly effective treatment strategy that balances cancer cell killing and healthy cell sparing when healthy cells express antigen. Specifically, we identified that the use of high doses of weak CAR T-cells with intermediate CD4^+^:CD8^+^ ratio and a maximized difference between cancer and healthy cell antigen expression produces the most effective treatments. By investigating these effective treatments in `tissue` context, we determined that differences in spatial distributions of cancer and healthy cells in `dish` and `tissue` contexts explain differences in treatment performance between contexts.

CARCADE is a first pass toward demonstrating the utility of models for generating hypotheses and informing design strategies for this class of problem, and it is important to consider that this model makes several assumptions and simplifications. First, the model is not tuned to a specific context. Results are general and might not hold in specific tumor contexts. A major strength of the model is that it can be easily tuned to a specific CAR and/or tumor type, and to interrogate specific design questions of interest. For example, CARCADE does not currently specify the CAR construct’s intracellular co-stimulatory domain (ICD), which is known to be an important factor in dictating CAR T-cell efficacy, persistence, and dynamics; rather, we approximate CAR behavior independent of ICD and find that broad trends hold despite not accounting for this factor explicitly. The model could be tuned to capture the effect of different ICD choices on CAR T-cell function. Similarly, the analysis can be tuned to change the definition of effective treatment outcomes to further penalize healthy cell killing (e.g., when considering treatments in which damage to CAR target antigen-expressing healthy cells is less tolerable from a safety standpoint). When treating B-cell cancers, off-tumor effects like B-cell aplasia are manageable with treatment, and healthy cell killing is less of a concern. In glioblastomas, EGFR is expressed on cancer cells, healthy brain cells, and other tissues, making healthy cell killing a greater risk of morbidity and mortality (Caruso et al., 2015). Another assumption made in the current CARCADE model is that there is no T-cell-mediated killing of bystander cells unless those bystander cells express the target antigen, which represents an ideal case. This assumption could easily be relaxed to interrogate the consequences of various forms of non-ideal T-cell killing. Additionally, the process by which CAR T-cells traffic to the tumor has been simplified and idealized, as CAR T-cells spawn at sites closest to cancer cells. The model could be adjusted to contemplate other scenarios, such as spawning CAR T-cells at the simulation edge while including CAR T-cell and environmental features that influence CAR T-cell trafficking to the tumor.

Expanding the agents, environment, or subcellular functions included in CARCADE offers opportunities for future model development and use in the field of CAR T-cell engineering. The present model comprises CAR T-cells and cancerous and healthy tissue cells; addition of macrophages, regulatory T-cells, natural killer cells, and other regulatory or supporting cell types or environmental factors could enable investigation of CAR T-cell therapy in a more complete and complex immune environment. In future studies, it may be particularly important to include the cell and environmental factors that contribute to immunosuppressive environments, as this is a common issue faced with *in vivo* CAR T-cell therapy. Additionally, while the current model was designed to facilitate analysis of treatments for solid tumors, particularly through the use of the `tissue` simulations, the `dish` simulations could be adapted to investigate liquid cancer treatment strategies. Future expansions could also incorporate trogocytosis, a processes by which CAR T-cells pick up tumor antigens from cancer cells and then experience fratricidal killing by other CAR T-cells, to investigate how this phenomena affects CAR T-cell persistence (Hamieh et al., 2019).

Overall, we believe that CARCADE will prove valuable for CAR T-cell designers and enable cross-cutting collaborations to facilitate further model development or tuning to specific contexts and questions of interest. By further refining the model using experimental data, CARCADE could help suggest potential promising strategies for experimental pursuit by testing strategies in `dish` and/or `tissue` contexts. CARCADE is designed to enable interrogation of questions and phenomena that are beyond the scope of the current study. For example, future studies using the current model could include a more granular consideration of CAR T-cell trafficking within the tumor by removing the assumption of perfect trafficking. Tumor immune escape could be investigated by creating multiple tumor subpopulations with variable antigen expression levels or susceptibly to killing. Additionally, inter-and intra-tumor heterogeneity could be integrated by simulating tumors that comprise multiple populations with different parameters and/or differing levels of heterogeneity. Ultimately, integrating CARCADE into the CAR T-cell design process could accelerate the design-build-test cycle, saving resources and time associated with new therapeutic development.

## 4 Materials and Methods

CARCADE was developed by extending ARCADE, an existing multi-scale, multi-class agent-based model that includes tissue cells and hemodynamic environments. We used CARCADE to generate *in silico* experiments where we treated monoculture `dish`, ideal and realistic co-culture `dish`, and `tissue` contexts with CAR T-cells. All model details, including adaptation of tissue cell agents, development of CAR T-cell agents, development and adaptation of subcellular modules, development of `dish` plating, and all simulation setups and analyses are described in detail in the Supplementary Methods Details section of the **Supplementary Material** (Kuse et al., 1985; Lauffenburger and Linderman, 1993; Robertson et al., 1996; Schwartz, 1996; Frauwirth et al., 2002; De Boer et al., 2003;

Deenick et al., 2003; Iwashima, 2003; Schwartz, 2003; Chmielewski et al., 2004; Macian et al., 2004; Janas et al., 2005; Jacobs et al., 2008; Busse et al., 2010; Malek and Castro, 2010; Pearce, 2010; Yoon et al., 2010; Akbar and Henson, 2011; Wang et al., 2011; Wherry, 2011; Altman and Dang, 2012; Gerriets and Rathmell, 2012; Robertson-Tessi et al., 2012; Stone et al., 2012; Cheng et al., 2013; Crespo et al., 2013; Hegde et al., 2013; Liao et al., 2013; MacIver et al., 2013; Rosenberg, 2014; Buck et al., 2015; Heskamp et al., 2015; Kinjyo et al., 2015; Liadi et al., 2015; Liu et al., 2015; Long et al., 2015; Obst, 2015; Wherry and Kurachi, 2015; Chang and Pearce, 2016; Cherkassky et al., 2016; Golubovskaya and Wu, 2016; Harris and Kranz, 2016; Hegde et al., 2016; Liu et al., 2016; Maus and June, 2016; Sommermeyer et al., 2016; Verbist et al., 2016; Arcangeli et al., 2017; Borghans and Ribeiro, 2017; Gong et al., 2017; Mehta et al., 2017; Gherbi et al., 2018; Guedan et al., 2018; Huang et al., 2018; Kasakovski et al., 2018; Rafiq et al., 2018; Ross and Cantrell, 2018; Salter et al., 2018; Watanabe et al., 2018; Yost et al., 2019; Yu and Bagheri, 2020; Hernandez-Lopez et al., 2021; Yu and Bagheri, 2021).

The agent-based model developed during this study is available at the Bagheri Lab GitHub CARCADE repository: https://github.com/bagherilab/CARCADE.

The scripts developed to process and analyze the data during this study will be published along with the final manuscript at the Bagheri Lab GitHub carcade_mapping_design_space repository: \https://github.com/bagherilab/carcade_mapping_design_space.

## 6 Author Contributions

A.N.P., J.N.L, and N.B. conceived the project. A.N.P developed CARCADE by extending the ARCADE model, originally created by J.S.Y. J.S.Y. provided mentorship on the model development process. A.N.P. designed and conducted *in silico* experiments; processed data; and analyzed and plotted results. A.N.P. prepared the initial draft of the manuscript; A.N.P., J.S.Y., J.N.L, and N.B. wrote, refined, and edited the manuscript. J.N.L and N.B. supervised the project.

## Acknowledgments

The authors acknowledge computational resources and staff contributions by the Quest high performance computing facility at Northwestern University, which is jointly supported by the Office of the Provost, the Office for Research, and the Northwestern University Information Technology. The authors also thank members of the Leonard and Bagheri Labs for invaluable project discussions and feedback.

## 9 Data Availability Statement

The *in silico* datasets generated and analyzed for this study can be provided upon request.

## 10 Supplementary Material

The Supplementary Material for this article are included with the online publication and include (i) an extended dataset of monoculture simulations with two additional CD4^+^:CD8^+^ ratios tested (**Supplementary Notes**); (ii) a detailed description of the methods including details on the model framework, tissue cell agents, new CAR T-cell agents, model environment, cell placement and treatment, simulated experiment setups, and data analysis (**Supplementary Methods Details**); (iii) figures describing CAR T-cell agent function, details on monoculture/co-culture/tissue simulation time courses and treatment outcomes not shown in main text, such as cell counts and states over time, cell cycle and volume distributions, and environmental IL-2 and glucose concentrations over time, and `dish` spatial dynamics (**Supplementary Figures**); (iv) and tables showing model parameters, simulation setups, and co-culture/tissue treatment ranking and live normalized cell outcomes (**Supplementary Tables**).

