## Supplementary Materials for "Mapping CAR T-cell design space using agent-based models"

### 1 Supplementary Data

**SUPPLEMENTARY DATA 1. Simulated monoculture `dish` setup files.** Compressed (.zip) files containing .xml files needed to create untreated and treated monoculture `dish` simulations. Files correspond to those in **Supplementary Table 4**.

**SUPPLEMENTARY DATA 2. Simulated co-culture `dish` setup files.** Compressed (.zip) files containing .xml files needed to create untreated and treated co-culture `dish` simulations. Files correspond to those in **Supplementary Table 5**.

**SUPPLEMENTARY DATA 3. Simulated monoculture `dish` setup files with expanded CAR T-cell doses (E:T ratios).** Compressed (.zip) files containing .xml files needed to create untreated and treated monoculture `dish` simulations with expanded CAR T-cell dose range.

**SUPPLEMENTARY DATA 4. Simulated monoculture `dish` setup files with expanded $CD4^+$:$CD8^+$ ratios.** Compressed (.zip) files containing .xml files needed to create untreated and treated monoculture `dish` simulations with expanded $CD4^+$:$CD8^+$ ratios.

**SUPPLEMENTARY DATA 5. Normalized antigen and percent lysis from cited papers shown in Figure 2G.** This document (.xlsx) details estimated antigen and lysis data from the cited papers as well as the calculations and values of the normalized antigen and percent lysis for each paper. The figure from the paper where information was drawn and notes about the experimental context are noted for each reference.

**SUPPLEMENTARY DATA 6. Simulated `tissue` setup files.** Compressed (.zip) files containing .xml files needed to create untreated and treated `tissue` simulations and corresponding graph simulations. Files correspond to those in **Supplementary Table 8**.

### 2 Supplementary Notes

#### 2.1 Supplementary Note 1

Here we provide additional rationale as to how the range of ratios of $CD4^+$:$CD8^+$ CAR T-cells was selected following initial investigation in **Figure 2** and **Supplementary Figure 3**. In these studies, expanding the range of tested $CD4^+$:$CD8^+$ ratios to include 90:10 and 10:90 in monoculture and co-culture, these more extreme ratio cases further validate trends observed in the originally tested set (**Supplementary Figure 4**). The 90:10 ratio behaves in a way that is intermediate between the 75:25 and the 100:0 ratios, and the 10:90 ratio behaves similarly to the 25:75 ratio for trends across cell dynamics and killing (**Supplementary Figure 4A-B**), IL-2 production (**Supplementary Figure 4C-D**), and the holistic datasets (**Supplementary Figure 4E-F**). The strong similarity in treatment efficacy between the 25:75 and the 10:90 ratio cases in `dish` indicates that they may perform similarly in `tissue`. The choice of which strategy to pursue *in vitro* or *in vivo* may eventually be guided by cost or feasibility with respect to $CD4^+$ and $CD8^+$ acquisition and culturing.

### 3 Supplementary Figures and Tables

#### 3.1 Supplementary Figures

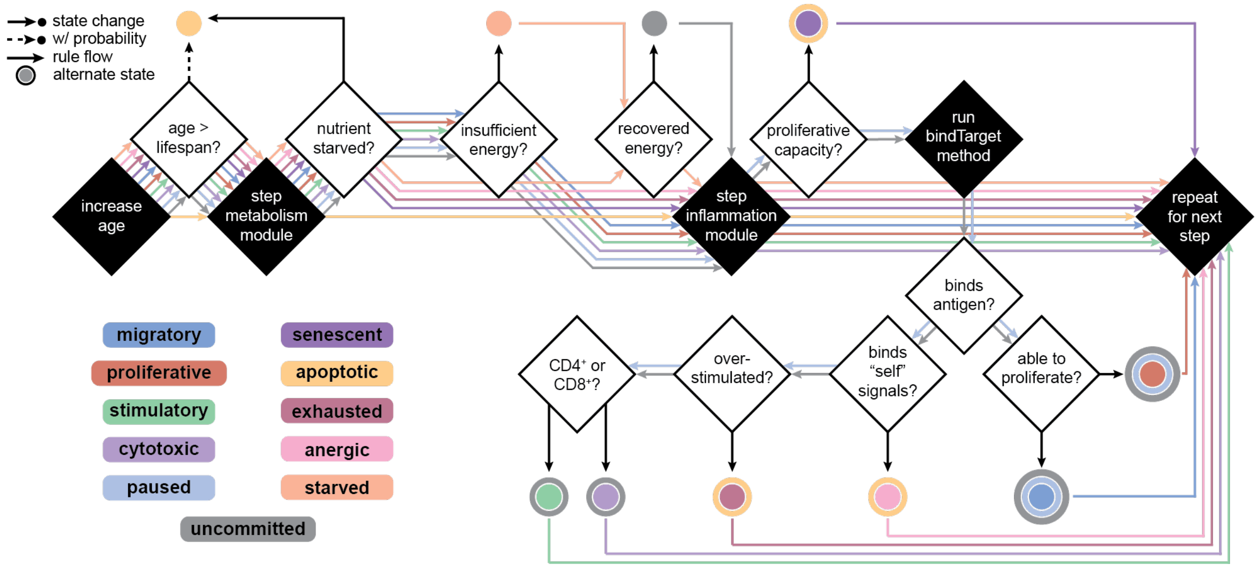

**SUPPLEMENTARY FIGURE 1. CAR T-cell state diagram flow chart.** This flowchart outlines the rules governing CAR T-cell state transitions as a function of current cell state at each tick of the simulation. Black boxes indicate actions and white boxes indicate checks cells perform to change state. Some state transitions occur with probability, as indicated in the legend. Each agent first increases their age and then compares their age to their defined lifespan. If it is older than the lifespan, then the cell will become apoptotic with a given probability, and this probability increases with the age difference above the lifespan. Next, the metabolism module determines cell nutrient uptake and energy level. CAR T-cells that are nutrient starved become apoptotic. Those with low energy, but not below the starvation threshold, become starved until they can recover this energy or die. CAR T-cell agents then step their inflammation modules to determine how much IL-2 in the environment they've bound to, which has downstream effects on their metabolism and effector functions. Cells then either commit to a new cell state based on their surroundings or remain in the cell state that they are already committed to until their action is completed or they die. Cells in uncommitted or paused states will assess their surroundings and calculate the probability with which they bind to a neighboring tissue cell, giving the possibility of becoming activated (stimulatory or cytotoxic depending on cell type), dysfunctional (exhausted or anergic), or failing to bind and thus either migrating or proliferating. Each state is described in more detail in the **Supplementary Methods Details**.

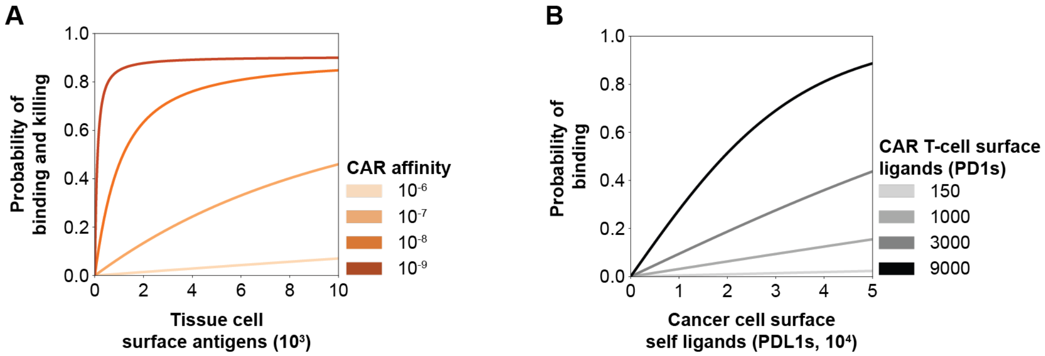

**SUPPLEMENTARY FIGURE 2. Binding probability heuristic function simulations.** Binding probability heuristic function simulations for **(A)** binding and killing when CAR T-cells binding to target cell where CAR affinity is reported in units of M and **(B)** probability of CAR T-cell binding to self-ligands on cancer cell surface where CAR T-cell surface ligands and self-ligand are reported in units of ligands/cell.

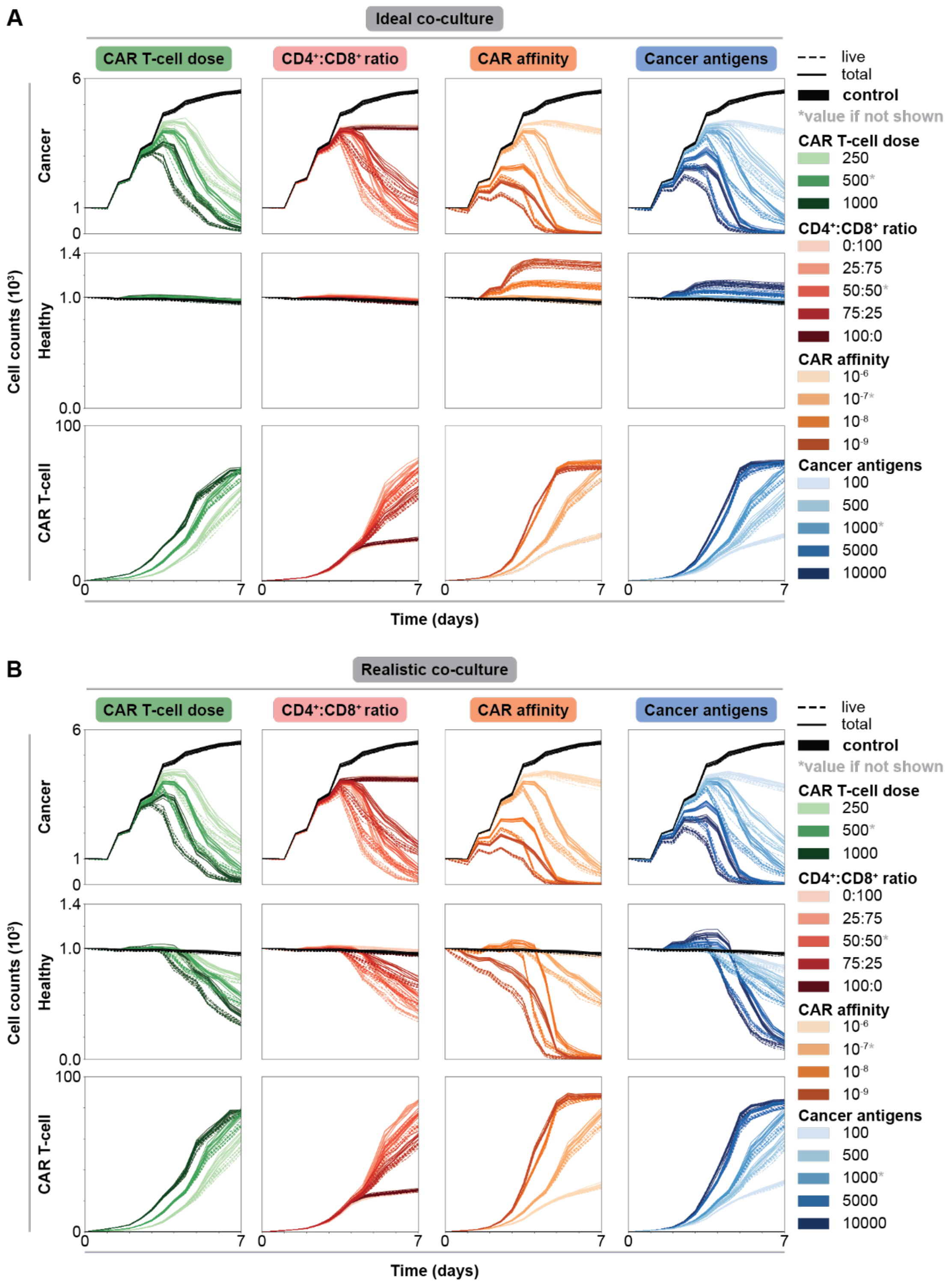

**SUPPLEMENTARY FIGURE 3. Impact of individual CAR T-cell and tumor features on cell dynamics in an ideal and realistic co-culture dish.** Cell counts over time for **(A)** ideal and **(B)** realistic co-cultures. Each column shows the axis being changed, where all other features are held constant at indicated intermediate value (indicated by asterisk, CAR T-cell dose–500 CAR T-cells, $CD4^+$:$CD8^+$ ratio–50:50, CAR affinity–$10^{-7}$ M, cancer antigens–1000 antigens/cell) while rows show the cell type being plotted. The solid lines represent total cell counts (total), while dashed lines represent live cell counts (live, excludes necrotic and apoptotic states).

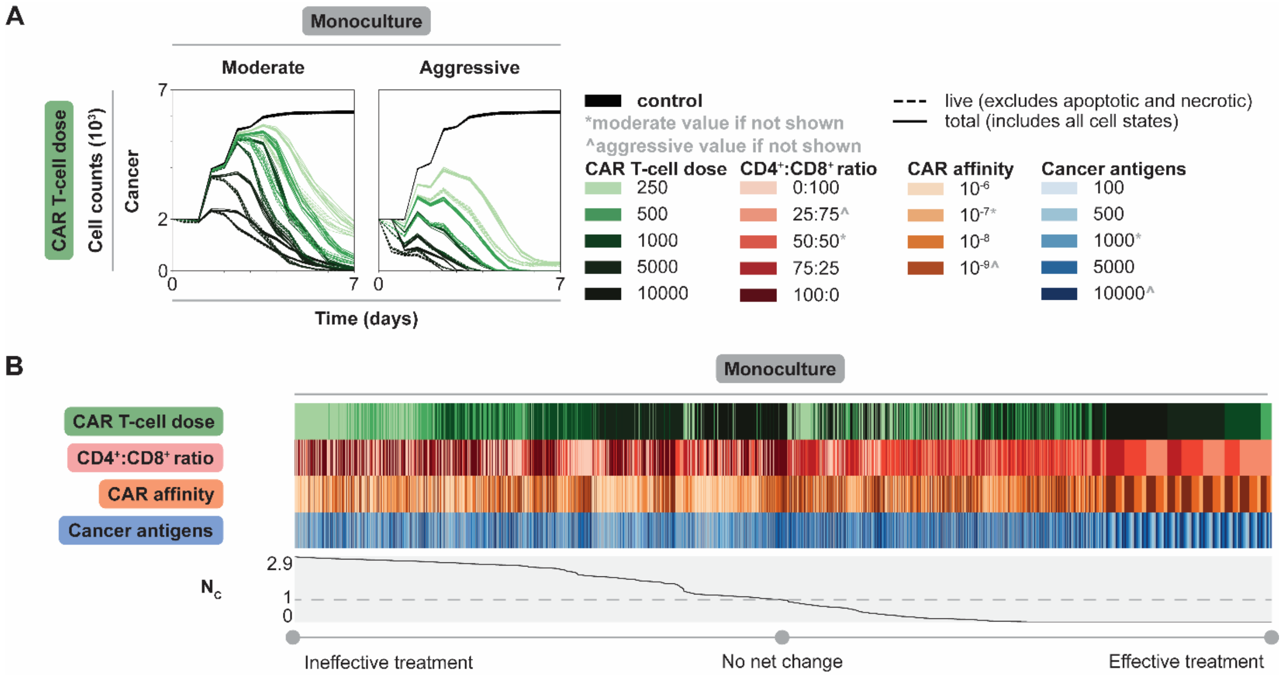

**SUPPLEMENTARY FIGURE 4. Increasing the effector-to-target (E:T) ratio results in increased and accelerated cancer cell killing. (A)** Cancer cell counts over time colored by CAR T-cell dose with all other features set to moderate (indicated by asterisk, CD4$^+$:CD8$^+$ ratio–50:50, CAR affinity–10$^{-7}$ M, cancer antigens–1000 antigens/cell) or aggressive (indicated by carrot, CD4$^+$:CD8$^+$ ratio–25:75, CAR affinity–10$^{-9}$ M, cancer antigens–10000) values. **(B)** Heatmap reports values for each feature; the corresponding line plot shows normalized live cancer cell count ($N_C$) sorted from highest (left) to lowest (right). The dashed line indicates a value of 1, meaning no net change due to treatment. Values of $N_C$ above 1 indicate growth and values below 1 indicate net killing. Legend is consistent with panel A.

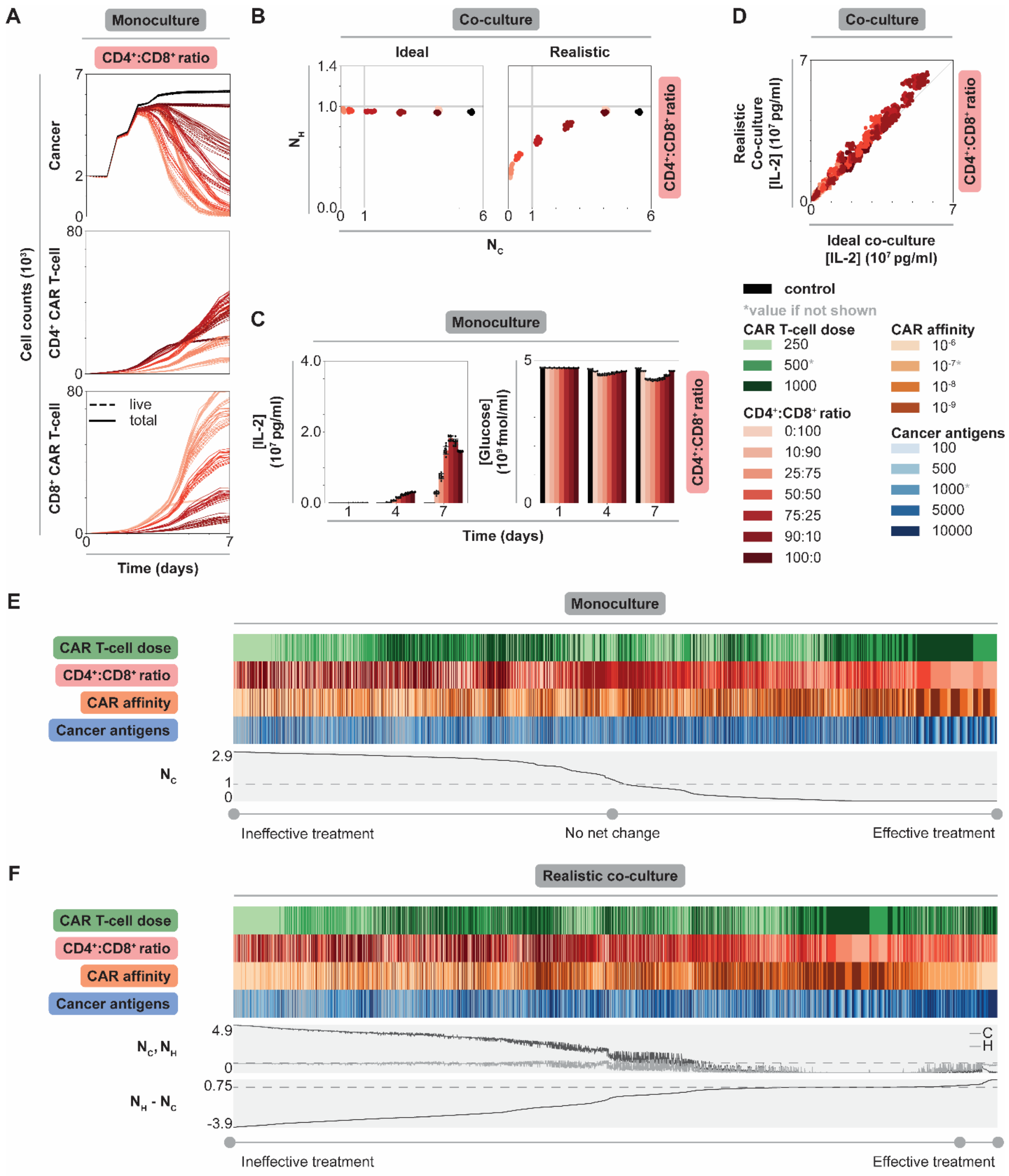

**SUPPLEMENTARY FIGURE 5. Treatment outcomes for additional edge case CD4$^+$:CD8$^+$ ratio conditions in monoculture and co-culture. (A)** Cell counts over time of untreated (black) and treated conditions (graded hues) holding all but CD4$^+$:CD8$^+$ ratio constant at an intermediate value (indicated by asterisk, CAR T-cell dose–500 CAR T-cells, CAR affinity–10$^{-7}$ M, cancer antigens–1000 antigens/cell) in monoculture. Each row shows the cell type being plotted. The solid lines represent total cell counts (total), while dashed lines represent live cell counts (live, excludes necrotic and apoptotic states). Legend is consistent with panel C. **(B)** Scatter plots of normalized live cancer cell count ($N_C$, x-axis) vs normalized live healthy cell count ($N_H$, y-axis) for untreated (black) and treated conditions (graded hues) holding all but CD4$^+$:CD8$^+$ ratio constant at an intermediate value. Data are separated by ideal and realistic co-culture context. Legend is consistent with panel C. **(C)** IL-2 and glucose concentrations over time holding all but CD4$^+$:CD8$^+$ ratio constant at an intermediate value in the monoculture context. **(D)** Parity plot of IL-2 concentration at t = 7 d for all conditions in realistic (y-axis) vs ideal (x-axis) co-culture contexts colored by CD4$^+$:CD8$^+$ ratio. **(E)** Heatmap showing values for each feature with line plots showing normalized live cancer cell count ($N_C$) sorted from highest (left) to lowest (right). The dashed indicates value of 1, meaning no net change due to treatment. Values of $N_C$ above 1 indicates growth and below 1 indicates net killing. Legend is consistent with panel C. **(F)** Heatmap showing values for each feature with line plots showing normalized live cancer and healthy cell count ($N_C$ and $N_H$, respectively) (dashed line indicates value of 1) and the difference in normalized live healthy and cancer cell counts ($N_H - N_C$) for each realistic co-culture simulation individually (dashed line indicates value of 0). The heatmap has been sorted from lowest (left) to highest (right) difference. All the metrics shown were calculated at final time point (t = 7 d). Legend is consistent with panel C.

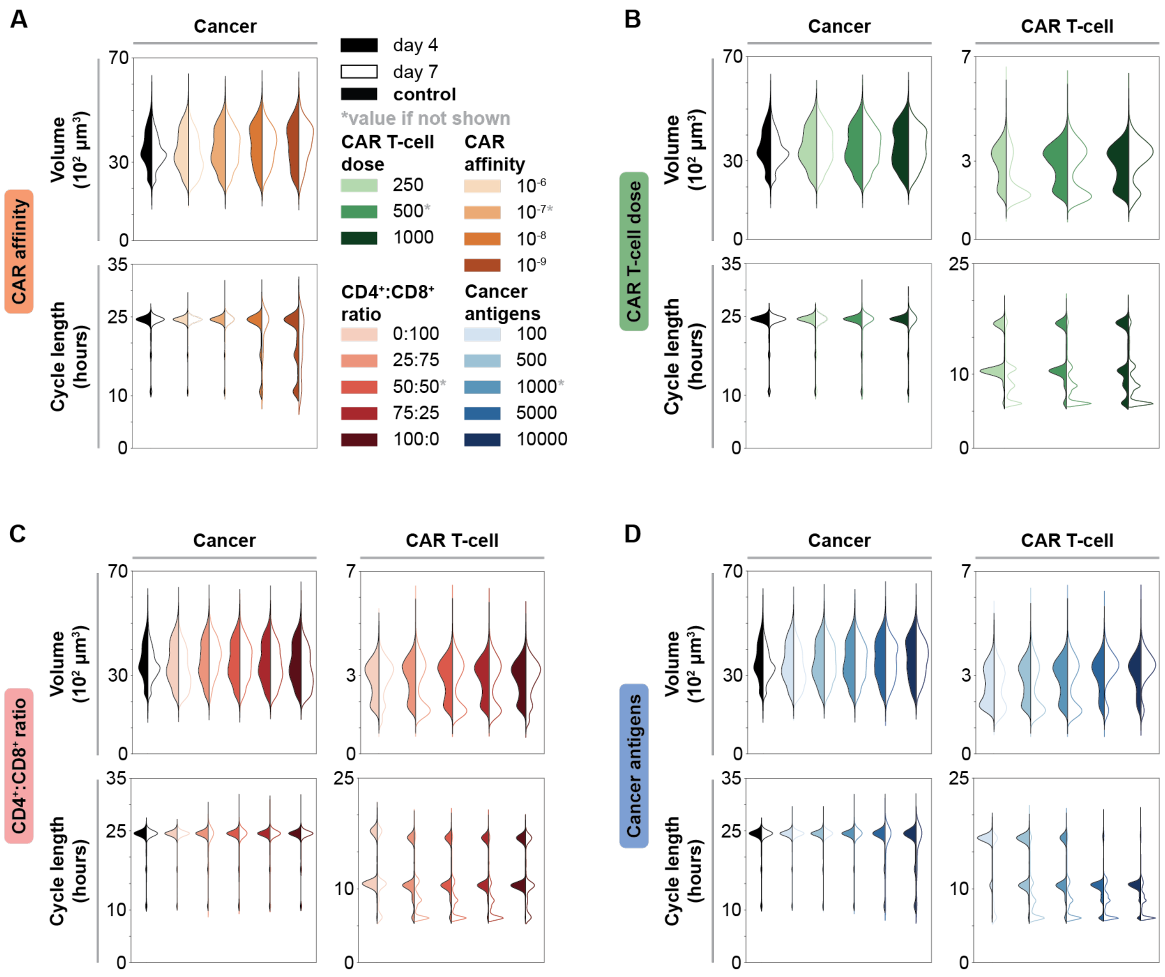

**SUPPLEMENTARY FIGURE 6. Impact of individual CAR T-cell and tumor features on cancer and CAR T-cell volume and cell cycle distributions over time in monoculture. (A)** Volume and cell cycle length distributions for cancer cells when changing CAR affinity. Volume and cell cycle length distributions for cancer and CAR T-cell populations when changing **(B)** CAR T-cell dose, **(C)** CD4$^{+}$:CD8$^{+}$ ratio, and **(D)** cancer antigens. Legend for all is consistent with panel A. When one feature is changing, all other features are held constant at indicated intermediate value (indicated by asterisk, CAR T-cell dose–500 CAR T-cells, CD4$^{+}$:CD8$^{+}$ ratio–50:50, CAR affinity–10$^{-7}$ M, cancer antigens–1000 antigens/cell). Each column shows the cell type being plotted, while rows show the distribution being plotted. Within each distribution, data are shown at t = 4 d (filled) and t = 7 d (outline).

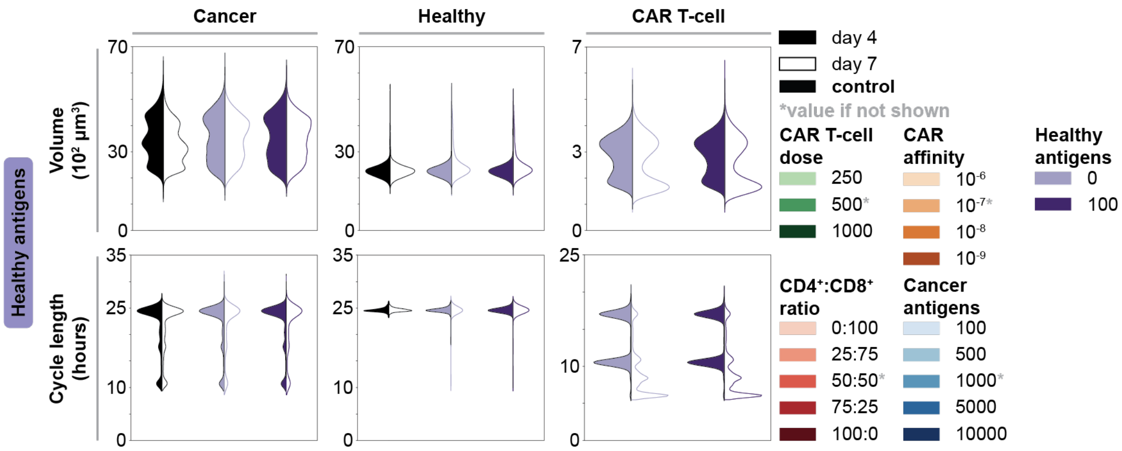

**SUPPLEMENTARY FIGURE 7. Impact of changing healthy cell antigen expression on cell volume and cell cycle distributions in co-culture.** All features are held constant at indicated intermediate value (indicated by asterisk, CAR T-cell dose–500 CAR T-cells, $CD4^+$:$CD8^+$ ratio–50:50, CAR affinity–$10^{-7}$ M, cancer antigens–1000 antigens/cell). Healthy antigens shown in units of antigens/cell. Each column shows the cell type being plotted, while rows show the distribution being plotted. Within each distribution, data are shown at t = 4 d (filled) and t = 7 d (outline).

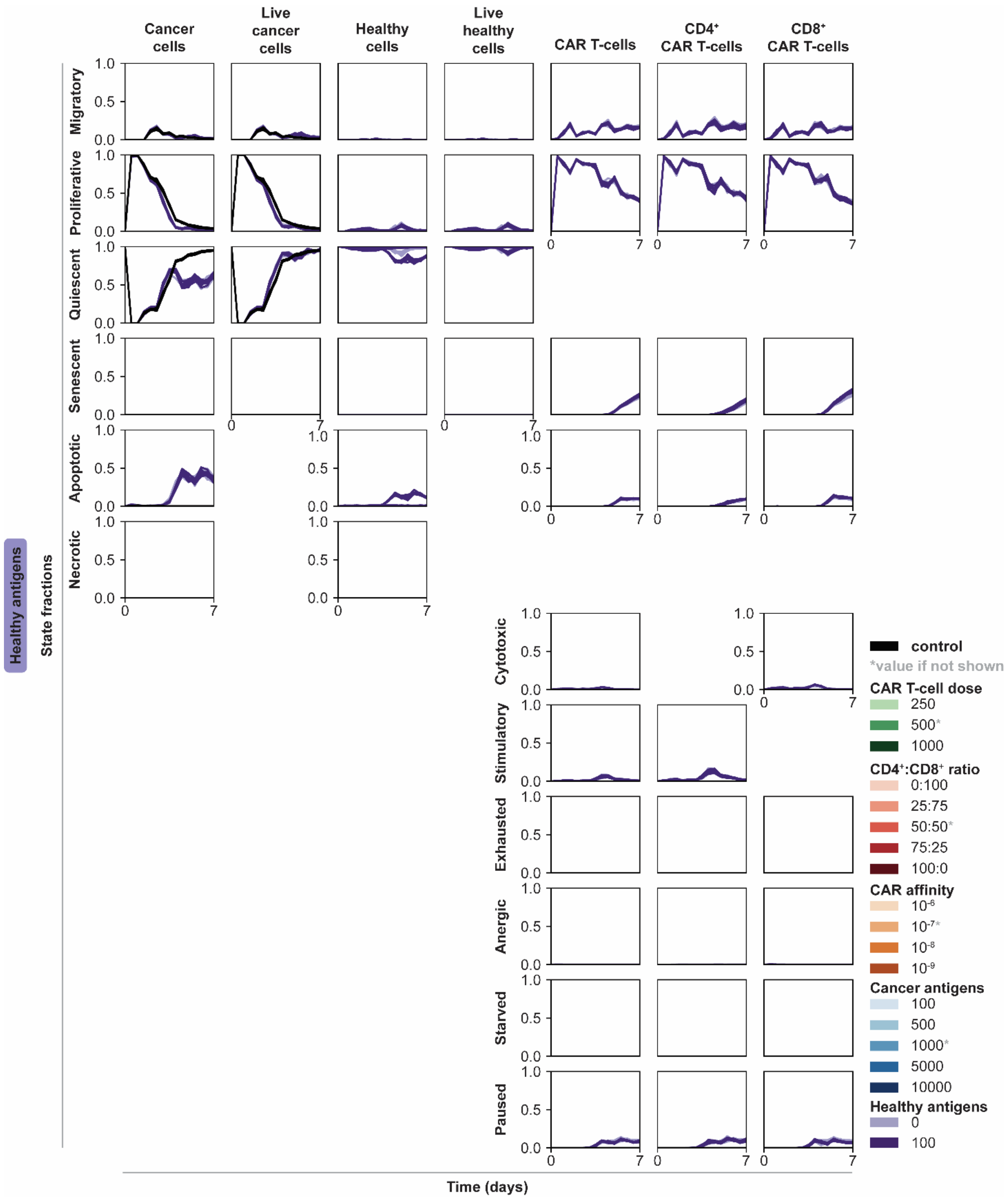

**SUPPLEMENTARY FIGURE 8. Impact of changing healthy cell antigen expression on cell states in co-culture.** All features are held constant at indicated intermediate value (indicated by asterisk, CAR T-cell dose–500 CAR T-cells, $CD4^+$:$CD8^+$ ratio–50:50, CAR affinity–$10^{-7}$ M, cancer antigens–1000 antigens/cell). Healthy antigens reported in units of antigens/cell. Each column shows the cell type being plotted, while rows show the cell state fraction being plotted.

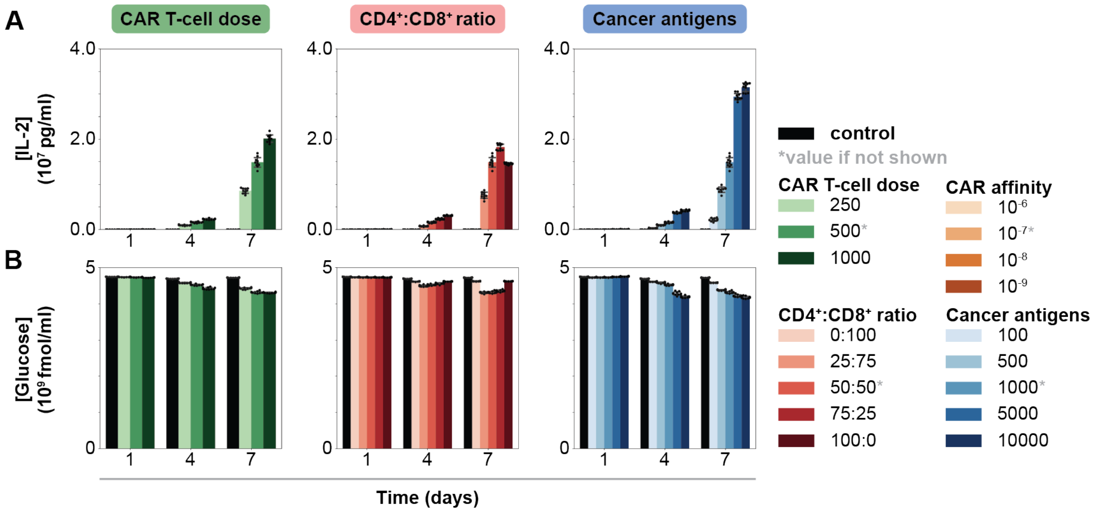

**SUPPLEMENTARY FIGURE 9. Impact of changing CAR T-cell and tumor features on IL-2 and glucose concentrations in monoculture. (A)** IL-2 and **(B)** glucose concentrations over time as each specified feature is varied. Each column shows the axis being changed, where all other features are held constant at indicated intermediate value (indicated by asterisk, CAR T-cell dose–500, $CD4^+$:$CD8^+$ ratio–50:50, CAR affinity–$10^{-7}$ M, cancer antigens–1000 antigens/cell), while rows show the environmental species being plotted.

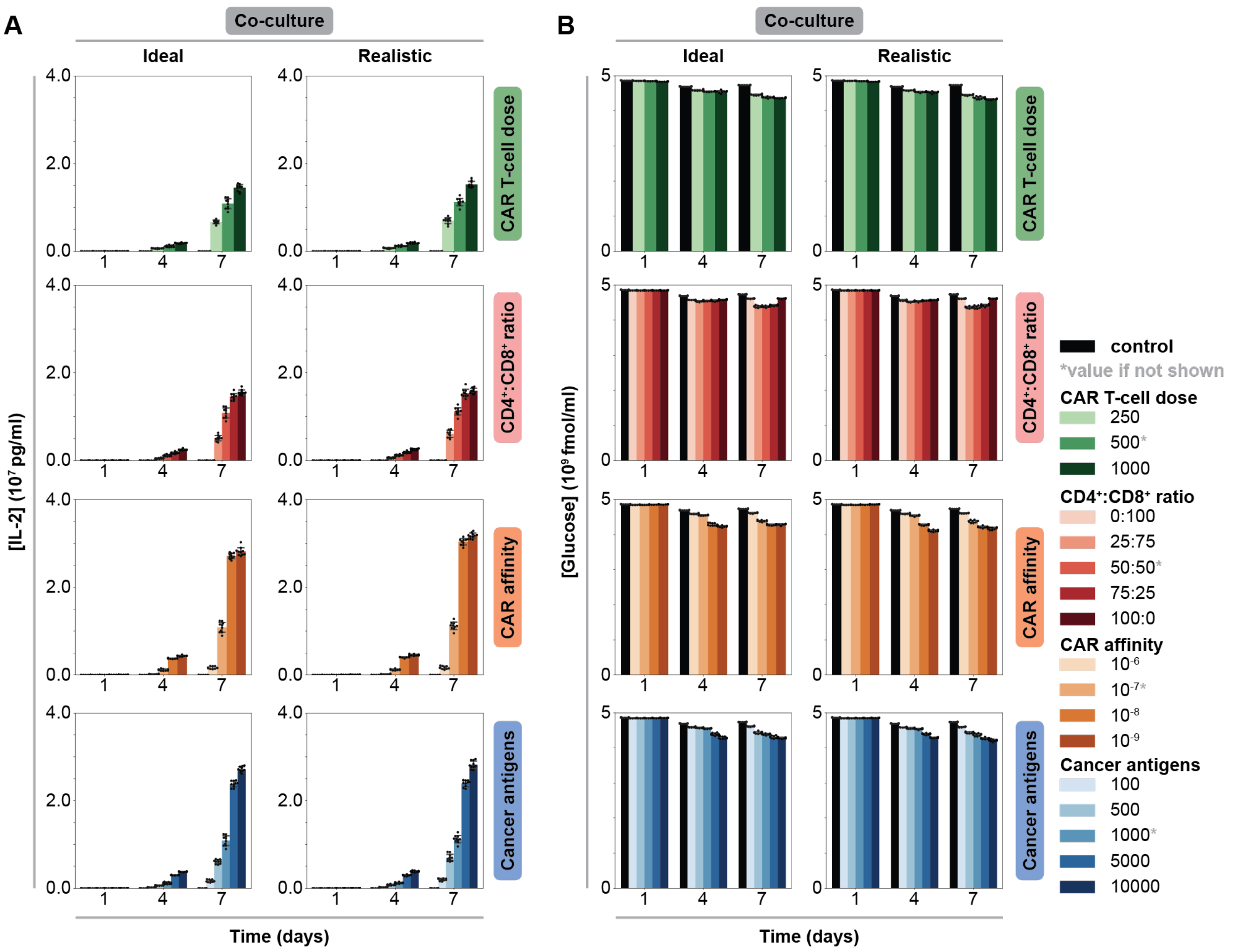

**SUPPLEMENTARY FIGURE 10. Impact of changing CAR T-cell and tumor features on IL-2 and glucose concentrations in co-culture. (A)** IL-2 and **(B)** glucose concentrations over time as each specified feature is varied. Each column shows the co-culture context, while rows show the axis being changed, where all other features are held constant at indicated intermediate value (indicated by asterisk, CAR T-cell dose–500 CAR T-cells, $CD4^+$:$CD8^+$ ratio–50:50, CAR affinity–$10^{-7}$ M, cancer antigens–1000 antigens/cell).

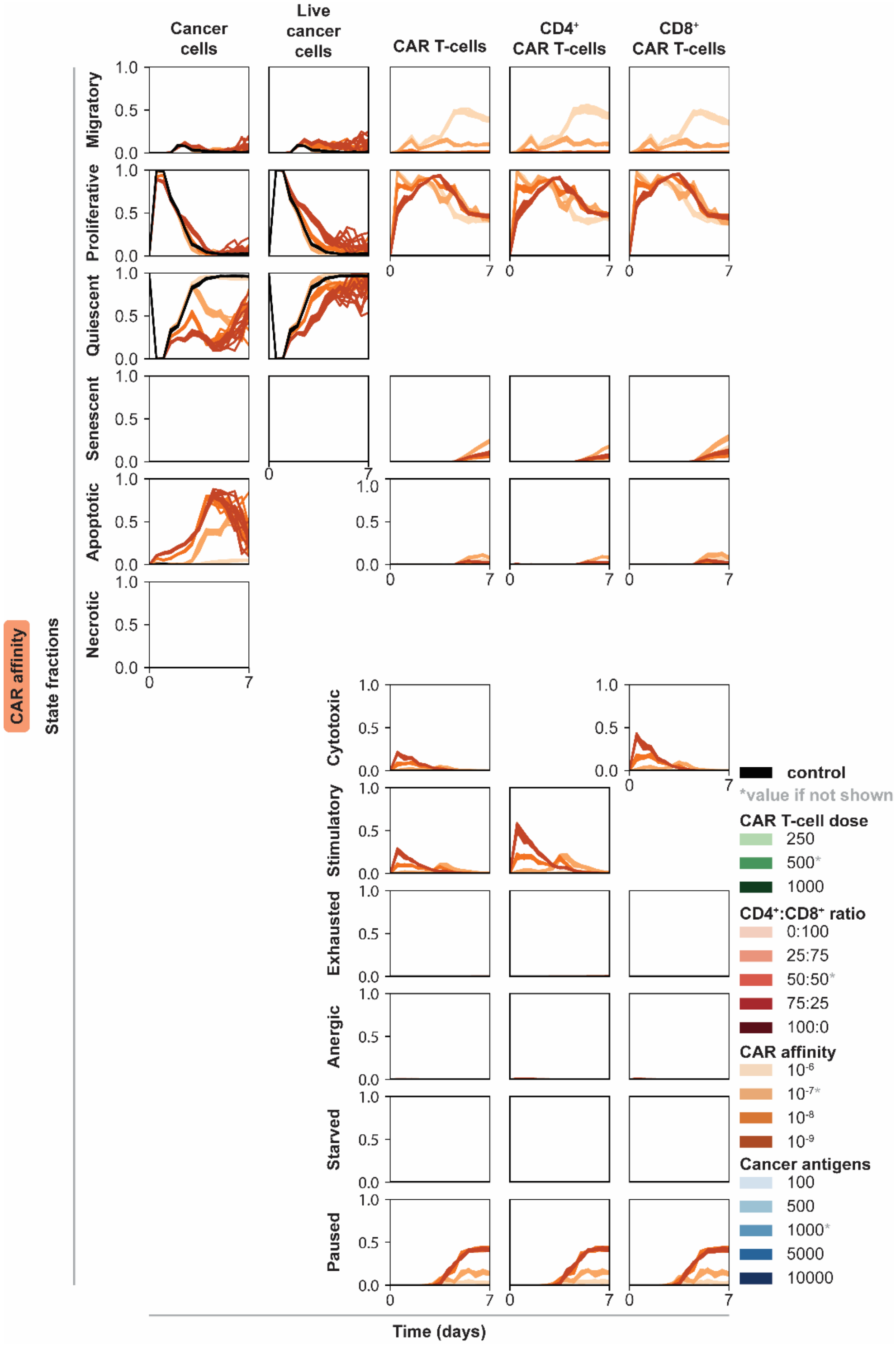

**SUPPLEMENTARY FIGURE 11. Impact of changing CAR affinity cell states over time in monoculture.** All other features are held constant at indicated intermediate value (indicated by asterisk, CAR T-cell dose–500 CAR T-cells, $CD4^{+}$:$CD8^{+}$ ratio–50:50, cancer antigens–1000 antigens/cell). CAR affinity reported in units of M. Each column shows the cell type being plotted, while rows show the cell state fraction being plotted.

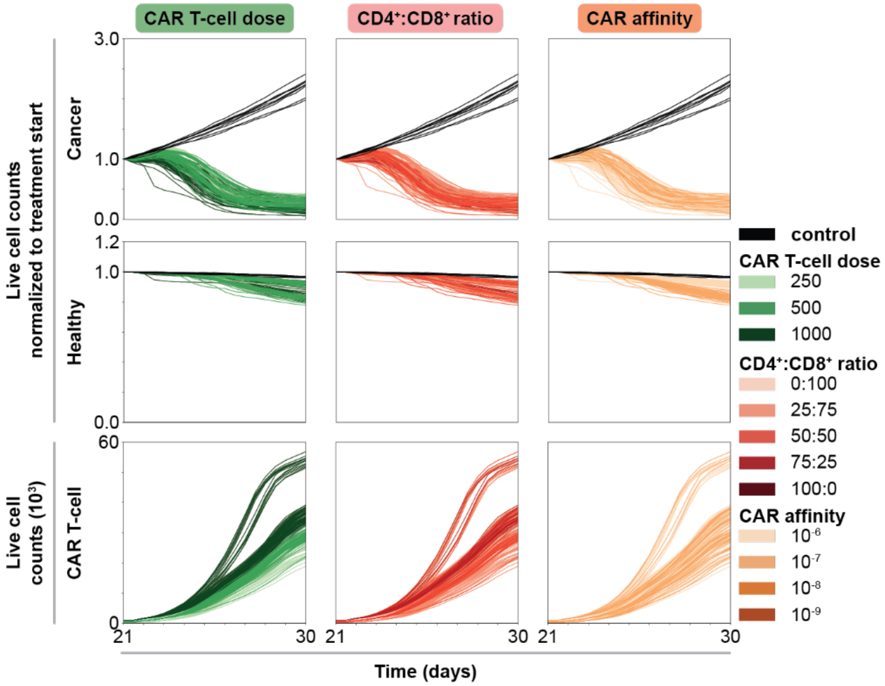

**SUPPLEMENTARY FIGURE 12. Dynamic and spatial outcomes for selected promising treatment combinations in `tissue`.** Normalized live cell counts over time of untreated (black) and treated conditions (graded hues), normalized to live cell count at start of treatment (t = 21 d), for all simulations, colored by CAR T-cell dose reported in units of CAR T-cells, $CD4^+$:$CD8^+$ ratio, and CAR affinity reported in units of M. Note: not all combinations of features were simulated, see **Supplementary Table 6** for combinations tested.

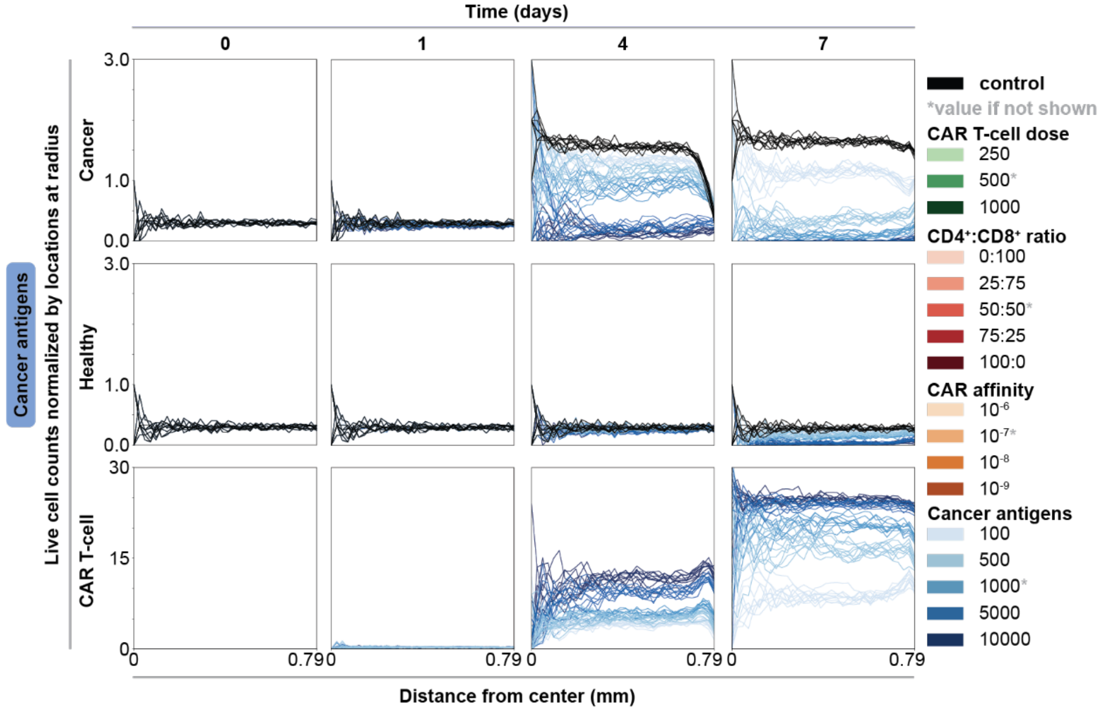

**SUPPLMENTAL FIGURE 13. Spatial dynamics for each cell type in realistic co-culture.** Normalized live cell counts over time (t = 0, 1, 4, and 7 d shown) for untreated (black) and treated conditions (graded hues), normalized to locations per radius, for all simulations. Data are colored by cancer antigens while all other features are held constant at indicated intermediate value (indicated by asterisk, CAR T-cell dose–500 CAR T-cells, $CD4^+$:$CD8^+$ ratio–50:50, CAR affinity–$10^{-7}$ M). Cancer antigens reported in units of antigens/cell. The columns indicate the timepoint in the simulation (day), while the rows indicate cell type plotted, and the x-axis for each plot shows the distance from the center.

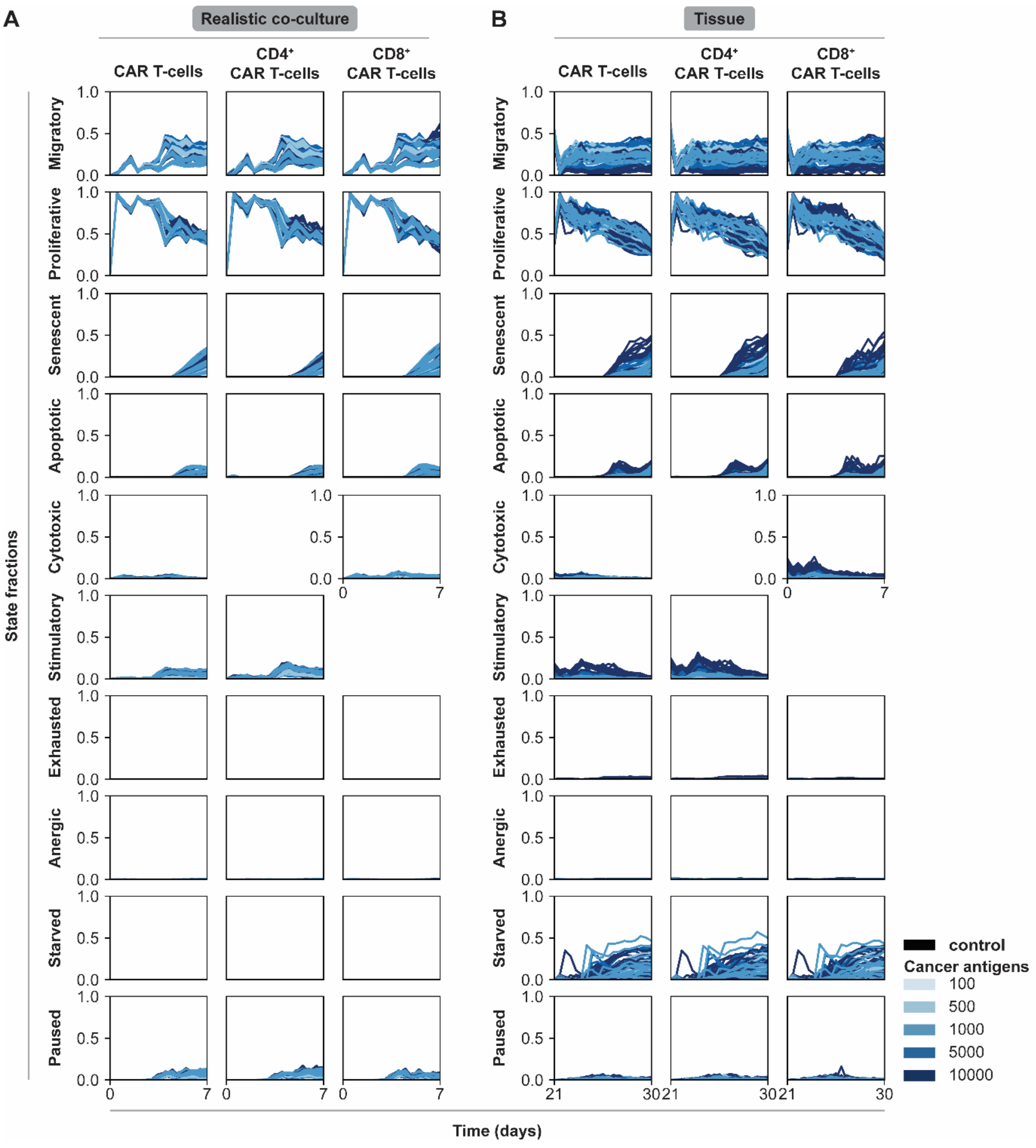

**SUPPLEMENTARY FIGURE 14. CAR T-cells that share locations with cancer cells show similar dynamics in realistic co-culture `dish` and `tissue`.** CAR T-cell state fractions for CAR T-cells that share a location with at least one cancer cell in selected effective treatment conditions in both **(A)** realistic co-culture `dish` and **(B)** `tissue` (after treatment start) over time. Colors represent varying cancer antigen levels, which are reported in units of antigens/cell. Each column shows the cell type being plotted, while rows show the cell state fraction being plotted. Note: not all combinations of features were simulated, see **Supplementary Table 6** for combinations tested.

#### 3.2 Supplementary Tables

**SUPPLEMENTARY TABLE 1. CARCADE parameter names, descriptions, values, and sources/derivations.**

| Parameter | Code | Value | Citation/Derivation |
|---|---|---|---|
| IL-2 Diffusivity in Blood | DIFFUSIVITY_IL2 | 10.0 um$^2$/s | 1 x 10$^{-7}$ cm$^2$/s (Gong et al., 2017)<br><br>36,000 um$^2$/hr (Busse et al., 2010)<br><br>Both convert to 10 um$^2$/s |
| Initial IL-2 Concentration | CONCENTRATION_IL2 | 0 molecules/ cm$^3$ | Assumption |
| Maximum damage value at which T-cells can spawn next to in source or pattern source | MAX_DAMAGE_SEED | 1e7 (unitless) | Arbitrarily high value such that CAR T-cells can spawn at any vasculature point |
| Minimum radius value at which T-cells can spawn next to in graph source | MIN_RADIUS_SEED | 2 um | Minimum diameter of vasculature edge is 4 um in ARCADE (Yu and Bagheri, 2021) so 2 um is minimum that the radius will ever be, so at this value CAR T-cells can spawn everywhere |
| CAR T-cell life span average | DEATH_AGE_AVG_T | 6 weeks<br>60480 min | 6 weeks (Borghans and Ribeiro, 2017) |
| CAR T-cell minimum age when T-cells are spawned | T_CELL_AGE_MIN | 0 min | Assumption |
| CAR T-cell maximum age when T-cells are spawned | T_CELL_AGE_MIN | 1 week<br>10080 min | Assumption |
| T-cell DNA synthesis time distribution average | SYNTHESIS_TIME_T | 360 min<br>6 h | Same as tissue cell DNA synthesis time chosen in ARCADE (Yu and Bagheri, 2020)<br><br>T-cell division time has been found to have a wide range of reported values, including 2 h (Yoon et al., 2010); 4-6 h (Altman and Dang, 2012); and 13.4 +/- 5.4 and 14.3 +/- 4.4 h, but also slow cell cycle times of 24 h or can also be as low or less than as 600 min (10 h) (Kinjyo et al., 2015)<br><br>Additionally, a range of other values have been calculated by other models, such as 8 h for CD8$^+$s and 11 h for CD4$^+$s (De Boer et al., 2003) and being capable of 4-5 divisions in 99 h with high IL-2 saturation giving 19-24 h per |

| | | | |
|---|---|---|---|
| | | | division (Deenick et al., 2003); or used in other models, such as 8 h (Gong et al., 2017) |
| T-cell DNA synthesis time distribution range | SYNTHESIS_TIME_T_RANGE | 0 min | Same as tissue cell DNA synthesis time chosen in ARCADE (Yu and Bagheri, 2020) |
| Duration a CAR T-cell stays bound to a cell upon binding to antigen distribution average | BOUND_TIME | 360 min<br>6 h | Assumption, chose 1/3rd of apoptosis time of cancer cells as dictated by ARCADE (Yu and Bagheri, 2020) |
| Duration a CAR T-cell stays bound to a cell upon binding to antigen distribution range | BOUND_TIME_RANGE | 0 min | Assumption |
| CAR T-cell volume average | T_CELL_VOL_AVG | 175 $um^3$ | ~140 fL (noting that activated T cells are larger) (Jacobs et al., 2008)<br><br>206 +/- 14.4 fl (Kuse et al., 1985)<br><br>Approximate average between two sources (fl = $um^3$) |
| CAR T-cell volume range | T_CELL_VOL_RANGE | 10.0 $um^3$ | ~10% of T-cell volume (similar approach as tissue cell volume range in ARCADE) (Yu and Bagheri, 2020) |
| Energy requirement of an activated CAR T-cell to perform effector functions after antigen-induced activation | ACTIVE_ENERGY | 0.002 fmol ATP/$um^3$ cell/min | Estimated |
| Increase in fraction of glucose used in cell mass production due to antigen-induced activation | FRAC_MASS_ACTIVE | 0.25 (unitless) | Estimated; evidence of parameter existence (Wang et al., 2011) |
| Maximum increase in overall metabolic preference for glycolysis due to IL-2 feedback | META_PREF_IL2 | 0.1 (unitless) | Estimated; original parameter META_PREF in ARCADE (Yu and Bagheri, 2020) |
| Increase in overall metabolic preference for glycolysis due to antigen-induced activation | META_PREF_ACTIVE | 0.3 (unitless) | Estimated; original parameter META_PREF in ARCADE (Yu and Bagheri, 2020) |

| | | | |
|---|---|---|---|
| Maximum increase in CAR T-cell glucose uptake rate due to impact of IL-2 on metabolism | `GLUC_UPTAKE_RATE_IL2` | 0.56 fmol glucose/ $um^2$ cell/min/M glucose | Estimated; half of basal GLUCOSE_UPTAKE_RATE as defined by ARCADE (Yu and Bagheri, 2020) |
| Increase in CAR T-cell glucose uptake rate due to antigen-induced activation | `GLUC_UPTAKE_RATE_ACTIVE` | 3.78 fmol glucose/ $um^2$ cell/min/M glucose | ~5-fold increase in glucose uptake rate in activated T-cells (used 4.375-fold because Figure 3A of reference looks like it goes from 400 before activation and 1750 after) (Frauwirth et al., 2002)<br><br>10-fold increase in glucose uptake rate in activated T-cells (Jacobs et al., 2008)<br><br>Used first reference and original value of GLUCOSE_UPTAKE_RATE (1.12) to calculate full fold increase (which includes impact of antigen-induced activation and IL-2):<br><br>1.12 x 4.375 = 4.9 fmol glucose/$um^2$ cell/min/M glucose<br><br>Subtract from total the increase from IL-2 independently to calculate increase from antigen-induced activation independently: 4.9 mol glucose/$um^2$ cell/min/M glucose – 0.56 fmol glucose/$um^2$ cell/min/M glucose = 3.78 fmol glucose/$um^2$ cell/min/M glucose |
| Time required for CAR T-cell to sustain antigen-induced activation before switching increasing metabolic preference (represents time delay in protein production to alter metabolism) | `META_SWITCH_DELAY` | 60 min | Estimated; source noted "metabolic changes in T cells after activation occur extremely rapidly, as changes in calcium flux and lactate production can be observed only minutes after ligand binding" (Altman and Dang, 2012) |
| Distance above a cell that cells can sense molecules and proteins | `SHELL_THICKNESS` | 2 um | (Gherbi et al., 2018) |
| Total IL-2 receptors per CAR T-cell (same both before and after IL-2Rα, as this is not limiting receptor part) | `IL2_RECEPTORS` | 2 x $10^3$ receptors/ cell | Reference pg. 30, 1993 (Lauffenburger and Linderman, 1993) |

| | | | |
|---|---|---|---|
| IL-2 binding rate to IL-2Rβγ$_c$ complex ($k_{on}$) | `IL2_BINDING_ON_RATE_MIN` | 3.8193 x $10^{-2}$ $um^3$ $molecules^{-1}$ $min^{-1}$ | 2.3 x $10^7$ $M^{-1}min^{-1}$<br>Reference pg. 30, 1993 (Lauffenburger and Linderman, 1993)<br><br>2.3 x $10^7$ $M^{-1}min^{-1}$ = 2.3 x $10^7$ L $mol^{-1}$ $min^{-1}$ = 2.3 x $10^{22}$ $um^3$ $mol^{-1}$ $min^{-1}$<br><br>Divide by Avogadro's number to get from moles to molecules:<br>2.3x$10^{22}$ $um^3$ $mol^{-1}$ $min^{-1}$ / 6.022 x$10^{22}$ molecules $mol^{-1}$ = 0.038193 $um^3$ $molecules^{-1}$ $min^{-1}$ |
| IL-2 binding rate to IL-2Rβγ$_c$α complex ($k_{on}$) | `IL2_BINDING_ON_RATE_MAX` | 3.155 $um^3$ $molecules^{-1}$ $min^{-1}$ | 1.9 x $10^9$ $M^{-1}min^{-1}$<br>Reference pg. 30, 1993 (Lauffenburger and Linderman, 1993)<br><br>111.6 $nM^{-1}h^{-1}$ (1.86 x $10^9$ $M^{-1}min^{-1}$) (Busse et al., 2010)<br><br>1.9 x $10^9$ $M^{-1}min^{-1}$ = 1.9 x $10^9$ L $mol^{-1}$ $min^{-1}$ = 1.9 x $10^{24}$ $um^3$ $mol^{-1}$ $min^{-1}$<br><br>Divide by Avogadro's number to get from moles to molecules:<br>1.9 x $10^{24}$ $um^3$ $mol^{-1}$ $min^{-1}$<br>/ 6.022 x$10^{22}$ molecules $mol^{-1}$ = 3.115 $um^3$ $molecules^{-1}$ $min^{-1}$ |
| Binding off rate of IL-2 to IL-2Rβγ$_c$ complex and IL-2Rβγ$_c$α complex ($k_{off}$) | `IL2_BINDING_OFF_RATE` | 0.015 $min^{-1}$ | 0.015 $min^{-1}$ for heavy chain; 0.014 $min^{-1}$ for heterodimer<br>Reference pg. 30, 1993 (Lauffenburger and Linderman, 1993)<br><br>0.83 $hr^{-1}$ (0.01388 $min^{-1}$) (Busse et al., 2010) |
| Rate of conversion of IL-2Rβγ$_c$ two-chain complex to IL-2Rβγ$_c$α, independent of if IL-2 bound to receptor complex (parameter found within Inflammation module class) | `K_CONVERT` | 1 x$10^{-3}$ $s^{-1}$ | Estimated |
| Rate of recycling of any IL-2R complexes bound to IL-2 or IL-2Rβγ$_c$α complex back to IL-2Rβγ$_c$ (parameter found within Inflammation module class) | `K_REC` | 1 x$10^{-5}$ $s^{-1}$ | Estimated |

| | | | |
|---|---|---|---|
| Maximum production rate of IL-2 by $CD4^+$ CAR T-cells due to IL-2 feedback | IL2_PROD_RATE_IL2 | 16.62 molecules IL-2/cell/min | 1000 molecules/cell-min (Busse et al., 2010) |
| Maximum production rate of IL-2 by $CD4^+$ T-cells due to antigen-induced activation | IL2_PROD_RATE_ACTIVE | 293.27 molecules IL-2/cell/min | $4.87 \times 10^{-16}$ umol/cell-min (Busse et al., 2010)<br><br>$4.87 \times 10^{-16}$ umol/cell-min * 1 mol/ $10^6$ umol * $6.022 \times 10^{23}$ molecules/mol = 293.27 molecules IL-2/cell/min<br><br>Note: production rate is 0 molecules IL-2/cell/min for inactivated cells (Jacobs et al., 2008; Guedan et al., 2018) |
| Maximum production rate of IL-2 by $CD4^+$ CAR T-cells due to antigen-induced activation and IL-2 feedback | Not a formal parameter; more for reference/note | N/A | $10^{-5}$ ng/cell-day (calculated to be 1.79 fmol IL-2/cell-s Using IL-2 is 15.5kDa; winds up being $4.48 \times 10^{-16}$ umol/cell-min) (Robertson-Tessi et al., 2012)<br><br>Estimated $5.6 \times 10^{-17}$ umol/min-cell (calculated from 1 ng IL-2/ml produced by 4.5 x $10^5$ cells in a 48 well plate assumed to have at most 500 ul in it and MW of IL2 taken to be 15.5 kDa – but can be 1-40 ng/ml; this equates to $1.12 \times 10^{-16}$ umol/$cm^3$-min-cell) (if you add together $4.87 \times 10^{-16}$ and $2.76 \times 10^{-17}$ you get $5.14 \times 10^{-17}$ which is about this) (Guedan et al., 2018)<br><br>Calculated to be $4.87 \times 10^{-16}$ umol/cell-min (from 17600 IL-2 molecules/cell-h in Gong ABM) (Gong et al., 2017)<br><br>23400 molecules/cell-hr in first 8 h of antigen stimulation ($6.47 \times 10^{-16}$ umol/cell-min) and 6000 molecules/cell-hr ($1.66 \times 10^{-16}$ umol/cell-min) between 8 and 12 h (Busse et al., 2010) (noting that they estimated this from another study) |
| Time CAR T-cells must sustain antigen binding to produce IL-2 mRNA | IL2_SYNTHESIS_DELAY | 180 min | 2-4 h (Iwashima, 2003)<br>(chose midpoint of 3 h = 180 min) |
| Time required for CAR T-cells to maintain bound contact with target antigen before activation signal | GRANZ_SYNTHESIS_DELAY | 15 min | Estimated |

| | | | |
|---|---|---|---|
| induces granzyme production | | | |
| Moles of granzyme produced by $CD8^+$ CAR T-cells per mol IL-2 bound (parameter found within InflammationCD8 module class) | `GRANZ_PER_IL2` | 0.005 mol granzyme/ mol IL-2 | Estimated |
| Moles granzyme required by $CD8^+$ CAR T-cells to kill target cell (parameter found indirectly KillerCARTHelper helper) | | 1 mole | Estimated |
| CAR T-cell division potential | `DIVISION_POTENTIAL_T` | 10 divisions/cell | 9 divisions/cell (Kinjyo et al., 2015)<br><br>Average of 7 divisions/cell with range 4-10 divisons/cell for $CD4^+$ T-cells; Range of 15-19 divisions/cell for $CD8^+$ T-cells (Obst, 2015) (note: used 17 as middle of 15-19 range given for $CD8^+$ T-cells from provided citation)<br><br>8 divisions per T-cell (Gong et al., 2017)<br><br>Average of 9, 7, 17, and 8 = 10.25, rounded down to 10 |
| CAR T-cell fraction that becomes proliferative (as opposed to migratory) if not activated via antigen-induced activation | `PROLI_FRAC` | 0.5 (unitless) | Estimated |
| CAR T-cell fraction that becomes exhausted vs apoptotic | `EXHAU_FRAC` | 0.5 (unitless) | Estimated |
| CAR T-cell fraction that becomes anergic vs apoptotic | `ANERG_FRAC` | 0.5 (unitless) | Estimated |
| Maximum number of cells a CAR T-cell could attempt to make contact with per time step | `SEARCH_ABILITY` | 1 cell | Estimated |

| | | | |
|---|---|---|---|
| Number of times a CAR T-cell can bind to a target before becoming exhausted | `MAX_ANTIGEN_BINDING` | 10 binding events | Estimated |
| Average number of CARs on a CAR T-cell's surface | `CARS` | 50000 receptors/cell | >50000 but varies receptors/cell (Harris and Kranz, 2016)<br><br>Noted that value is the same on both 41BB and CD28 CAR T-cells (Salter et al., 2018) |
| Average number of CAR antigens on a healthy tissue cell | `CAR_ANTIGENS_HEALTHY` | 100 antigens/cell | minimum of 100 antigens/cell (Harris and Kranz, 2016)<br><br>1000 antigens/cell (Robertson et al., 1996; Stone et al., 2012; Liu et al., 2015; Harris and Kranz, 2016)<br><br>10000 (maximum 1 million) antigens/cell (Robertson et al., 1996; Stone et al., 2012; Liu et al., 2015)<br><br>1000-7500 antigens/cell (Arcangeli et al., 2017) |
| Average number of CAR antigens on a cancerous tissue cell<br>(Should be higher on cancer cells than healthy cells) | `CAR_ANTIGENS_CANCER` | 1000 antigens/cell | minimum of 100 antigens/cell (Harris and Kranz, 2016)<br><br>1000 antigens/cell (Robertson et al., 1996; Stone et al., 2012; Liu et al., 2015; Harris and Kranz, 2016)<br><br>10000 (maximum 1 million) antigens/cell (Robertson et al., 1996; Stone et al., 2012; Liu et al., 2015)<br><br>1000-7500 antigens/cell (Arcangeli et al., 2017) |
| Average number of self-receptors (PD1s) on a CAR T-cell | `SELF_RECEPTORS` | 150 surface ligands/cell | 150 surface ligands/cell before antigen-induced activation, 9000 after antigen-induced activation (Cheng et al., 2013) |
| Average number of self-receptors (PDL1s) on a tissue cell | `SELF_TARGETS` | 3,600 surface ligands/cell | MDA-MB-231 breast cancer cells: 47,700 +/- 2,900 (89.4% $PDL1^{+}$)<br>SK-Br-3 breast cancer cells: 2,000 +/- 100 (2.9% $PDL1^{+}$)<br>SUM149 breast cancer cells: 3,600 +/- 400 for (8.9% $PDL1^{+}$)<br>(Heskamp et al., 2015) |
| Affinity of CAR for target antigen | `CAR_AFFINITY` | $1 \times 10^{-7}$ M | Weak = 1000 nmol/L ($10^{-6}$ M)<br>Strong = 0.1 nmol/L ($10^{-10}$ M)<br>(Hegde et al., 2013; Harris and Kranz, 2016; Hegde et al., 2016) |

| | | | |
|---|---|---|---|
| Fitting factor α in CAR binding function | `CAR_ALPHA` | 3 (unitless) | Estimated |
| Fitting factor β in CAR binding function | `CAR_BETA` | 0.01 antigens/M | Estimated |
| Affinity of self-receptor for self (based on PD1 for PDL1) | `SELF_RECEPTOR_AFFINITY` | 7.8 x $10^{-6}$ M | 7.8 uM (at 37°C) (Cheng et al., 2013) Converted to 7.8 x $10^{-6}$ M |
| Fitting factor α in self-receptor (PD1) binding function | `SELF_ALPHA` | 3 (unitless) | Estimated |
| Fitting factor β in self-receptor (PD1) binding function | `SELF_BETA` | 0.02 antigens/M | Estimated |
| Fraction of cell surface contacting a bound cell during a binding event (γ) | `CONTACT_FRAC` | 0.2 (unitless) | Estimated |

#### SUPPLEMENTARY TABLE 2. Tuned feature values for monoculture `dish` simulations.

| Modified Features | Simulated Values |
|---|---|
| Dose of CAR T-Cells (CAR T-cell dose) | 250, 500, 1000 |
| $CD4^+$:$CD8^+$ CAR T-Cell Ratio ($CD4^+$:$CD8^+$ratio) | 100:0, 75:25, 50:50, 25:75, 0:100 |
| CAR-Antigen Affinity (CAR affinity, units: M) | $1x10^{-6}$, $1x10^{-7}$, $1x10^{-8}$, $1x10^{-9}$ |
| Antigens Per Cancer Cell (cancer antigens) | 100, 500, 1000, 5000, 10000 |
| Total combinations | 300 |
| Total simulations | 3000 |

#### SUPPLEMENTARY TABLE 3. Tuned feature values for co-culture `dish` simulations.

| Modified Parameters | Simulated Values |
|---|---|
| Dose of CAR T-Cells (CAR T-cell dose) | 250, 500, 1000 |
| $CD4^+$:$CD8^+$ CAR T-Cell Ratio ($CD4^+$:$CD8^+$ratio) | 100:0, 75:25, 50:50, 25:75, 0:100 |
| CAR-Antigen Affinity (CAR affinity, units: M) | $1x10^{-6}$, $1x10^{-7}$, $1x10^{-8}$, $1x10^{-9}$ |
| Antigens Per Cancer Cell (cancer antigens) | 100, 500, 1000, 5000, 10000 |
| Antigens Per Healthy Cell (healthy antigens) | 0 (ideal), 100 (realistic) |
| Total combinations | 600 |
| Total simulations | 6000 |

**SUPPLEMENTARY TABLE 4. Input options used to run monoculture `dish` simulations.** For clarity, wrapping `<set>` tags, `<series>` name attributes, `<profilers>` simulation tags, `<globals>` environment tags are not shown. All sets use growth, parameter, and lysis profilers, each using interval 720. All cancer and healthy cells use default modules. For each set, simulations were run for every combination of bold options grouped by square brackets and separated by pipes.

<table>
<tr><th>Set</th><th>Input</th></tr>
<tr><td>Untreated</td><td><pre>
<series start="0" end="10" days="7">
   <simulation type="growth">
   </simulation>
   <agents initialization="2000" plate="dish">
      <populations>
         <population type="C" fraction="1.0">
            <variables>
               <variable id="CAR_ANTIGENS_CANCER" value="1000"/>
            </variables>
         </population>
         <population type="H" fraction="0">
            <variables>
               <variable id="CAR_ANTIGENS_HEALTHY" value="0"/>
            </variables>
         </population>
         <population type="4" fraction="0.0">
         </population>
         <population type="8" fraction="0.0">
         </population>
      </populations>
      <helpers>
         <helper type="treat" delay="10" dose="0"/>
      </helpers>
   </agents>
   <environment coordinate="hex">
      <components>
         <component type="sites" class="source">
            <specifications>
               <specification id="X_SPACING" value="*" />
               <specification id="Y_SPACING" value="*" />
               <specification id="SOURCE_DAMAGE" value="0.0" />
            </specifications>
         </component>
      </components>
   </environment>
</series>
</pre></td></tr>
<tr><td>Treated</td><td><pre>
<series start="0" end="10" days="7">
   <simulation type="growth">
   </simulation>
   <agents initialization="2000" plate="dish">
      <populations>
         <population type="C" fraction="1.0">
            <variables>
               <variable id="CAR_ANTIGENS_CANCER"
                    value="[100|500|1000|5000|10000]"/>
            </variables>
         </population>
         <population type="H" fraction="0">
            <variables>
</pre></td></tr>
</table>

```
                <variable id="CAR_ANTIGENS_HEALTHY" value="0" />
            </variables>
        </population>
        <population type="4" fraction="0.0">
            <variables>
                <variable id="CAR_AFFINITY" value="[1e-6|1e-7|
                1e-8|1e-9]" />
            </variables>
        </population>
        <population type="8" fraction="0.0">
            <variables>
                <variable id="CAR_AFFINITY" value="[1e-6|1e-7|
                1e-8|1e-9]" />
            </variables>
        </population>
    </populations>
    <helpers>
        <helper type="treat" delay="10" dose="[250|500|1000]"
            ratio="[1:0|0.75:0.25|0.5:0.5|0.25:0.75|0:1]" />
    </helpers>
</agents>
<environment coordinate="hex">
    <components>
        <component type="sites" class="source">
            <specifications>
                <specification id="X_SPACING" value="*" />
                <specification id="Y_SPACING" value="*" />
                 <specification id="SOURCE_DAMAGE" value="0.0" />
            </specifications>
        </component>
    </components>
</environment>
</series>
```

**SUPPLEMENTARY TABLE 5. Input options used to run co-coculture dish simulations.** For clarity, wrapping <set> tags, <series> name attributes, <profilers> simulation tags, <globals> environment tags are not shown. All sets use growth, parameter, and lysis profilers, each using interval 720. All cancer and healthy cells use default modules. For each set, simulations were run for every combination of bold options grouped by square brackets and separated by pipes. Simulations that where the healthy population had a value of 0 for CAR_ANTIGENS_HEALTHY are part of the ideal co-culture data, while simulations that where the healthy population had a value of 0 for CAR_ANTIGENS_HEALTHY are part of the realistic co-culture data.

| Set | Input |
|---|---|

**Untreated**

```
<series start="0" end="10" days="7">
   <simulation type="growth">
   </simulation>
   <agents initialization="2000" plate="dish">
      <populations>
         <population type="C" fraction="0.5">
            <variables>
               <variable id="CAR_ANTIGENS_CANCER" value="1000"/>
            </variables>
         </population>
         <population type="H" fraction="0.5">
             <variables>
               <variable id="CAR_ANTIGENS_HEALTHY" value="100"/>
            </variables>
         </population>
            <population type="4" fraction="0.0">
               <variables>
                  <variable id="CAR_AFFINITY" />
               </variables>
            </population>
            <population type="8" fraction="0.0">
               <variables>
                  <variable id="CAR_AFFINITY" />
               </variables>
            </population>
      </populations>
      <helpers>
         <helper type="treat" delay="10" dose="0"/>
      </helpers>
   </agents>
   <environment coordinate="hex">
      <components>
         <component type="sites" class="source">
            <specifications>
               <specification id="X_SPACING" value="*" />
               <specification id="Y_SPACING" value="*" />
               <specification id="SOURCE_DAMAGE" value="0.0" />
            </specifications>
         </component>
      </components>
   </environment>
</series>
```

**Treated**

```
<series start="0" end="10" days="7">
   <simulation type="growth">
   </simulation>
```

```
    <agents initialization="2000" plate="dish">
      <populations>
        <population type="C" fraction="0.5">
          <variables>
            <variable id="CAR_ANTIGENS_CANCER"
                 value="[100|500|1000|5000|10000]"/>
          </variables>
        </population>
        <population type="H" fraction="0.5">
          <variables>
            <variable id="CAR_ANTIGENS_HEALTHY" value="[0|100]" />
          </variables>
        </population>
        <population type="4" fraction="0.0">
          <variables>
            <variable id="CAR_AFFINITY" value="[1e-6|1e-7|
            1e-8|1e-9]" />
          </variables>
        </population>
        <population type="8" fraction="0.0">
          <variables>
            <variable id="CAR_AFFINITY" value="[1e-6|1e-7|
            1e-8|1e-9]" />
          </variables>
        </population>
      </populations>
      <helpers>
        <helper type="treat" delay="10" dose="[250|500|1000]"
          ratio="[1:0|0.75:0.25|0.5:0.5|0.25:0.75|0:1]" />
      </helpers>
    </agents>
    <environment coordinate="hex">
      <components>
        <component type="sites" class="source">
          <specifications>
            <specification id="X_SPACING" value="*" />
            <specification id="Y_SPACING" value="*" />
             <specification id="SOURCE_DAMAGE" value="0.0" />
          </specifications>
        </component>
      </components>
    </environment>
  </series>
```

**SUPPLEMENTARY TABLE 6. Reference information from cited papers' *in vitro* experiments shown in Figure 2G.**

| Citation | Reference Figure Containing Data | CAR Construct Used in Study | E:T Ratio | Notes |
|---|---|---|---|---|
| Arcangeli 2017 | Figure 4 | CD123-CD28-OX40-CD3ζ | 5:1 | Short-Term Assay (4 h); co-culture with low antigen expressing cells |
| Caruso 2015 | Figure 1 for 1e6 and 0, other data from Figure 4 | EGFR-CD28-CD3ζ | 5:1 | |
| Chmielewski 2004 | Figure 4 | ErbB2-Fc-CD3ζ | 1:1 | Antigen levels are MFI; They provided viability data so convereted to % lysis by using: % Kill = 1- Viability %; Used E:T ratio closed to 1:1 |
| Ghorashian 2019 | Figure 1 | CD19-41BB-CD3ζ | 6.4:1 | |
| Hernandez-Lopez | Figure 2A upper plot | HER2-41BB-CD3ζ | not found | |
| Liu 2015 | Figure 2 | ErbB2-41BB-CD3ζ | 1:1 | Antigen level = ErbB2 RNA ug |
| Watanabe 2014 | Figure 2C | CD20-CD28-CD3ζ | 1:1 | |

**SUPPLEMENTARY TABLE 7. Effective treatments identified in co-culture `dish` simulations**. Simulations were averaged across replicates and ordered from highest to lowest difference metric.

| CAR T-Cell Dose | $CD4^+$:$CD8^+$ T-Cell Ratio | CAR Affinity (M) | Antigens Cancer | Antigens Healthy | Normalized Cancer Cell Count | Normalized Healthy Cell Count | Normalized T-cell Count | Difference Metric Value |
|---|---|---|---|---|---|---|---|---|
| 1000 | 0.25 | 1E-06 | 10000 | 100 | 0.03 | 0.75 | 73.09 | 0.72 |
| 500 | 0.25 | 1E-06 | 10000 | 100 | 0.13 | 0.83 | 141.33 | 0.69 |
| 1000 | 0.5 | 1E-06 | 10000 | 100 | 0.12 | 0.80 | 70.68 | 0.68 |
| 1000 | 0.25 | 1E-06 | 5000 | 100 | 0.36 | 0.83 | 65.38 | 0.48 |
| 500 | 0.5 | 1E-06 | 10000 | 100 | 0.40 | 0.86 | 131.76 | 0.46 |
| 1000 | 0.75 | 1E-06 | 10000 | 100 | 0.61 | 0.86 | 61.97 | 0.25 |
| 250 | 0.25 | 1E-06 | 10000 | 100 | 0.80 | 0.92 | 231.88 | 0.12 |
| 1000 | 0.5 | 1E-06 | 5000 | 100 | 0.75 | 0.87 | 59.82 | 0.12 |
| 500 | 0.25 | 1E-06 | 5000 | 100 | 0.78 | 0.88 | 118.79 | 0.10 |
| 500 | 0.5 | 1E-07 | 1000 | 100 | 0.42 | 0.50 | 137.30 | 0.09 |
| 1000 | 0.75 | 1E-07 | 1000 | 100 | 0.59 | 0.56 | 64.57 | -0.03 |
| 1000 | 0.5 | 1E-07 | 500 | 100 | 0.68 | 0.52 | 63.45 | -0.16 |
| 250 | 0.25 | 1E-07 | 1000 | 100 | 0.83 | 0.63 | 240.64 | -0.20 |
| 500 | 0.25 | 1E-07 | 500 | 100 | 0.75 | 0.54 | 126.98 | -0.21 |

**SUPPLEMENTARY TABLE 8. Input options used to run `tissue` simulations.** For clarity, wrapping `<set>` tags, `<series>` name attributes, `<checkpoint>` name and path attributes, `<profilers>` simulation tags, and `<globals>` environment tags are not shown. All untreated and treated sets use growth, parameter, and lysis profilers, each using interval 720, except the untreated set used to produce the graph in which a graph profiler was used instead of a parameter profiler. All cancer and healthy cells use default modules. For the treated set, simulations were run for each of the 14 treatment conditions shown in **Supplementary Table 5**. Variable values shown are chosen in the listed order of the values in square brackets and separated by pipes, where a single simulation uses one value from each variable list such that all values are at the same list position. For example, if parameter A has list [$\mathbf{a}_1$ | $\mathbf{a}_2$] and parameter B has list [$\mathbf{b}_1$ | $\mathbf{b}_2$], then the first simulation would use parameter values $a_1$ and $b_1$, while the second simulation would use parameter values $a_2$ and $b_2$.

**Set** — **Input**

Graph

```
<series start="0" end="1" days="1">
   <simulation type="growth">
      <checkpoints>
         <checkpoint type="graph" class="save" day="0" />
      </checkpoints>
   </simulation>
   <agents initialization="0" />
   <environment coordinate="hex">
      <components>
         <component type="sites" class="graph" complexity="simple">
            <specifications>
               <specification id="GRAPH_LAYOUT" value="S" />
               <specification id="ROOTS_LEFT" value="50A" />
               <specification id="ROOTS_RIGHT" value="50A" />
               <specification id="ROOTS_TOP" value="50V" />
               <specification id="ROOTS_BOTTOM" value="50V" />
            </specifications>
         </component>
      </components>
   </environment>
</series>
```

Untreated

```
<series start="0" end="10" days="31">
   <simulation type="growth">
      <checkpoints>
         <checkpoint type="graph" class="load" day="0" />
      </checkpoints>
    </simulation>
    <agents initialization="FULL">
       <populations>
          <population type="C" fraction="0.0">
             <variables>
                <variable id="CAR_ANTIGENS_CANCER" value="1000"/>
             </variables>
          </population>
          <population type="H" fraction="1.0">
             <variables>
                <variable id="CAR_ANTIGENS_HEALTHY" value="100" />
             </variables>
          </population>
          <population type="4" fraction="0.0">
             <variables>
                <variable id="CAR_AFFINITY" />
             </variables>
```

```
          </population>
          <population type="8" fraction="0.0">
             <variables>
                <variable id="CAR_AFFINITY" />
             </variables>
          </population>
       </populations>
       <helpers>
          <helper type="insert" delay="1440" populations="0"
          bounds="0.05" />
          <helper type="treat" delay="31680" dose="0" />
       </helpers>
   </agents>
   <environment coordinate="hex">
      <components>
         <component type="sites" class="graph" layout="(S)"
         left="(50A)" right="(50A)" top="(50V)" bottom="(50V)" />
         <component type="remodel" interval="60" />
         <component type="degrade" interval="1" />
      </components>
   </environment>
</series>
```

Untreated (generate graph images for Figure 1)

```
<series start="0" end="10" days="31">
   <simulation type="growth">
      <checkpoints>
         <checkpoint type="graph" class="load" day="0" />
      </checkpoints>
    </simulation>
    <agents initialization="FULL">
       <populations>
          <population type="C" fraction="0.0">
             <variables>
                <variable id="CAR_ANTIGENS_CANCER" value="1000"/>
             </variables>
          </population>
          <population type="H" fraction="1.0">
             <variables>
                <variable id="CAR_ANTIGENS_HEALTHY" value="100" />
             </variables>
          </population>
          <population type="4" fraction="0.0">
             <variables>
                <variable id="CAR_AFFINITY" />
             </variables>
          </population>
          <population type="8" fraction="0.0">
             <variables>
                <variable id="CAR_AFFINITY" />
             </variables>
          </population>
       </populations>
       <helpers>
          <helper type="insert" delay="1440" populations="0"
          bounds="0.05" />
          <helper type="treat" delay="31680" dose="0" />
       </helpers>
   </agents>
```

```
      <environment coordinate="hex">
         <components>
            <component type="sites" class="graph" layout="(S)"
            left="(50A)" right="(50A)" top="(50V)" bottom="(50V)" />
            <component type="remodel" interval="60" />
            <component type="degrade" interval="1" />
         </components>
      </environment>
   </series>
```

Treated

```
   <series start="0" end="10" days="31">
      <simulation type="growth">
         <checkpoints>
            <checkpoint type="graph" class="load" day="0" />
               </checkpoints>
           </simulation>
      <agents initialization="FULL">
         <populations>
            <population type="C" fraction="0.0">
               <variables>
                  <variable id="CAR_ANTIGENS_CANCER"
                  value="[10000|10000|10000|5000|10000|10000|10000|
                  5000|5000|1000|1000|500|1000|500]"/>
               </variables>
            </population>
            <population type="H" fraction="1.0">
               <variables>
                  <variable id="CAR_ANTIGENS_HEALTHY" value="100" />
               </variables>
            </population>
            <population type="4" fraction="0.0">
               <variables>
                  <variable id="CAR_AFFINITY" value="[1e-6|1e-6|1e-6|
                  1e-6|1e-6|1e-6|1e-6|1e-6|1e-6|1e-7|1e-7|1e-7|1e-7|
                  1e-7]" />
               </variables>
            </population>
            <population type="8" fraction="0.0">
               <variables>
                  <variable id="CAR_AFFINITY" value="[1e-6|1e-6|1e-6|
                  1e-6|1e-6|1e-6|1e-6|1e-6|1e-6|1e-7|1e-7|1e-7|1e-7|
                  1e-7]" />
               </variables>
            </population>
         </populations>
         <helpers>
            <helper type="insert" delay="1440" populations="0"
            bounds="0.05" />
            <helper type="treat" delay="31680" dose="[1000|500|1000|
            1000|500|1000|250|1000|500|500|1000|1000|250|500]"
            ratio="[0.25:0.75|0.25:0.75|0.5:0.5|0.25:0.75|0.5:0.5|
            0.75:0.25|0.25:0.75|0.5:0.5| 0.25:0.75|0.5:0.5|0.75:0.25|
            0.5:0.5|0.25:0.75|0.25:0.75]" />
         </helpers>
      </agents>
      <environment coordinate="hex">
         <components>
            <component type="sites" class="graph" layout="(S)"
            left="(50A)"
               right="(50A)" top="(50V)" bottom="(50V)" />
```

```
                  <component type="remodel" interval="60" />
                  <component type="degrade" interval="1" />
               </components>
            </environment>
         </series>
```

**SUPPLEMENTARY TABLE 9. Difference metric for `tissue` simulations.** Simulations were averaged across replicates, ranked by difference metric in `tissue` simulations, but are shown with rank in `dish` simulations.

| CAR T-cell Dose | $CD4^+$:$CD8^+$ T-Cell Ratio | CAR Affinity (M) | Antigens Cancer | Antigens Healthy | Normalized Cancer Cell Count | Normalized Healthy Cell Count | Normalized T-cell Count | Difference Metric Value | Rank in `dish` | Rank in `tissue` |
|---|---|---|---|---|---|---|---|---|---|---|
| 1000 | 0.25 | 1E-06 | 10000 | 100 | 0.15 | 0.90 | 39.20 | 0.04 | 1 | 1 |
| 1000 | 0.5 | 1E-06 | 10000 | 100 | 0.17 | 0.91 | 39.27 | 0.02 | 3 | 2 |
| 1000 | 0.25 | 1E-06 | 5000 | 100 | 0.21 | 0.91 | 39.01 | -0.02 | 4 | 3 |
| 500 | 0.25 | 1E-06 | 10000 | 100 | 0.23 | 0.91 | 58.23 | -0.03 | 2 | 4 |
| 1000 | 0.75 | 1E-07 | 1000 | 100 | 0.22 | 0.84 | 39.65 | -0.04 | 11 | 5 |
| 1000 | 0.75 | 1E-06 | 10000 | 100 | 0.24 | 0.92 | 38.46 | -0.05 | 6 | 6 |
| 1000 | 0.5 | 1E-06 | 5000 | 100 | 0.27 | 0.92 | 38.50 | -0.07 | 8 | 7 |
| 1000 | 0.5 | 1E-07 | 500 | 100 | 0.25 | 0.82 | 40.36 | -0.08 | 12 | 8 |
| 500 | 0.5 | 1E-06 | 10000 | 100 | 0.28 | 0.92 | 56.74 | -0.08 | 5 | 9 |
| 500 | 0.5 | 1E-07 | 1000 | 100 | 0.26 | 0.82 | 56.78 | -0.09 | 10 | 10 |
| 500 | 0.25 | 1E-06 | 5000 | 100 | 0.33 | 0.93 | 56.29 | -0.13 | 9 | 11 |
| 500 | 0.25 | 1E-07 | 500 | 100 | 0.31 | 0.82 | 59.52 | -0.13 | 14 | 12 |
| 250 | 0.25 | 1E-06 | 10000 | 100 | 0.33 | 0.92 | 88.55 | -0.13 | 7 | 13 |
| 250 | 0.25 | 1E-07 | 1000 | 100 | 0.33 | 0.84 | 96.43 | -0.15 | 13 | 14 |

### 4 Supplementary Methods Details

#### 4.1 Model framework

The model integrates CAR T-cell agents into the agent-based modeling framework ARCADE (Yu and Bagheri, 2020; 2021). The ARCADE framework utilizes interfaces to enable modular model composition. CARCADE implements the Cell interface for CAR T-cells. This CAR T-cell class extends into two subclasses representing $CD4^+$ and $CD8^+$ CAR T-cells. Each subclass contains two modules controlling metabolism and inflammation. All existing parameters are kept at default values (Yu and Bagheri, 2020; 2021). All new parameters specific to CARCADE are listed in **Supplementary Table 1** and described below.

#### 4.2 Tissue cell agents

Tissue cell agents, which represent cancer and healthy tissue cells, can enter any one of seven cell states: quiescent, migratory, proliferative, apoptotic, necrotic, senescent, and uncommitted by following a specific set of rules (Yu and Bagheri, 2020). Tissue cells in this study use the default metabolism and signaling modules. Cancer cell agents are identical to healthy cells except that they can escape quiescence, are more amenable to cell crowding upon looking for new locations while migrating or proliferating, and they differ in parameter name for antigen expression level. To interact with CAR T-cell agents, two parameters were added to tissue cell agents: the number of antigens (`CAR_ANTIGENS_CANCER` for cancer cells and `CAR_ANTIGENS_HEALTHY` for healthy cells) and the number of PD-1 ligand "self targets"expressed on the cell surface (`SELF_TARGETS`).

#### 4.3 CAR T-cell agents

##### 4.3.1 Initialization

All CAR T-cells are initialized with an age pulled from a uniform distribution with a specified minimum (`T_CELL_AGE_MIN`) and maximum (`T_CELL_AGE_MAX`) age, a volume pulled from a normal distribution with a specified average (`T_CELL_VOL_AVG`) and range (`T_CELL_VOL_RANGE`), and an approximate age at which death is more likely to occur from a normal distribution with specified average (`DEATH_AGE_AVG_T`) and range (`DEATH_AGE_RANGE`, parameter as in ARCADE) (Yu and Bagheri, 2020). To account for the Hayflick limit, each CAR T-cell is also initiated with a maximum division potential (`DIVISION_POTENTIAL_T`). Additionally, CAR T-cells are initiated with a number of surface CAR receptors (CARS) and surface self (PD1) receptors (`SELF_RECEPTORS`).

##### 4.3.2 States and rules

Though it is difficult to directly observe transitions between states in individual cells, discrete CAR T-cells states are generally agreed to exist and mechanisms are hypothesized for state transitions (Kasakovski et al., 2018; Martinez and Moon, 2019; Xu et al., 2019). When needed, we rely on more general T-cell studies to define parameters and state transitions. CAR T-cells can enter one of eleven states: migratory, proliferative, cytotoxic, stimulatory, paused, senescent, apoptotic, exhausted, anergic, starved, and uncommitted.

CAR T-cell agents move through the state diagram shown in **Supplementary Figure 1** at each time point as a function of their state at the start of the time point.

###### 4.3.2.1 Migratory state

Migratory is the default state for a healthy, activated or yet-to-be activated T-cell agent as it travels around looking for potential threats. Upon entering the migratory state, the time it takes a cell to migrate is determined as a function of the distance the cell is moving and the speed at which the cell is moving. Cells assess their current and surrounding locations to determine valid locations to which they can move. To be a valid location, a location must meet the following checks: (i) adding the cell to the location must not increase the total volume of all cells in that location to be greater than the volume of that location, and (ii) adding the cell to the location cannot increase the number of agents in that location beyond the max number of allowed agents. CAR T-cells assign a score to each location meeting the above criteria, where the score is a function of the amount of free glucose and the number of cancer cells in the location. CAR T-cells move towards the location with the highest score, though there is a level of inaccuracy introduced in assessing the amount of glucose in each location. The score ($S_{loc}$) is given by:

$$S_{loc} = \left[\beta \frac{G_i}{G^{\circ}} + (1-\beta)u\right] + C_i$$

where

- $\beta$ is the accuracy (`ACCURACY`)
- $G^{\circ}$ is the source concentration of glucose (`CONC_GLUC`)
- $G_i$ is the amount of glucose in the location being assessed
- $u$ is a random number drawn from a uniform distribution $U([0,1])$
- $C_i$ is the number of cancer cells in the location

Accounting for the number of cancer cells in each location within the score serves as a proxy for the bias of T-cells to move towards cytokines and chemokines indicating necessary immune activity. If there are no locations available, the cell becomes paused.

###### 4.3.2.2 Proliferative state

Proliferative CAR T-cell agents, like typical T-cells, asymmetrically divide, splitting their volume unevenly, to produce daughter cell agents (Verbist et al., 2016). Upon entering the proliferative state, cells either divide or exit the proliferative state if the cell becomes no longer able to proliferate. Specifically, at each time step, the cell checks whether it has entered a state, such as apoptotic, such that it is no longer able to proliferate and checks if there are no locations into which the cell can divide. If there are no available locations for the daughter cell, the dividing cell becomes paused. To successfully proliferate, the cell must double in volume, where the rate at which this occurs is dictated by the metabolism module. Once this check has passed, the cell checks if the time since entering the proliferative state has exceeded the required DNA replication time, which is calculated as the average DNA synthesis time (`SYNTHESIS_TIME_T`) plus or minus a randomly drawn value within the DNA synthesis time range (`SYNTHESIS_TIME_T_RANGE`). If both checks are met, the cell divides, creating a daughter cell with 50% plus or minus up to 5% of the volume, bound IL-2, and granzyme (if $CD8^+$). The division count for both the parent and daughter cell is decreased. The daughter cell inherits the number of “self” (PD1) receptors, number of times the parent cell bound to antigen, the number of times the parent cell bound to “self” (PD1) ligands, and the activation condition. The duration of time spent in the proliferative state is defined as a cell cycle length, which is recorded each time a cell divides. The new daughter cell's location is determined in the same way as described for migratory cells.

###### 4.3.2.3 Paused state

Cell agents become paused when they are unable to migrate or proliferate. Paused agents have no active cell behavior, but they will remain paused until they enter a different state. Agents may accumulate in the paused state over time, as this behavior represents a biological phenomenon.

###### 4.3.2.4 Stimulatory and cytotoxic states

Stimulatory and cytotoxic states represent the effector functions of $CD4^+$ and $CD8^+$ CAR T-cells, respectively. Stimulatory CAR T-cell agents produce IL-2, while cytotoxic CAR T-cell agents produce granzyme and, once bound to a target, kill the target cell, which becomes apoptotic. Effector cells enter the uncommitted state after a time delay to represent how long a T-cell stays bound to a target. This time delay is calculated as the average time bound to a target (`BOUND_TIME`) plus or minus a randomly drawn value from within the bound time range (`BOUND_TIME_RANGE`). Once a cell is activated it remains activated until it (i) becomes deactivated over time by not interacting with antigen for 7 days (Pearce, 2010), (ii) enters the anergic or exhausted states, as these cause cells to lose effector function, or (iii) dies. Activation biases previously activated cells towards proliferation over migration when the agent has not bound to a surrounding target in subsequent time steps. Additionally, activation strongly influences cell effector function and metabolism.

###### 4.3.2.5 Anergic state

Anergy is a non-functional, undesired T-cell state induced from either (i) T-cell stimulation via antigen interaction in the absence of proper co-stimulation (Macian et al., 2004; Wherry, 2011; Crespo et al., 2013; Wherry and Kurachi, 2015; Kasakovski et al., 2018) or by (ii) simultaneous T-cell stimulation with antigen and co-inhibitory signals, such as self-identification signals (Crespo et al., 2013; Kasakovski et al., 2018). Proper co-stimulation is often conferred by co-stimulatory receptors such as CD28 (Schwartz, 1996; Crespo et al., 2013). Since second-generation CARs and beyond contain these co-receptors, anergy induction caused by T-cell stimulation via antigen interaction in the absence of proper co-stimulation is less likely. Only the latter possible mechanism is considered in the model, as we assume CARs in the model are at least second generation, as these are the only FDA approved and studied CARs in current research. CARCADE CAR T-cell agents enter the anergic state when they receive mixed signals, binding to both the antigen and the "self" receptors. Upon binding to both signals, cells have some probability (`ANERG_FRAC`) of undergoing apoptosis or otherwise become anergic. Though most anergy studies come from work on $CD4^+$ T-cells (Wherry, 2011), some studies show $CD8^+$ T-cells can enter this state (Schwartz, 2003). In the model, both $CD4^+$ and $CD8^+$ T-cell agents can become anergic.

Anergic cells characteristically exhibit little to no effector function (Wherry and Kurachi, 2015), proliferative potential (Macian et al., 2004; Wherry and Kurachi, 2015), no IL-2 production (Schwartz, 2003; Macian et al., 2004), and an inability to respond to subsequent proper stimulation (Macian et al., 2004; Wherry, 2011). Thus, agents in the anergic state de-activate if previously activated, turning off all effector function, and they can only escape from the anergic state through eventual cell death.

Anergic T-cells escape this state either by eventual induction of apoptosis or, if induced due to lack of proper co-stimulation, by sufficient uptake of IL-2 to recover proper cell function (Schwartz, 2003; Macian et al., 2004). Anergy occurs within the time frame of a few days (Wherry and Kurachi, 2015), making it particularly relevant for this model, which can simulate tumor growth for up to a few months, and for understanding CAR T-cell dynamics. Since IL-2 only recovers cells induced into anergy by lack of proper co-stimulation, and this is less likely to be an issue in CAR T-cells, IL-2 recovery from anergy is not included in CARCADE and anergy is an irreversible state in the model.

###### 4.3.2.6 Exhausted state

Similar to anergy, exhaustion is a distinct, non-functional T-cell state induced by repeated activation (Crespo et al., 2013; Kasakovski et al., 2018). Though exhaustion occurs on the time scale of weeks (Wherry and Kurachi, 2015), this state is highly prevalent in CAR T-cell work and is one attributed cause of low therapeutic efficacy (Long et al., 2015). Including exhaustion as a state in the model is therefore relevant for understanding and improving CAR T-cell dynamics.

Most research on exhaustion focuses on $CD8^+$ T-cells (Akbar and Henson, 2011), but exhaustion also occurs in $CD4^+$ T-cells (Akbar and Henson, 2011; Wherry and Kurachi, 2015). Exhausted $CD8^+$ T-cells lose cytotoxic activity (Akbar and Henson, 2011), while exhausted $CD4^+$ T-cells express significantly decreased levels of effector cytokines (Wherry and Kurachi, 2015). In the model, both subtypes of CAR T-cell agents can become exhausted.

To capture the dynamics that induce exhaustion over time, CAR T-cells count of the number of times they have bound to antigen. If a cell goes 24 h without binding antigen, the count decreases by one. If this count exceeds a set maximum (`MAX_ANTIGEN_BINDING`), the next binding event will cause them to become exhausted. Upon exceeding the maximum antigen binding count, cells have some probability (`EXHAU_FRAC`) of undergoing apoptosis but otherwise become exhausted. Exhausted T-cells characteristically exhibit little to no proliferative potential (Akbar and Henson, 2011; Wherry, 2011; Wherry and Kurachi, 2015), high expression of PD-1 (Akbar and Henson, 2011; Crespo et al., 2013; Wherry and Kurachi, 2015; Kasakovski et al., 2018), and higher rates of apoptosis (Long et al., 2015). Though not fully inert (Wherry and Kurachi, 2015), exhausted T-cells lose some effector functions before others (Akbar and Henson, 2011). For simplicity, CAR T-cell agents in the model lose all effector function simultaneously upon entering the exhausted state. Exhausted agents de-activate if previously activated, turning off all effector function, and can only escape from the exhausted state through eventual cell death.

Exhaustion was thought to be reversible with PD-1 blockades (Akbar and Henson, 2011; Wherry, 2011; Wherry and Kurachi, 2015), but new data suggest that PD-1 blockades promote expansion of T-cell populations outside of those exhausted within the tumor (Yost et al., 2019). Both hypotheses motivate combining CAR T-cell therapy with either internally-engineered or intravenously injected PD-1 blockades (Cherkassky et al., 2016; Liu et al., 2016; Maus and June, 2016; Rafiq et al., 2018). However, PD-1 blockades are not included in the model at present, and exhaustion is an irreversible state in the model.

###### 4.3.2.7 Senescent state

Due to the natural ageing process, all cells can become senescent, which is a non-reversible state causing cells to undergo cell cycle arrest and stop proliferating (Akbar and Henson, 2011; Crespo et al., 2013; Kasakovski et al., 2018). Upon hitting their division limit (`DIVISION_POTENTIAL_T`), cells have some probability (`SENES_FRAC`) of becoming senescent or apoptotic. Senescent cells remain in this state until they are eventually removed from the simulation due to age-induced apoptotic cell death.

###### 4.3.2.8 Starved state

Cells require sufficient nutrients to sustain normal cellular function. Cells that do not meet their energy needs, as dictated by the metabolism module, become starved. Cells can escape the starved state upon recovering from the energy deficient by accumulation of energy. Recovered cells are set to the uncommitted state to then continue in the decision sequence.

###### 4.3.2.9 Apoptotic state

Cells can enter the apoptotic state due to age or sustained lack of nutrients or energy. Cells have an increased probability of entering the apoptotic state once they exceed their average life span (`DEATH_AGE_AVG_T`) as defined by a cumulative normal distribution where the mean is set to the `DEATH_AGE_AVG_T` and the standard deviation is set to the `DEATH_AGE_RANGE`. Cells can also become apoptotic under conditions of sustained energy deficiency, meaning their energy goes below a set threshold (`ENERGY_THRESHOLD`). Upon entering the apoptotic state, the cell is removed from the simulation after a time delay (`DEATH_TIME`) representing the time it takes a cell to die by apoptosis.

##### 4.3.3 Antigen-induced activation process

Upon antigen-induced activation, CAR T-cells enter an effector state based on subtype, entering either the cytotoxic state to cytotoxically kill target agents or the stimulatory state to stimulate other T-cells by releasing cytokines (Pearce, 2010). While there is evidence that both T-cell subtypes can become cytotoxic and stimulatory, $CD8^+$ T-cells primarily provide cytotoxic functions, while $CD4^+$ T-cells primarily provide stimulatory functions (Liadi et al., 2015; Golubovskaya and Wu, 2016; Sommermeyer et al., 2016). For simplicity, the model assumes only $CD8^+$ CAR T-cell agents can enter the cytotoxic state and only $CD4^+$ CAR T-cell agents can enter the stimulatory state to perform associated functions. As the time required for CAR T-cells to form stable and functional immune synapses is shorter than two min (Watanabe et al., 2018), we assume signal binding and activation occur within a single time step within the model, which equates to one min.

The probability of an antigen binding or "self" (PD1) binding events occurring are a function of affinity of a receptor for its ligand, the number of ligands on the target cell surface, the number of receptors on the CAR T-cell surface, distance from and contact with a target cell, and probability of receptors making contact. We developed a sequence of events and a binding probability heuristic that captures these general trends. After stepping their metabolism module, CAR T-cell agents in the paused or uncommitted states assess their surroundings and randomly select one neighboring agent. If that agent is a CAR T-cell agent, nothing happens and the searching CAR T-cell agent goes on to assess another target as until it hits the max number of neighbors assessable in a given time point (`SEARCH_ABILITY`). If the found agent is a tissue cell, the probability of binding and killing $P(binding\ and\ killing)$ is calculated according to a heuristic equation:

$$P(binding\ and\ killing) = 2\left(\frac{1}{1+e^{-x}}\right) - 1$$

where

$$x = \left(\frac{\gamma L_{target}}{\beta K_D V_{loc} N_A + \gamma L_{target}}\right)\left(\frac{R}{R_{avg}}\right)\alpha$$

where:

- $L_{target}$ is the number of ligands on the target cell (CAR_CANCER_ANTIGENS or CAR_HEALTHY_ANTIGENS for cancer and healthy cells, respectively, for CAR-antigen binding events and SELF_LIGANDS for self-receptor binding events)
- $R$ is the number of CARs for CAR-antigen binding events or number of self-receptors for self-receptor binding events on the CAR T-cell
- $R_{avg}$ is the average number of receptors on the CAR T-cell (CARS) for CAR-antigen binding events and is the number of self-receptors (SELF_RECEPTORS) a cell starts with for self-receptor binding events
- $K_D$ is the affinity of the receptor for the antigen in M ( CAR_AFFINITY for CAR-antigen binding events and SELF_AFFINITY for self-receptor binding events)
- $V_{loc}$ is the volume of the location in L
- $N_A$ is Avogadro's Number
- $\gamma$ is the contact fraction (CONTACT_FRAC)
- $\alpha$ (CAR_ALPHA for CAR-antigen binding events and SELF_ALPHA for self-receptor binding events) and $\beta$ (CAR_BETA for CAR-antigen binding events and SELF_BETA for self-receptor binding events) are fitting factors

Overall, this function produces trends fitting with the expected outcomes where increasing antigen/ligand number, receptor number, and $K_D$ will result in a higher probability if binding (for self-receptor binding events) and/or killing (for CAR binding events), as shown in **Supplementary Figure 2**, which matches previously determined T-cell activation curves (Hernandez-Lopez et al., 2021). This calculation is done for both the CAR-antigen binding event and the PD1-PDL1 binding event. If the CAR T-cell binds to antigen and not to "self", the agent will become activated, enter its effector state, and increase the antigen-binding counter. If the CAR T-cell binds to both antigen and "self", the agent will become anergic (more detail described below) and increase the antigen-binding counter. If the CAR T-cell binds to neither antigen nor "self" or only to "self", the agent becomes either migratory or proliferative, biasing towards proliferative if the CAR T-cell agent is activated. If the cell is not activated, cells become proliferative with a given probability (PROLI_FRAC); otherwise, it becomes migratory.

Additionally, upon each binding event in which cells bind to antigen, independent of binding to "self" receptor, CAR T-cells increase the amount of "self" receptors on their surface according to the following equation:

$$R_{i+1} = R_i + uR_0$$

where:

- $R_0$ is the initial number of "self" receptors on the cell
- $u$ is a random number drawn from a uniform distribution $U([0.95,1.05])$
- $R_i$ is the number of "self" receptors on the cell after $i$ binding events
- $R_{i+1}$ is the number of "self" receptors on the cell after the new binding event

Thus, after each binding event, the number of "self" receptors on the cell surface will increase by 95-105% of the original number of receptors (`SELF_RECEPTORS` for initialized cells) after each binding event. This increase serves as both a marker for exhaustion (described in more detail below) and can increase the probability of a cell becoming anergic (described in more detail below) over time.

##### 4.3.4 Subcellular modules

###### 4.3.4.1 Inflammation module

T-cell cytokine signaling is dynamic and provides self-feedback. Additionally, cytokines influence effector function, metabolism, growth, and proliferation (Huang et al., 2018). Though many important cytokines exist, the model only utilizes IL-2, which is a well-studied driver of immune response and is FDA approved as an intravenously administered immunotherapy treatment (Rosenberg, 2014; Ross and Cantrell, 2018).

Unstimulated T-cells do not express IL-2Rα until after antigen-induced activation or stimulation with IL-2, making cells more sensitive to IL-2 after activation to amplify the immune response (Malek and Castro, 2010; Liao et al., 2013; Hernandez-Lopez et al., 2021). Upon stimulation, IL-2 (i) promotes T-cell growth and proliferation by upregulating glycolysis and glucose uptake and (ii) induces effector function by activating relevant genes (Malek and Castro, 2010; Liao et al., 2013). Effector function varies for T-cell subtypes; $CD4^+$ T-cells primarily secrete cytokines, such as IL-2, while $CD8^+$ T-cells produce cytolytic material, such as granzymes, that kills bound target cells when secreted (Liao et al., 2013).

All CAR T-cells are equipped with inflammation modules specific to their cell type. Broadly, the $CD4^+$ Inflammation module produces IL-2, while the $CD8^+$ Inflammation module produces granzyme that is used to kill target cells. Both T-cell subtypes bind IL-2 using a receptor complex composed of three receptor chains: IL-2Rα, IL-2Rβ, and IL-2R$\gamma_c$. IL-2 binds to IL-2Rα weakly, the two-chain receptor complex IL-2R$\beta\gamma_c$ with intermediate affinity, and the full three-part complex with high affinity (Malek and Castro, 2010; Liao et al., 2013; Ross and Cantrell, 2018). Both cell agents use the same set of ordinary differential equations (ODEs) to determine the amount of IL-2 bound to their surface. All species and parameters within the ODEs are detailed in the table below.

| **Species** | | |
|---|---|---|
| **Name** | **Description** | **Symbol** |
| External IL-2 | IL-2 in the environment accessible to the cell | $X_1$ |
| IL-2R$\beta\gamma_c$ | lower affinity two-chain IL-2 receptor complex | $X_2$ |
| IL-2R$\beta\gamma_c\alpha$ | higher affinity three-chain receptor complex | $X_3$ |
| IL-2R$\beta\gamma_c$:IL-2 | IL-2 bound two-chain IL-2 receptor complex | $X_4$ |
| IL-2R$\beta\gamma_c\alpha$:IL-2 | IL-2 bound three-chain IL-2 receptor complex | $X_5$ |
| Total unbound IL-2 receptors | Total receptors (two- and three-chain) on cell surface not bound to IL-2 | $X_6$ |
| Total bound IL-2 | Total receptors (two- and three-chain) on cell surface bound to IL-2 | $X_7$ |

| **Parameters** | | | |
|---|---|---|---|
| **Name** | **Description** | **Parameter Name** | **Symbol** |
| Two-chain complex IL-2 binding on rate | Rate of IL-2 binding to IL-2R$\beta\gamma_c$ | `IL2_BINDING_ON_RATE_MIN` | $k_{on,2}$ |
| Three-chain complex IL-2 binding on rate | Rate of IL-2 binding to IL-2R$\beta\gamma_c\alpha$ | `IL2_BINDING_ON_RATE_MAX` | $k_{on,3}$ |
| IL-2 binding off rate | Rate of IL-2 unbinding from IL-2R$\beta\gamma_c$ or IL-2R$\beta\gamma_c\alpha$ | `IL2_BINDING_OFF_RATE` | $k_{off}$ |
| Three-chain receptor conversion rate | Captures rate of conversion of the two-chain complex IL-2R$\beta\gamma_c$, whether | `K_CONVERT` | $k_{convert}$ |

| | | | |
|---|---|---|---|
| | bound or unbound, into the three-chain complex IL-2Rβγα though positive feedback | | |
| IL-2 receptor recycle rate | Rate at which IL-2Rβγc:IL-2, IL-2Rβγcα:IL-2, or IL-2Rβγcα are internalized be converted via recycle back into unbound IL-2Rβγc chains. | K_REC | $k_{rec}$ |

External IL-2 ($X_1$) binds reversibly to unbound receptor complexes $X_2$ and $X_3$ with on rate $k_{on,2}$ and $k_{on,3}$, respectively, to from the bound receptor complexes $X_4$ and $X_5$, respectively. The high affinity and lower affinity receptors bind with the same off rate $k_{off}$ but different on rates (Lauffenburger and Linderman, 1993). The equation describing $X_1$ kinetics are as follows:

$$\frac{dX_1}{dt} = k_{off}X_4 + k_{off}X_5 - k_{on,2}X_1X_2 - k_{on,3}X_1X_3$$

The IL-2Rα subunit, which converts two-chain complexes into three-chain complexes with higher IL-2 affinity, is only produced after initial binding of IL-2 to the two-chain complex. To reduce the model complexity and the number of species tracked, the production of the IL-2Rα chain alone is not explicitly modeled. However, the impact of IL-2 binding on conversion of the two-chain complex IL-2Rβγc, whether bound or unbound, into the three-chain complex IL-2Rβγα though positive feedback is captured through the parameter $k_{convert}$, where the summed number of bound IL-2 complexes is meant to represent the magnitude of signal that produces the IL-2Rα chain (Busse et al., 2010). Unbound and bound receptors without the IL-2Rα chain, $X_2$ and $X_4$, can be converted to three-chain receptor complexes $X_3$ and $X_5$, respectively, through the convert mechanism. Additionally, since the $X_2$ is constitutively expressed (Malek and Castro, 2010), we capture this process by having any IL-2 bound chains $X_4$ and $X_5$ or unbound chain $X_3$ that are internalized be converted via recycle, represented by $k_{rec}$, back into unbound $X_2$. This recycling process enables the response of the production of the IL-2Rα chain to be pulsatory and will eventually stop in the prolonged absence of IL-2. The equation describing $X_2$, $X_3$, $X_4$, and $X_5$ kinetics are as follows:

$$\frac{dX_2}{dt} = k_{off}X_4 - k_{on,2}X_1X_2 - k_{convert}(X_4 + X_5)X_2 + k_{rec}(X_3 + X_4 + X_5)$$

$$\frac{dX_3}{dt} = k_{off}X_5 - k_{on,3}X_1X_3 + k_{convert}(X_4 + X_5)X_2 - k_{rec}X_3$$

$$\frac{dX_4}{dt} = k_{on,2}X_1X_2 - k_{off}X_4 - k_{convert}(X_4 + X_5)X_4 - k_{rec}X_4$$

$$\frac{dX_5}{dt} = k_{on,3}X_1X_3 - k_{off}X_5 + k_{convert}(X_4 + X_5)X_4 - k_{rec}X_5$$

For convenience, we also track $X_6$ and $X_7$.

$$\frac{dX_6}{dt} = \frac{dX_2}{dt} + \frac{X_3}{dt}$$

$$\frac{dX_7}{dt} = \frac{dX_4}{dt} + \frac{dX_5}{dt}$$

Each species is in units of molecules of IL-2 in ODEs, but the environment keeps track of the concentration of IL-2 in units of molecules/cm$^3$. The ODEs are run within each individual cell at each model step (one minute), using a Runge-Kutta solver, with a time step of 1/3$^{rd}$ of a second. This set of reduced equations captures a few key aspects of IL-2 signaling with the brevity necessary to run the model without excess delay, as these ODEs are run in all CAR T-cells and on a large scale are very computationally expensive.

The amount of external IL-2 is determined based on both the cell's location within the environment and the distance from the cell surface a cell can sense ($d_{shell}$, `SHELL_THICKNESS`). T-cells are relatively small compared to the volume of a location in the model, and they have access to all the IL-2 present in the environment. A shell thickness value was set such that cells can sense a few microns out from their external surfaces (Gherbi et al., 2018). The cell's volume ($V_{cell}$) is known for each agent, and thus, assuming the cell to be a sphere, the cell radius ($r_{cell}$) can be calculated as follows:

$$r_{cell} = \sqrt[3]{\frac{3}{4\pi} V_{cell}}$$

Subsequently, the shell radius ($r_{shell}$), which is the distance of the cell radius and the distance from the cell surface that a cell can sense ($d_{shell}$, `SHELL_THICKNESS`), can be calculated as follows:

$$r_{shell} = r_{cell} + d_{shell}$$

Thus, the volume within the shell ($V_{shell}$) that exists between the external surface of the cell, assuming the cell to be a sphere, and the distance from the cell defined by this shell's thickness is calculated as follows:

$$V_{shell} = V_{r_{shell}} - V_{cell} = V_{cell}\left(\frac{r_{shell}^3}{r_{cell}^3} - 1\right)$$

The fraction of total volume in the environment that makes up this shell volume ($f_{shell}$) is the calculated as follows:

$$f_{shell} = \frac{V_{shell}}{V_{loc}}$$

The amount of external IL-2 ($X_1$) a cell as access to at each time point is calculated as follows:

$$X_1 = f_{shell} n_{IL2}$$

where $n_{IL2}$ is the total number of molecules of IL-2 in the environment at the cell's location.

The inflammation module includes a memory of the total amount of IL-2 bound on the surface to capture a time delay in various cellular processes that are a function of IL-2 binding, as no process is instantaneous and must first undergo internal signaling networks to initiate cellular responses.

When a CAR T-cell divides, the amount of each species, with the exception of IL-2Rβγc but including granzyme in the CD8+ CAR T-cells, is divided between the two daughter cells, splitting according to the same fraction described in the proliferative state section. Since IL-2Rβγc is assumed to be constitutively expressed, the amount in the daughter cell is the steady state value of IL-2 receptors total (`IL2_RECEPTORS`) minus the amount of receptors that are already bound to IL-2.

Each CAR T-cell subtype has specific functions corresponding to the typical functions of T-cell subtypes. While IL-2 is secreted primarily by CD4+ T-cells after antigen-induced activation, the model assumes IL-2 is exclusively secreted by this cell subtype (Malek and Castro, 2010; Liao et al., 2013; Rosenberg, 2014). Upon antigen-induced activation, IL-2 drives the production of granzyme and other cytotoxins in CD8+ T-cells (Liao et al., 2013). Though CD4+ CAR T-cells have been found to be capable of killing, it is at a much slower rate and most killing is done by CD8+ CAR T-cells (Liadi et al., 2015). For simplicity, the model assumes all granzyme production and cytotoxic killing is done exclusively by CD8+ CAR T-cells.

###### 4.3.4.1.1 CD4+ CAR T-cell agent IL-2 production

CD4+ CAR T-cell agents produce IL-2 in both an antigen-induced independent and dependent manner (Busse et al., 2010). Independent of antigen-induced activation, CD4+ T-cells produce IL-2 as a function of the amount of IL-2 bound on their surface due to positive feedback (Busse et al., 2010). The maximum amount of IL-2 produced due to IL-2 feedback (`IL2_PROD_RATE_IL2`) is scaled by the amount of IL-2 bound at a previous time point corresponding to the delay necessary to turn on IL-2 production (`IL2_SYNTHESIS_DELAY`) (Iwashima, 2003). The rate of IL-2 production as a function of IL-2 feedback at time step $t$ ($r_{IL2}^{t}$) in units of molecules/cell/min is calculated as follows:

$$r_{IL2}^{t} = R_{IL2,}\left(\frac{n_{IL2}^{t-\tau_{IL2}}}{N_{IL2}}\right)$$

where:

- $R_{IL2}$ is the maximum rate of production of IL-2 per time step due to IL-2 feedback in units of molecules/cell/min
- $n_{IL2}^{t-\tau_{IL2}}$ is the total amount of IL-2 bound to the cell surface at the previous time point corresponding to delay in IL-2 synthesis (`IL2_SYNTHESIS_DELAY`)
- $N_{IL2}$ is the maximum amount of IL-2 that can be bound to a cell, which corresponds to the total number of IL-2 receptors (`IL2_RECEPTORS`)

Upon antigen-induced activation and after a time delay (`IL2_SYNTHESIS_DELAY`), CD4+ CAR T-cells begin to produce additional IL-2 at a constant rate (`IL2_PROD_RATE_ACTIVE`) that is added to the rate of production of IL-2 per time step due to IL-2 feedback. The equation for the total amount of IL-2 produced by the cell in a given ($r$) in units of molecules/cell/min is as follows:

$$r = \begin{cases} r_{IL2} + r_{active}, & if\ active\ and\ t_{active} > \tau_{IL2} \\ r_{IL2}, & else \end{cases}$$

where:

- $r_{active}$ is the rate of IL-2 production due to antigen-induced activation (IL2_PROD_RATE_ACTIVE) in units of molecules/cell/min
- $t_{active}$ is the length of time since the T-cell was activated
- $\tau_{IL2}$ is the time delay required to synthesized IL-2 (IL2_SYNTHESIS_DELAY)

The IL-2 produced during this time step is then added to the IL-2 in the local environment.

###### 4.3.4.1.2 CD8$^+$ CAR T-cell agent granzyme production

CD8$^+$ CAR T-cell agents produce granzyme upon antigen-induced activation as a function of IL-2 (Janas et al., 2005). Granzyme increases linearly as a function of IL-2 until it eventually plateaus (Janas et al., 2005). Additionally, granzyme builds up in a cell over time. In the model, to account for delay in granzyme production after antigen-induced activation due to internal signal transduction, granzyme production begins after a time delay (GRANZ_SYNTHESIS_DELAY). The amount of IL-2 sensed by the cell is used to scale the maximum rate of granzyme production and is calculated using the same time delay. Overall, the amount of granzyme in a CD8$^+$ CAR T-cell each time step ($n_g^t$) in arbitrary units is calculated as follows:

$$n_g^t = n_g^{t-1} + G\left(\frac{n_{IL2}^{t-\tau_g}}{N_{IL2}}\right)$$

where:

- $n_g^{t-1}$ is the amount of granzyme in the cell in the previous time step in arbitrary units
- $G$ is the moles of granzyme produced per moles of IL-2 (GRANZ_PER_IL2)
- $n_{IL2}^{t-\tau_g}$ is the total amount of IL-2 bound to cell surface at the previous time point corresponding to the delay in granzyme synthesis ($\tau_g$, GRANZ_SYNTHESIS_DELAY)
- $N_{IL2}$ is the maximum amount of IL-2 that can be bound to a cell, which corresponds to the total number of IL-2 receptors (IL2_RECEPTORS)

As described in the cytotoxic state section, one unit of granzyme in arbitrary units is lost when a target cell is killed.

##### 4.3.4.2 Metabolism module

T-cell metabolism is complex, as it is a function of both antigen-induced activation and IL-2. Naïve, inactivated T-cells are metabolically quiescent, require less oxygen and glucose consumption, and primarily utilize oxidative phosphorylation (OXPHOS) and fatty acid oxidation (FAO) for energy (Pearce, 2010; Buck et al., 2015; Golubovskaya and Wu, 2016; Mehta et al., 2017). Antigen-induced activation causes T-cells to shift their metabolism by upregulating glycolysis (Frauwirth et al., 2002; Jones and Thompson, 2007; Pearce, 2010; Altman and Dang, 2012; Gerriets and Rathmell, 2012; MacIver et al., 2013; Chang and Pearce, 2016; Mehta et al., 2017), increasing glucose uptake (Frauwirth et al., 2002; Jones and Thompson, 2007; Jacobs et al., 2008; Pearce, 2010; Altman and Dang, 2012; Gerriets and Rathmell, 2012; Chang and Pearce, 2016; Golubovskaya and Wu, 2016), and downregulating mitochondrial metabolism (Gerriets and Rathmell, 2012; Mehta et al., 2017). This process is co-stimulation dependent, requiring signals like CD28, which is part of the CAR construct, to further activate the PI3K/Akt/mTOR pathway to increase Glut1 expression and thus increase glucose uptake and glycolysis (Frauwirth et al., 2002; Pearce, 2010; Gerriets and Rathmell, 2012; Chang and Pearce, 2016; Golubovskaya and Wu, 2016). IL-2 enhances this process, promoting glycolysis by further activating mTOR (Liao et al., 2013; Buck et al., 2015; Golubovskaya and Wu, 2016). Thus, both activation and IL-2 influence CAR T-cell metabolism in the model.

The metabolism module used for CAR T-cell agents uses IL-2 and antigen-induced activation to regulate T-cell energy requirements, nutrient uptake, metabolic preference for glycolysis, and cell mass production by building the default metabolism module used for tissue cells. The metabolism module calculates energy required to maintain antigen-induced activation (`ACTIVE_ENERGY`) in addition to basal, proliferative, and migratory energy requirements. Three parameters are altered as a function of antigen-induced activation and/or IL-2: (i) the metabolic preference for glycolysis over OXPHOS, (ii) the glucose uptake rate, and (iii) the fraction of internal nutrients converted to mass.

The metabolic preference for glycolysis over OXPHOS (`META_PREF`) is such that higher value dictates that a cell is getting more of its energy from glycolysis. This parameter changes as a function of both IL-2 bound to the cell agent as well as antigen induced activation independently. To account for delays in metabolic shifts due to intracellular signaling, a time delay is implemented much in the same way as for IL-2 and granzyme synthesis in the inflammation modules. The maximum possible influence of IL-2 on the metabolic preference (`META_PREF_IL2`) is scaled by the amount of IL-2 bound to the cell at the previous time point dictated by the time delay (`META_SWITCH_DELAY`) to calculate influence of IL-2 on the metabolic preference during time step $t$ ($m_{IL2}^{t}$) as follows:

$$m_{IL2}^{t} = M_{IL2}\left(\frac{n_{IL2}^{t-\tau_m}}{N_{IL2}}\right)$$

where:

- $M_{IL2}$ is the maximum possible influence of IL-2 on the metabolic preference (`META_PREF_IL2`)
- $N_{IL2}^{t-\tau_m}$ is the total amount of IL-2 bound to the cell surface at the previous time point corresponding to delay metabolic switching ($\tau_m$, `META_SWITCH_DELAY`)
- $N_{IL2}$ is the maximum amount of IL-2 that can be bound to a cell, which corresponds to the total number of IL-2 receptors (`IL2_RECEPTORS`).

This influence of IL-2 is added to the base metabolic preference value (`META_PREF`). Upon antigen-induced activation and after a time delay (`META_SWITCH_DELAY`), the influence of antigen-induced activation on the metabolic preference (`META_PREF_ACTIVE`) is also added to the base value. The total metabolic preference ($m$) during any given time step is calculated as:

$$m = \begin{cases} m_{base} + m_{IL2} + m_{active}, & if\ active\ and\ t_{active} > \tau_m \\ m_{base} + m_{IL2}, & else \end{cases}$$

where

- $m_{base}$ is the base value of the metabolic preference (`META_PREF`)

- $m_{IL2}$ is the calculated influence of IL-2 on metabolic preference
- $m_{active}$ is the influence of antigen-induced activation on metabolic preference (META_PREF_ACTIVE)
- $t_{active}$ is the length of time since the T-cell became activated
- $\tau_m$ is the time delay required to synthesize IL-2 (META_SWITCH_DELAY).

The glucose uptake rate parameter, like the metabolic preference, is a function of both IL-2 and antigen-induced activation, and follows the same formulation as the above parameter where the influence of IL-2 on glucose uptake rate during time step $t$ ($u_{IL2}^t$) is calculated as follows:

$$u_{IL2}^t = U_{IL2}\left(\frac{n_{IL2}^{t-\tau_m}}{N_{IL2}}\right)$$

where

- $U_{IL2}$ is the maximum possible influence of IL-2 on the glucose uptake rate (GLUC_UPTAKE_RATE_IL2)
- $n_{IL2}^{t-\tau_m}$ is the total amount of IL-2 bound to the cell surface at the previous time point corresponding to delay metabolic switching ($\tau_m$, META_SWITCH_DELAY)
- $N_{IL2}$ is the maximum amount of IL-2 that can be bound to a cell, which corresponds to the total number of IL-2 receptors (IL2_RECEPTORS).

This influence of IL-2 is added to the base glucose uptake rate (GLUC_UPTAKE_RATE). Upon antigen-induced activation and after a time delay ($\tau_m$, META_SWITCH_DELAY), the influence of antigen-induced activation on the metabolic preference (GLUC_UPTAKE_RATE_ACTIVE) is also added to the base value. The total glucose uptake rate ($u$) during any given time step is calculated as:

$$u = \begin{cases} u_{base} + u_{IL2} + u_{active}, & if\ active\ and\ t_{active} > \tau_m \\ u_{base} + u_{IL2}, & else \end{cases}$$

where:

- $u_{base}$ is the base value of the glucose uptake rate (GLUC_UPTAKE_RATE)
- $u_{IL2}$ is the calculated influence of IL-2 on glucose uptake rate
- $u_{active}$ is the influence of antigen-induced activation on glucose uptake rate (GLUC_UPTAKE_RATE_ACTIVE)
- $t_{active}$ is the length of time since the T-cell became activated
- $\tau_m$ is the time delay required to synthesized IL-2 (META_SWITCH_DELAY)

Using anabolic metabolism, which produces more growth-related intermediates than energy, enables effector T-cells to undergo rapid growth and proliferation, a necessary component of immune response (Pearce, 2010; MacIver et al., 2013). Additionally, through the support of upregulated glycolysis, T-cells upregulate biosynthesis pathways such as lipid, protein, and nucleic acid production (Jones and Thompson, 2007; MacIver et al., 2013). Thus, the fraction of internal nutrients converted to mass is also increased as function of antigen-induced activation. The total fraction of internal nutrients converted to mass ($f$) is calculated as follows:

$$f = \begin{cases} f_{base} + f_{active}, & if\ active\ and\ t_{active} > \tau_m \\ f_{base}, & else \end{cases}$$

where:

- $f_{base}$ is the base value of the fraction of nutrients converted to mass (`FRAC_MASS`)
- $f_{active}$ is the increase in the fraction of nutrients converted to mass as a function due to antigen-induced activation (`FRAC_MASS_ACTIVE`)
- $t_{active}$ is the length of time since the T-cell became activated
- $\tau_m$ is the time delay required to synthesized IL-2 (`META_SWITCH_DELAY`)

#### 4.4 Model environment

##### 4.4.1 Molecule diffusion

The model includes diffusion of IL-2 in addition to the default species (oxygen, glucose, and TGFα). The environment is initiated with a specified concentration of IL-2 (`CONCENTRATION_IL2`), which for this study was always zero. The diffusion of IL-2 is handled the same as the other species using a reaction-diffusion equation, where the diffusion rate of IL-2 is specified (`DIFFUSIVITY_IL2`). Parameters for oxygen, glucose, and TGFα are left at default values.

##### 4.4.2 Nutrient sources

The design of the nutrient (glucose and oxygen) sources in the simulation can be used to emulate specific contexts. `Dish` simulations utilize a constant source environment to replicate the evenly mixed nature of a well-mixed *in vitro* experiment (Yu and Bagheri, 2020). `Tissue` simulations utilize vasculature comprising two arteries and two veins that simulate realistic hemodynamics where vasculature can become increasingly damaged over time by cell crowding and movement (Yu and Bagheri, 2021).

#### 4.5 Cell placement and treatment

##### 4.5.1 Tissue cell placement

###### 4.5.1.1 Simulations in `dish`

For `dish` simulations, a specified total number of cells, with defined fractions of each population present, are plated randomly across the entire simulation environment. This plating aims to replicate an *in vitro* experiment where cells are plated in a monolayer in a cell culture dish with media.

###### 4.5.1.2 Simulations in `tissue`

For `tissue` simulations, a bed of healthy cells is placed throughout the simulation environment and a population of cancer cells is introduced to the center of the environment after a specified time delay. This tumor is then allowed to grow for 21 days before treatment with CAR T-cells. This setup aims to replicate a tumor growing within a bed of healthy, vascularized tissue that may present resource limitations in high cell density areas.

##### 4.5.2 CAR T-cell treatment

CAR T-cells are added at a specified time delay, with a specific CAR T-cell dose and $CD4^{+}$:$CD8^{+}$ ratio. Parameters for CAR T-cell populations, such as the affinity of the CAR for its antigen (`CAR_AFFINITY`), are specified in the population tags. CAR T-cell agents are biased toward spawning in locations with higher numbers of cancer cells to serve as a proxy for trafficking to tumors, which is not explicitly captured by the model. While placing CAR T-cells, each location is checked to ensure adding the agent will not (i) make the total cell volume exceed the volume of the location and (ii) cause tissue cells to exist beyond their tolerable heights (`MAX_HEIGHT`), which is described in ARCADE (Yu and Bagheri, 2020).

###### 4.5.2.1 Simulations in `dish`

CAR T-cells plated when sources or patterns are used for the nutrient environment, such as in `dish` simulations, can be placed in any location that does not exceed a maximum level of damage (`MAX_DAMAGE_SEED`). For simulations in this paper, source locations could not take damage.

###### 4.5.2.2 Simulations in `tissue`

When a dynamic graph vasculature is used for the nutrient environment, such as in `tissue` simulations, CAR T-cells can spawn in any location next to a vasculature graph edge where the radius of the vein is greater than or equal to the specified minimum value (`MIN_RADIUS_SEED`), representing CAR T-cell trafficking through vasculature and preventing cells from spawning at locations with excessive damage. Cells cannot spawn where there is no vasculature edge or there the radius is too small. This spawning setup aims to replicate CAR T-cell trafficking to tumors through vasculature without explicitly modeling transfer of the CAR T-cell from vasculature into the tissue.

#### 4.6 Simulated experiments

##### 4.6.1 Simulations of monoculture `dish`

We simulate *in vitro* monoculture experiments of cancer cells. Four different features (CAR T-cell dose, $CD4^{+}$:$CD8^{+}$ ratio, CAR affinity, and cancer antigens) were changed in this dataset, with 10 replicates for each possible combination of parameter choices, one in each category. **Supplementary Table 2** shows the set of simulated values per modified parameter. Simulated untreated cancer cells in `dish` (10 replicates) served as a negative control.

At the start of each simulation, 2000 cancer cells are plated randomly across a simulation environment with radius 34 and margin 6. Each cancer cell has the specified level of antigens (CANCER ANTIGENS, `CAR_ANTIGENS_CANCER`) for that combination of parameters. If treated, after a time delay of t = 10 min, a specified CAR T-cell dose (CAR T-CELL DOSE) with a specified $CD4^{+}$:$CD8^{+}$ ratio ($CD4^{+}$:$CD8^{+}$ RATIO) and CAR-antigen affinity (CAR AFFINITY) is added into the simulation. Data were collected every half day (720 time steps). In both treated and untreated simulations, the simulations lasted 7 days (10,080 time steps). Input files to create the simulations are shown in **Supplementary Table 7**.

##### 4.6.2 Simulations of ideal and realistic co-culture `dish`

This simulation setup aims to most closely replicate *in vitro* co-culture experiments with a mix of cancer and healthy cells. Four different axes (CAR T-cell dose, $CD4^{+}$:$CD8^{+}$ ratio, CAR affinity, cancer antigens) were changed to create both the ideal (antigen-negative healthy cells) and realistic (antigen-expressing healthy cells), with 10 replicates for each possible combination of parameter choices, one in each category. **Supplementary Table 3** shows the set of simulated values per modified parameter. Simulated untreated cancer cells in `dish` (10 replicates) served as a negative control.

At the start of each simulation, 2000 total cells (1000 cancer cells and 1000 healthy cells) are plated randomly across a 2D simulation with radius 34 and margin 6. Each cancer and healthy cell have the specified level of antigens (CANCER ANTIGENS/`CAR_ANTIGENS_CANCER` and HEALTHY ANTIGENS/`CAR_ANTIGENS_HEALTHY`, respectively) for that combination of parameters. If treated, after a time delay of t = 10 min, a specified CAR T-cell dose (CAR T-CELL DOSE) with a specified $CD4^+$:$CD8^+$ ratio ($CD4^+$:$CD8^+$ RATIO) and CAR-antigen affinity (CAR AFFINITY) is added into the simulation. Data were collected every half day (720 time steps). In both treated and untreated simulations, the simulations lasted 7 days (10,080 time steps). Input files to create the simulations are shown in **Supplementary Table 8**.

##### 4.6.3 Simulations of `tissue` with cancer and healthy cells

This simulation setup aims to most closely replicate *in vivo* experiments in which a tumor exists in a bed of healthy cells amongst a dynamic vasculature. We simulated the set of conditions described in **Supplementary Table 5**, co-culture `dish` conditions deemed as effective treatment conditions, in `tissue`. A set of 10 replicates of untreated cancer cells within a bed of healthy cells served as the control experiment.

At the start of each simulation, healthy cells are plated at one cell per location across the entire simulation with radius 34 and margin 6. The simulation environment uses the $S_{22}$ vasculature setup, where there are two arteries and two veins total, each one starting from a different side of the simulation and alternating between veins and arteries, described in the ARCADE vasculature study (Yu and Bagheri, 2021). A population of cancer cells is then inoculated into the model t = 1 d (1,440 time steps) into the simulation at the center out to a radius bounds of 0.05 (radii 1 and 2). Each cancer and healthy cell have the specified level of antigens (`CAR_ANTIGENS_CANCER` and `CAR_ANTIGENS_HEALTHY`, respectively) for that combination of parameters. The tumor is then allowed to grow until t = 31 d (44,640 time steps). If treated, the treatment begins at t = 22 d (31,680 time steps). A specified CAR T-cell dose (CAR T-CELL DOSE) with a specified $CD4^+$:$CD8^+$ ratio ($CD4^+$:$CD8^+$ RATIO) and CAR-antigen affinity (CAR AFFINITY) is added into the simulation. Data were collected every half day (720 time steps). To minimize variation as a result of the vasculature, all simulations use the same vasculature structure. Input files to create the simulations are shown in **Supplementary Table 9**.

#### 4.7 Data analysis

##### 4.7.1 Analysis of experimental data

Data from published studies of percent lysis at given antigen densities from various CARs, detailed in **Supplementary Table 4** and **Supplementary Data 1**, were estimated from plots included within each paper's results and processed for use in **Figure 2F** for the purpose of model validation. To ensure all the data could be viewed on plots with the same scaling, each percent lysis value was normalized to the maximum percent lysis for that CAR, and antigen values were normalized to the maximum value tested within a given dataset. The normalized values of the percent lysis or antigen values ($N_x$) plotted in **Figure 2F** were calculated as follows:

$$N_x = \frac{x}{X}$$

where:

- $x$ is the percent lysis or antigen value for a given data point within a dataset
- $X$ is the maximum percent lysis or antigen value within that given dataset

The error bars for both percent lysis and antigen value were also used, but they were normalized using the following error propagation formula using standard deviation for divided values (such as our normalized values) to calculate the error of the normalized percent lysis or normalized antigen value ($N_{\sigma_x}$):

$$N_{\sigma_x} = N_x \sqrt{\left(\frac{\sigma_x}{x}\right)^2 + \left(\frac{\Sigma_x}{X}\right)^2}$$

where

- $N_{\sigma_x}$ is the error of the normalized percent lysis or normalized antigen value
- $x$ is the percent lysis or antigen value for a given data point within a dataset
- $\sigma_x$ is the error for the percent lysis or antigen value for that given data point within a dataset
- $X$ is the maximum percent lysis or antigen value within that given data set
- $\Sigma_x$ is the error for the maximum percent lysis or maximum antigen value for that given data point within that dataset

When estimating data values from published plots, not all of the papers provided equivalent data, and some assumptions had to be made. Chmielewski et. al. recorded antigen levels in MFI and provided viability data rather than percent lysis (Chmielewski et al., 2004), so the data were converted to percent lysis ($P_{lysis}$) using the following formula:

$$P_{lysis} = 1 - P_{viability}$$

where $P_{viability}$ is the percent viability. Liu et. al. recorded antigen level in terms of ErbB2 RNA µg (Liu et al., 2015). Given that values were all normalized, this RNA quantity was used as a proxy for antigen expression level. Each of calculated value, assumptions, and notes associated with each published study are listed in **Supplementary Table 4** and **Supplementary Data 1**.

##### 4.7.2 Normalized cell counts

To compare outcomes between *in silico* experiments, we used normalized values of live cancer and healthy counts. Each normalized live cell metric ($N_P$) is calculated as follows:

$$N_P = \frac{n_F}{n_T}$$

where

- $n_F$ is the total number of live and cancer cell counts at the final time point (t = 7 d for `dish` and t = 30 d for `tissue`)

- $n_T$ is the total number of live cancer or healthy cells at the start of treatment timepoint (t = 0 d for `dish` and t = 21 for `tissue`)

To make a comparable metric to compare simulated monoculture `dish` outcomes to percent lysis values from experimental studies for use in **Figure 2F**, we calculated the percent lysis of simulated experiments ($P_{lysis}$) as follows:

$$P_{lysis} = 1 - N_C$$

where

- $N_C$ is the normalized live cancer cell metric

##### 4.7.3 Difference metric

To compare the outcomes of simulations with both cancer and healthy cells in a way that accounts for the tradeoff of cancer cell killing with healthy cell killing, we used a difference metric. The difference metric ($D$) is as follows:

$$D = N_H \left(\frac{c_\mathrm{T}}{h_\mathrm{T}}\right) - N_C$$

where

- $N_H$ is the normalized live healthy cell metric
- $N_C$ is the normalized live cancer cell metric
- $c_T$ is the number of live cancer cells at the start of treatment (t = 0 d for `dish` and t = 21 d for `tissue`)
- $h_T$ is the number of live healthy cells at the start of the treatment (t = 0 d for `dish` and t = 21 d for `tissue`)

The normalized live healthy cell metric is normalized by the ratio of initial live cancer to healthy cells at treatment start to account for differences in initial cell counts and ensure ratio components are equally weighted.
